# A Library of Induced Pluripotent Stem Cells from Clinically Well-Characterized, Diverse Healthy Human Individuals

**DOI:** 10.1101/2020.10.29.360909

**Authors:** Christoph Schaniel, Priyanka Dhanan, Bin Hu, Yuguang Xiong, Teeya Raghunandan, David M. Gonzalez, Rafael Dariolli, Sunita L. D’Souza, Arjun S. Yadaw, Jens Hansen, Gomathi Jayaraman, Bino Mathew, Moara Machado, Seth I. Berger, Joseph Tripodi, Vesna Najfeld, Jalaj Garg, Marc Miller, Colleen S. Lynch, Katherine C. Michelis, Neelima C. Tangirala, Himali Weerahandi, David C. Thomas, Kristin G. Beaumont, Robert Sebra, Milind Mahajan, Eric Schadt, Dusica Vidovic, Stephan C. Schürer, Joseph Goldfarb, Evren U. Azeloglu, Marc R. Birtwistle, Eric A. Sobie, Jason C. Kovacic, Nicole C. Dubois, Ravi Iyengar

## Abstract

A library of well-characterized human induced pluripotent stem cell (hiPSC) lines from clinically healthy human subjects could serve as a useful resource of normal controls for *in vitro* human development, disease modeling, genotype-phenotype association studies, and drug response evaluation. We report generation and extensive characterization of a gender-balanced, racially/ethnically diverse library of hiPSC lines from 40 clinically healthy human individuals who range in age from 22-61. The hiPSCs match the karyotype and short tandem repeat identity of their parental fibroblasts, and have a transcription profile characteristic of pluripotent stem cells. We provide whole genome sequencing data for one hiPSC clone from each individual, genomic ancestry determination, and analysis of Mendelian disease genes and risks. We document similar transcriptomic profiles, single-cell RNA-seq derived cell clusters and physiology of cardiomyocytes differentiated from multiple independent hiPSC lines. This extensive characterization makes this hiPSC library a valuable resource for many studies on human biology.

## Introduction

Since their groundbreaking discovery (Park et al., 2008; Takahashi et al., 2007; Yu et al., 2007), hiPSCs and cells differentiated from hiPSCs have become a powerful system to model *in vitro* human phenotypes, disease etiology and mechanisms, genotype-phenotype correlations and drug responses. However, such studies are often hampered by the small number of hiPSC lines or respective controls used for comparative analysis, reported somatic variability of derived hiPSCs, and heterogeneity of differentiated cells (Dubois et al., 2011; Fusaki et al., 2009; International Stem Cell et al., 2011; Nazor et al., 2012; Witty et al., 2014). In recent years, several groups and consortia reported on the establishment of various hiPSC libraries with a wide range in the number of hiPSC lines (Carcamo-Orive et al., 2017; Kilpinen et al., 2017; Panopoulos et al., 2017; Park et al., 2017; Streeter et al., 2017). These libraries included mostly disease/disorder-specific hiPSCs and control hiPSCs. The control hiPSC lines in these studies were derived from subjects without the specific diseases/disorders studied, either relatives or non-related individuals (Carcamo-Orive et al., 2017; Panopoulos et al., 2017), who were self-declared healthy subjects, or individuals with no medical disease history (Kilpinen et al., 2017; Panopoulos et al., 2017; Rouhani et al., 2014). However, in these studies a clinical health evaluation was not explicitly performed and no official documentation of health status reported. Nevertheless, such libraries have been used for the study of how genetic variants associated with complex genomic traits and phenotypes drive molecular and physiological variation in hiPSCs and their differentiated cells (Carcamo-Orive et al., 2017; D’Antonio-Chronowska et al., 2019; DeBoever et al., 2017; Karch et al., 2019; Kaserman et al., 2020; Kilpinen et al., 2017; Panopoulos et al., 2017; Park et al., 2017; Rouhani et al., 2014). Larger scale comparative and effective disease modeling, drug discovery and evaluation, and genotype-phenotype association studies suffer from the limited availability and inclusion of hiPSC lines from clinically screened, healthy individuals of various racial and ethnic backgrounds and age. Here, as part of the NIH-Common Fund Library of Integrated Network-Based Cellular Signatures (LINCS) program (Keenan et al., 2018), we report the creation of a hiPSC library from 40 selected individuals of diverse racial/ethnic backgrounds and ages who passed a rigorous clinical health evaluation. We provide the clinical characteristics of each of the participants from whom hiPSCs were derived, cytogenetics reports, short-tandem repeat (STR) authentication, and pluripotency analyses for all forty hiPSC lines, as well as whole genome sequencing data, genomic ancestry determination, and Mendelian disease gene and risk assessment.

Several studies have suggested a potential impact of donor cell source, cellular heterogeneity of established hiPSCs, as well as sex on cellular differentiation and physiological behavior (D’Antonio-Chronowska et al., 2019; Hu et al., 2016; Pianezzi et al., 2020; Sanchez-Freire et al., 2014). Variability in measured physiological parameters might also be affected by the lack of cellular homogeneity in differentiated hiPSCs. It remains unclear however, whether cells differentiated from multiple hiPSC clones from the same healthy subject will behave physiologically the same. Therefore, we studied the characteristics of ventricular and atrial cardiomyocytes differentiated from independent hiPSC clones from the same individual and from hiPSC clones from different individuals.

In summary, our diverse hiPSCs library from 40 clinically well-characterized healthy human individuals contributes a valuable resource to the scientific community for a broad variety of biomedical and pharmacological research.

## Results

### Recruitment, Health Evaluation, and Characterization of Individuals in the Mount Sinai Library of hiPSC Lines Derived from Diverse, Clinical Healthy Subjects

Potential healthy volunteers were approached across Mount Sinai through IRB-approved advertisements, and were pre-screened for potential inclusion in the study. Ninety-six male and female individuals who satisfied the initial pre-screening consented (Document S1 -*Blank Study-Consent Form* and Document S2 -*Blank HIV consent Form*) to the study and their sex, age, and race/ethnicity were recorded through an enrollment questionnaire (Figure 1A). Eighty-five underwent a formal and thorough evaluation by a screening study physician. Formal screening involved a full medical history, measurement of weight, height, waist and hip circumference, heart rate, blood pressure, respiratory rate, and oxygen saturation, a physical exam, and an electrocardiogram (EKG). Blood was drawn for analysis of clinically relevant parameters. A pregnancy test was included for female participants. All blood draws and pregnancy tests were sent to a certified clinical laboratory for analysis. The more than 20 exclusion criteria included abnormal EKG, family history of any cardiovascular disorder excluding hypertension in any first or second degree relative at age <50, family history of non-ischemic cardiomyopathy in any first or second degree relative at any age, prior organ transplantation, HIV positive status, history of myopathy, obesity, renal impairment, autoimmune disease, abnormal blood test results including brain natriuretic peptide, body mass index of ≥30 kg/m^2^, lifetime smoking of >2 pack-years or abnormality on physical exam. These data were entered onto a clinical report form (Document S3 -*Blank Clinical Report Form*) and a final assessment of study eligibility was then made by consensus of two senior study physicians (J.C.K., D.T.) and the screening study physician. If eligibility was approved by all three physicians, the subject then underwent skin biopsy and formally became one of the included study subjects. Of the 85 subjects who were screened, 42 (48.3%) were deemed eligible for final inclusion. Reasons for a screened subject’s exclusion from further participation are summarized in Figure 1B. Of the 42 eligible individuals, a skin biopsy was successfully scheduled and performed on 40. Table S1 lists the clinical parameters of the forty healthy subjects. These 40 healthy male (22) and female (18) individuals ranged in age from 22-61. These observations demonstrate that an inclusive health evaluation and strict selection criteria for what constitutes a healthy individual will produce more stringent definition of health status than simply relying on a participant’s self-declaration or medical history.

**Figure 1.**
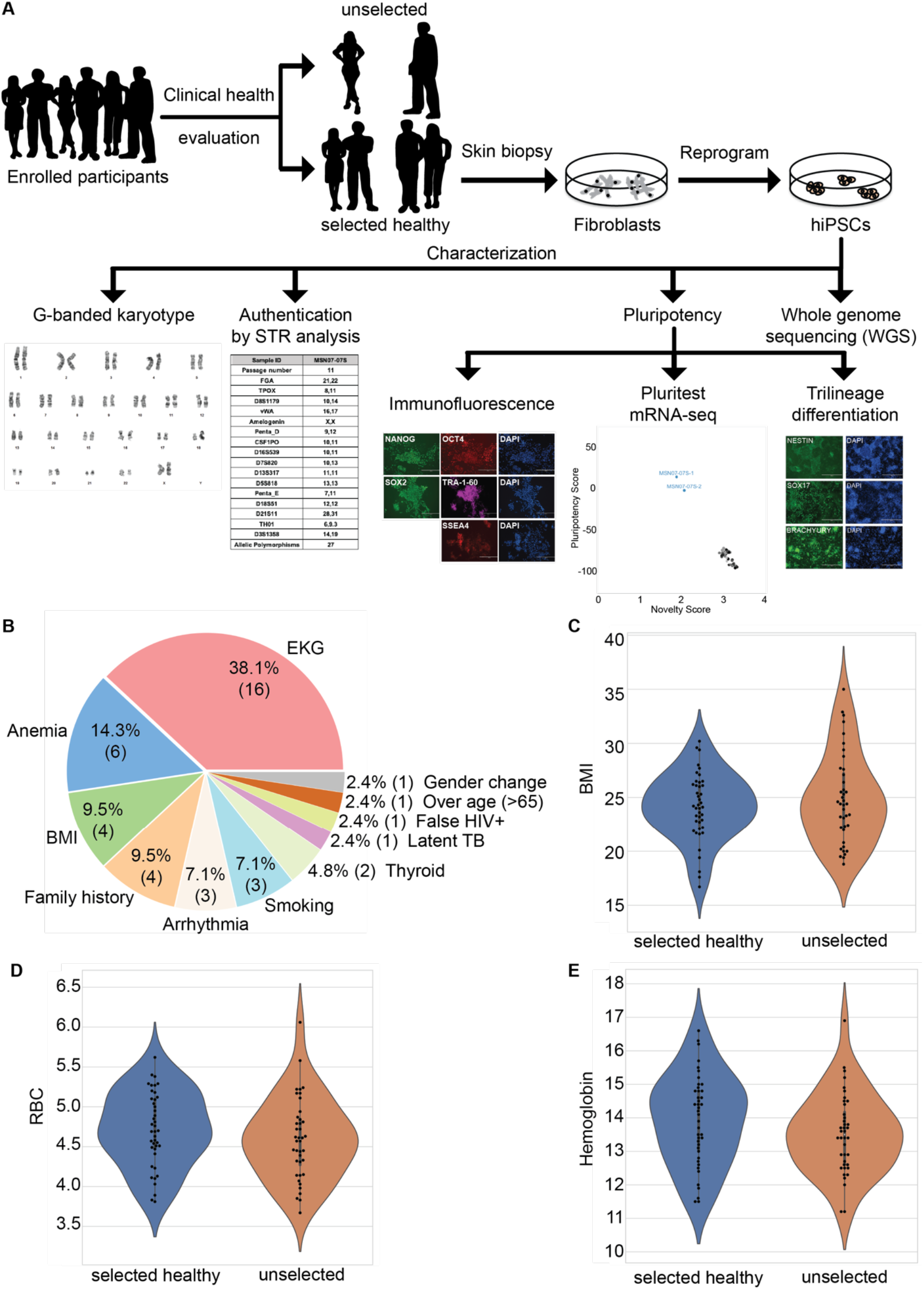
Description of subject selection for the Mount Sinai clinically healthy hiPSC library. (A) Flow chart for subject selection and hiPSC generation and characterization. See Table S1 for the summary characterizations performed for all 40 selected, clinically healthy subjects and derived hiPSCs. (B) Pie diagram of reasons for excluding screened participants from the final group of eligible subjects. The percentage as well as the number of participants (in parenthesis) excluded for each specific main reason is presented. (C-E) Violin plots of clinical selection criteria of body mass index (BMI, kg/m^2^), red blood cell counts (RBC; × 10^6^/uL) and hemoglobin level (g/dL) of healthy/selected and excluded participants. See Document S3 for the clinical report form and Table S1 for the clinical characteristics of all clinically healthy/selected individuals.

### Generation, Authentication and Characterization of hiPSC Clones

We derived fibroblast lines from skin biopsy samples taken from the 40 eligible healthy subjects. We used integration-free reprogramming methods (mRNA with microRNA boost (R) (Warren et al., 2010)) and/or Sendai virus (S) (Fusaki et al., 2009)) to generate hiPSCs and establish multiple clones from all 40 fibroblast lines. Our present resource consists of one hiPSC clone per individual plus 2 additional clones each for two hiPSC lines (Table S2). All data, as well as linked associated metadata, can also be found on the searchable LINCS data portal (Koleti et al., 2018; Stathias et al., 2020). All hiPSCs in our library match the karyotype of the parental fibroblast line. In all cases except one the karyotype was normal (Figure 2A, Table S3). The abnormal karyotype, which was a t(1;17)(p34;q23) translocation, was observed for MSN24. In total, we analyzed the karyotypes of 5 independent hiPSC clones derived from individual MSN24. They all carried the t(1;17)(p34;q23) translocation. It is not uncommon that a karyotype is abnormal (International Stem Cell et al., 2011; Mayshar et al., 2010; Peterson and Loring, 2014; Taapken et al., 2011); however, a frequency of 100% abnormal hiPSC clones is very unusual. Thus, we karyotyped the parental fibroblasts. Interestingly, the parental fibroblasts already carried this chromosomal abnormality. Hence, this may represent a case of a healthy individual, who is the carrier of a balanced translocation. No report exists about this specific translocation. Of 84 hiPSC clones we karyotyped, 26 (in addition to the 5 analyzed MSN24 hiPSC clones) showed an abnormal karyotype (31%). This is consistent with published reports of abnormal karyotypes in up to one third of derived human embryonic stem cells (ESCs) and hiPSCs (International Stem Cell et al., 2011; Mayshar et al., 2010; Peterson and Loring, 2014; Taapken et al., 2011). All hiPSC clones in our library are negative for SeV (Tables S2 and S4) and have been authenticated by STR analysis to match the profile of their respective parental fibroblast line (STR Table available at dbGAP under accession phs002088.v2.p1). This is an important and necessary quality control measure to ensure the origin and authenticity of the derived hiPSC clones.

**Figure 2.**
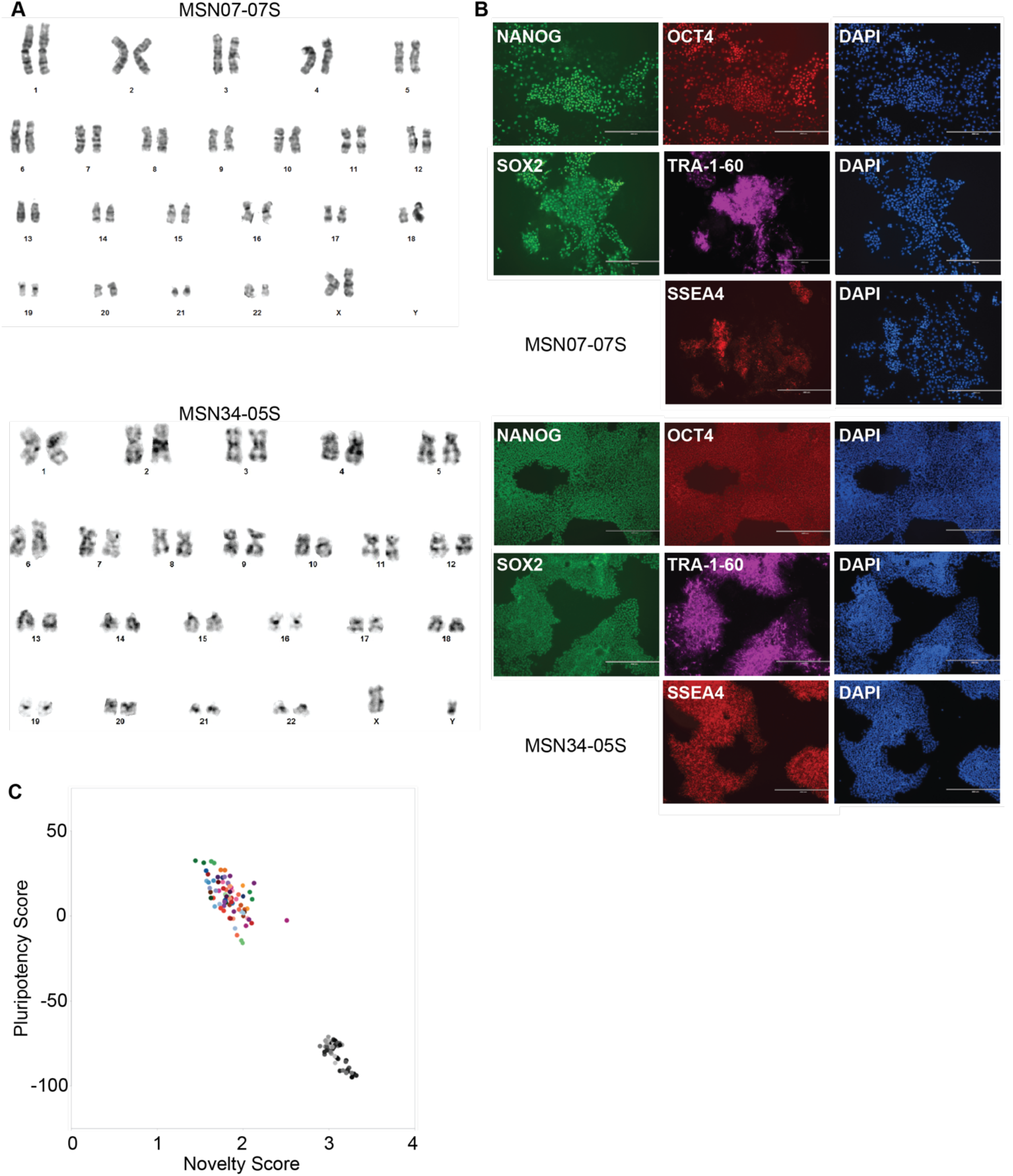
Characterization of two representative hiPSC lines/clones. (A) G-banded karyotypes for female hiPSC clone MSN07-07S and male hiPSC clone MS34-05S. See Table S3 for the cytogenetics of all 40 fibroblast lines and one associated derived hiPSC clone. (B) Immunocytochemistry of pluripotency markers, NANOG, OCT4, SOX2, TRA-1-60 and SSEA4 in representative hiPSC clones, MSN07-07S and MS34-05S. DAPI is used to stain nuclei. Bar, 400 μm. See Figure S1 for the immunocytochemistry of one representative hiPSC clone derived from each of the 40 fibroblast lines. (C) PluriTest summary plot of RNA-seq based transcriptomic analyses performed in duplicate for one hiPSC clone per clinically healthy individual (colored circles). As a reference, the PluriTest results of transcriptomic data of fibroblasts from 66 individuals (Hagai et al., 2018) are plotted in black/grey. See Figure S2 for the PluriTest plots of one representative hiPSC clone derived from each of the 40 fibroblast lines.

One karyotypically normal (with the exception of MSN24) and authenticated hiPSC clone generated from each of the forty clinically healthy individuals was further characterized by immunocytochemistry for expression of the pluripotency-associated markers NANOG, OCT4, SOX2, TRA-1-60 and SSEA4 (Figure 2B, Figure S1). We also conducted RNA-seq (Table S4) based PluriTest assay (Panopoulos et al., 2017) as an additional measure of the pluripotent status of all hiPSC clones (Figure 2C, Figure S2, Table S4). We included the pluripotency and novelty scores retrieved from fibroblasts of 66 individuals as comparative reference (Hagai et al., 2018). We next compared the gene expression profiles of the 40 hiPSC lines (in duplicates) with 77 randomly-chosen hiPSC lines described by Kilpinen and colleagues (Kilpinen et al., 2017)) as well as 66 human fibroblast lines (Hagai et al., 2018), from which Kilpinen and colleagues derived hiPSC lines from. Pairwise Pearson correlation and principal component analysis (PCA) demonstrate that the two sets of hiPSC lines are similar to each other and very distinct from the profiles of the fibroblast lines (Figure 3, Table S5A-C). Lastly, we assessed pluripotency of a selection of hiPSC lines/clones by *in vitro* tri-lineage differentiation (Figure S3). All assayed hiPSC lines/clones differentiated efficiently to cells representing (neuro)ectoderm (NESTIN^+^), endoderm (SOX17^+^) and mesoderm (BRACHYURY^+^) demonstrating their pluripotency.

**Figure 3.**
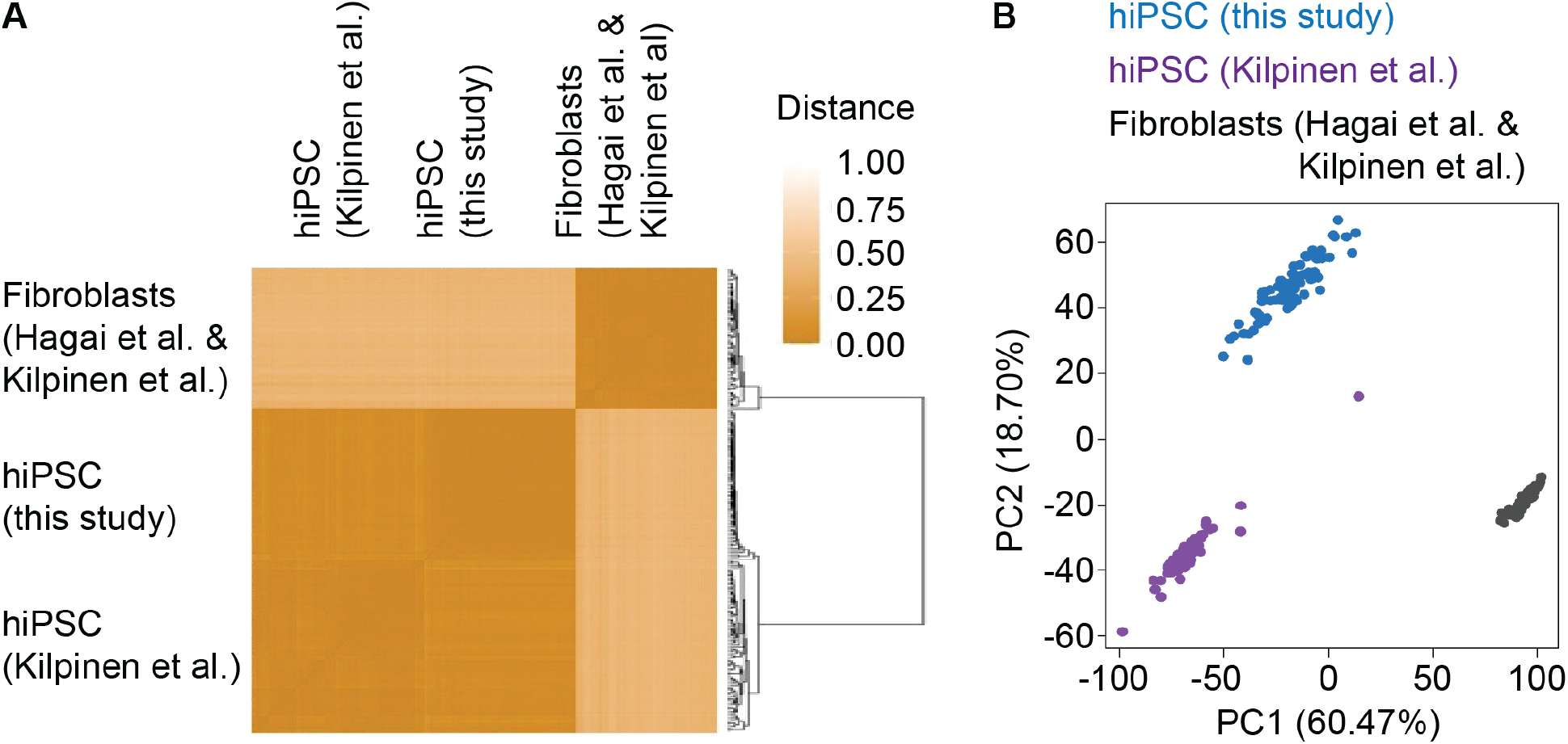
Comparative hiPSC transcriptome analysis. (A) Pairwise Pearson correlation was determined between RNA-seq based gene expression profiles of the 40 hiPSC samples (in duplicates) from this study with 77 randomly selected hiPSC samples (Kilpinen et al., 2017) and 66 human fibroblast samples (Hagai et al., 2018). Hierarchical clustering documents close correlation of the two hiPSC sample sets with clear separation from the fibroblast lines. (B) Principal component analysis of all three combined sets identifies principal component (PC) 1 with ~60% of the total variance to not significantly distinguish between the two hiPSC sets but to clearly separate them from the fibroblast lines.

We performed genetic ancestry determination using the whole-genome sequencing information from all forty hiPSC lines (Table S6). This analysis provided additional confirmation of the authenticity of the hiPSC lines. It additionally reflected the diverse nature of the subjects’ racial and ethnic backgrounds. The selected individuals have origins in East Africa, West Africa, East Asia, the Central India subcontinent, the Southern India subcontinent, Eastern Mediterranean, Northeast Europe, Northern and Central Europe, Southwest Europe, and the Anatolia/Caucasus/Iranian Plateau region.

Our data support our conclusion that we established a gender-balanced, racially/ethnically diverse library of well-characterized hiPSCs from forty clinically healthy individuals who range in age from 22-61.

### Clinically Healthy Individuals Can Be Carriers of Genetic Variants with Disease Risks

Although all forty individuals from whom we established the hiPSC library were determined to be clinically healthy, we next assessed whether they carry potential genetic disease risks. To assess this, we interrogated the whole-genome sequencing information by performing annotation of gene variants associated with both dominant and recessive disorders (variant call format (vcf) files are provided along with the whole genome sequencing data in dbGAP under accession phs002088.v2.p1). All variants called in the samples with non-conflicting interpretations of at least likely-pathogenic in Clinvar (Landrum et al., 2014) with defined assertion criteria were annotated using Annovar. For nine individuals (22.5%) zero pathogenic or likely pathogenic variants were detected. Two subjects (5%) carried one likely pathogenic variant, seven individuals (17.5%) carried one to two pathogenic variants, three subjects (7.5%) carried one pathogenic/likely pathogenic variant, and nineteen of the selected subjects (47.5%) were carriers of a combination of pathogenic and/or likely pathogenic and/or pathogenic/likely pathogenic variants (Table S7). The maximum of pathogenic and/or likely pathogenic variants per subjects detected were six. Variants were reviewed by a Board Certified Medical geneticist (S.I.B.). The variants associated with dominant diseases are noted in Figure 4A. These variants are consistent with the history of coming from a healthy adult as they can result in mild common phenotypes (such as eczema) or disorders with later onset and incomplete penetrance (cancer risks and late onset cardiomyopathies). These findings highlight that clinically well-characterized healthy human individuals can harbor genetic variants with disease risks, a fact that may not be surprising but should to be considered when using control hiPSCs, including those from our library, as controls in specific disease modeling research and drug evaluation/toxicity studies where certain variants might impact the results and their interpretation.

**Figure 4.**
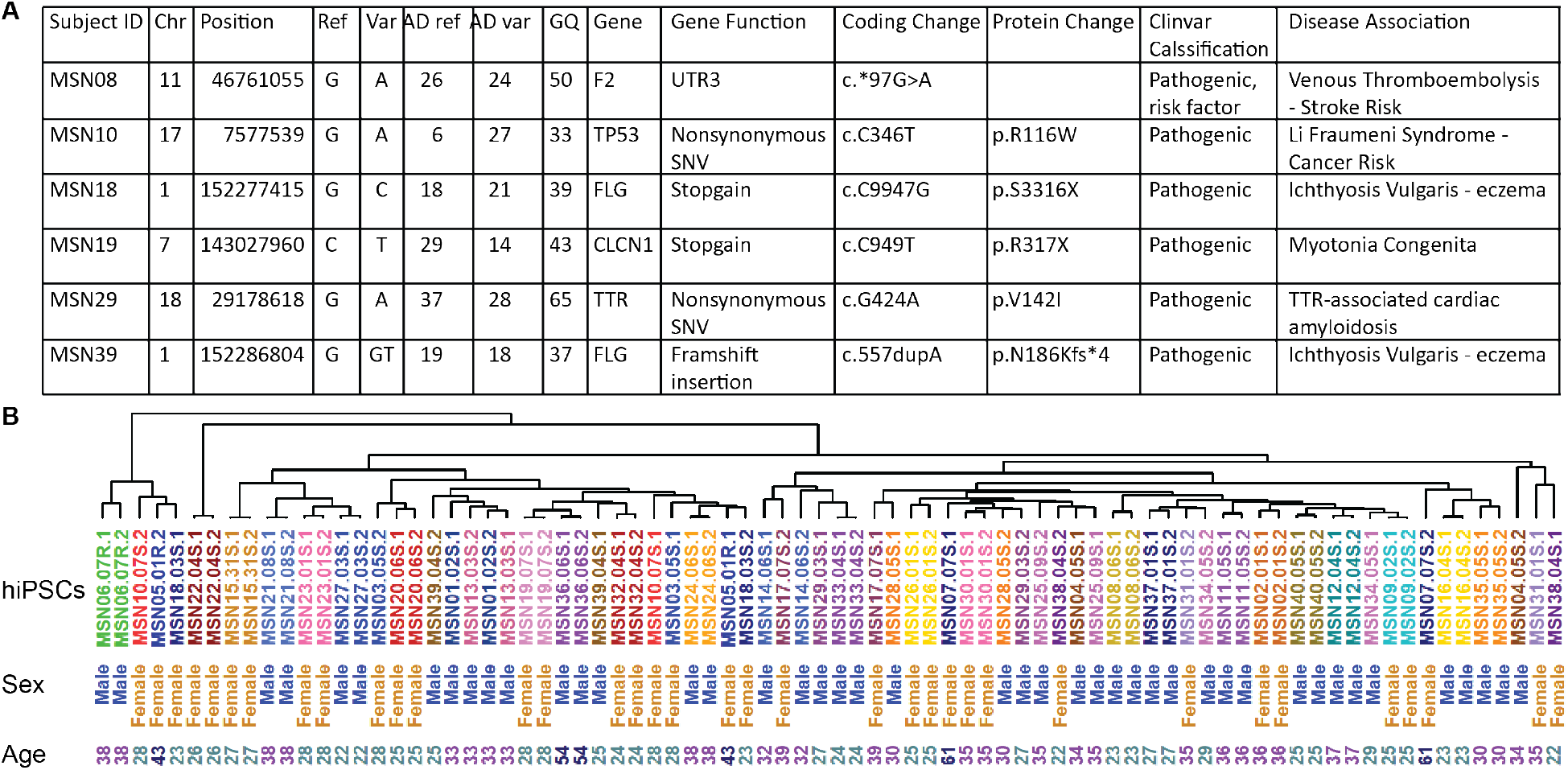
Genetic variants with disease risks and global gene expression independence of age and sex. (A) Subjects with genetic disease variants associated with a dominant presentation. (B) Pairwise correlation was determined between RNA-seq raw read counts of duplicate samples of all forty hiPSC lines, followed by hierarchical clustering. The resulting dendrogram documents high similarity between duplicate samples and hiPSC lines. No influence of age or sex was found on gene expression.

### hiPSC Gene Signature is Independent of Age or Sex of Individuals from whom hiPSCs are Derived

To assess whether age or sex of the included individuals could segregate the derived hiPSC lines based on global gene expression, we determined pairwise correlations between bulk mRNA-seq raw read counts of duplicate samples of all forty hiPSC lines. This was followed by hierarchical clustering. The resulting dendrogram documents a high similarity both between duplicate samples and across the forty hiPSC lines (Figure 4B). Similar to pluripotency marker expression (Figure 2B, Figure S1) and PluriTest scores (Figure 2C, Figure S2), we observed no significant influence of age or sex on global gene expression. These results indicate that global gene expression, within the wide age range we studied, is independent of age and sex of clinically well-characterized healthy human subjects from whom hiPSC lines are derived.

### Similar functional characteristics of cardiomyocytes derived from multiple hiPSC lines

A recent report determined that that sex affects cardiac ventricular and atrial differentiation outcomes whereas inherited genetic variation does not (D’Antonio-Chronowska et al., 2019). It remains unknown, however, whether hiPSCs derived from different healthy individuals will exhibit comparable molecular and physiological characteristics when differentiated using similar protocols. We sought to answer this question by performing both molecular and physiological analysis on cardiomyocytes. For this, we differentiated several age-matched, male and female hiPSC lines into cardiac mesoderm (KDR^+^/PDGFRA^+^) and ventricular cardiomyocytes (SIRPA^+^/CD90^−^)(Dubois et al., 2011; Kattman et al., 2011) followed by purification using metabolic selection (Tohyama et al., 2013). We measured calcium transients and compared several features between the different hiPSC lines (Figure S4A, Table S8A). We also conducted bulk and single cell (sc) RNA-seq experiments to interrogate global transcriptomes at the population and single cell level (Figure S4B-C). High-level comparisons of both the physiological and scRNA-seq data failed to reveal any obvious observable differences between cardiomyocytes differentiated from the various hiPSC lines (Figure S4A-C).

For a more in-depth comparison, we performed additional experiments in cardiomyocytes differentiated from hiPSCs from two individuals, in order to examine multiple hiPSC clones from the same individual. For this, we differentiated three independent, karyotypically normal hiPSC clones derived from two age-matched males into atrial and ventricular cardiac mesoderm (KDR^+^/PDGFRA^+^) and cardiomyocytes (SIRPA^+^/CD90^−^)(Devalla et al., 2015; Dubois et al., 2011; Kattman et al., 2011; Lee et al., 2017; Zhang et al., 2011) (Figure 5). As expected, we observed some variability in the differentiation efficiency between both experimental replicates (separate differentiations) and between the two hiPSC lines, both for cardiac mesoderm and differentiated cardiomyocytes (Figure 5). Similar variability was present in both the atrial and ventricular differentiations. We again measured intracellular calcium waveforms in preparations purified by metabolic selection (Tohyama et al., 2013) (Figure 6, Table S8B). As expected, atrial preparations tended to exhibit shorter calcium waveforms and faster beating rates than ventricular preparations (Figure 6A). We quantified various metrics from these waveforms (see Supplemental Experimental Procedures) and performed PCA on the collected results (Figure 6B). These results suggest that clear differences in physiological metrics can be seen between atrial and ventricular preparations, but not between cells from the two donors, or between the different clones tested. An examination of selected metrics (beating rate and calcium transient duration) is consistent with this impression, as significant differences are observed between atrial and ventricular cells, but are difficult to detect between donors or between clones. (Figure 6C–E). Overall, these results suggest that cardiomyocytes differentiated from hiPSCs originally derived from different clinically healthy individuals, or independent clones from the same individual exhibit similar characteristics, at least when differentiated under well controlled conditions.

**Figure 5.**
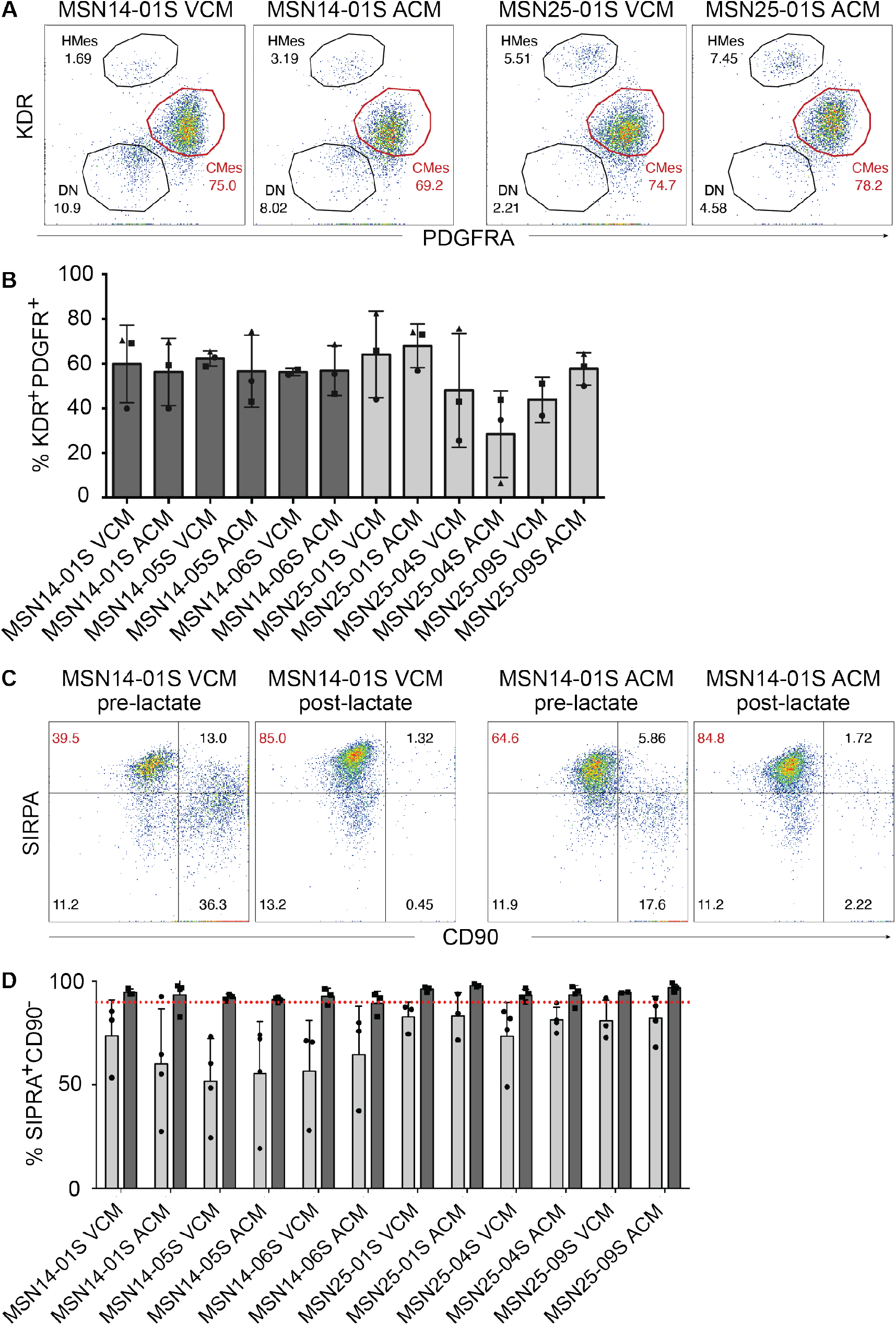
Atrial and ventricular differentiation of hiPSCs. (A) Representative flow cytometry plots of KDR and PDGFR expression on cells at day 5 of atrial (ACM) and ventricular (VCM) cardiomyocyte differentiation from two racially/ethnically distinct, age-matched male clinically healthy subjects (MSN14 and MSN25, respectively) (top). (B) Percentage of cells expressing both KDR and PDGFR across three independent hiPSC clones each established from subjects MSN14 (dark grey) and MSN25 (light grey). Symbols (circles, squares and triangles) represent independent differentiations. Data are represented as the mean±SD. (C) Representative flow cytometry plots of SIRPA and CD90 expression at day 20 of atrial (ACM, right) and ventricular (VCM, left) differentiation from hiPSC clone MSN14-01 pre-lactate and post-lactate selection. (D) Percentage of cells expressing SIRPA but not CD90 across the three independent hiPSC clones established from the same two racially/ethnically distinct, age-matched male clinically healthy subjects as in (A) pre-lactate (light grey) and post-lactate (dark grey) selection. Individual dots represent independent differentiations. Data is represented as the mean±SD. The red dotted line indicates 95% purity, which was the cut off set for a differentiated line to be used for the subsequent physiology assays.

**Figure 6.**
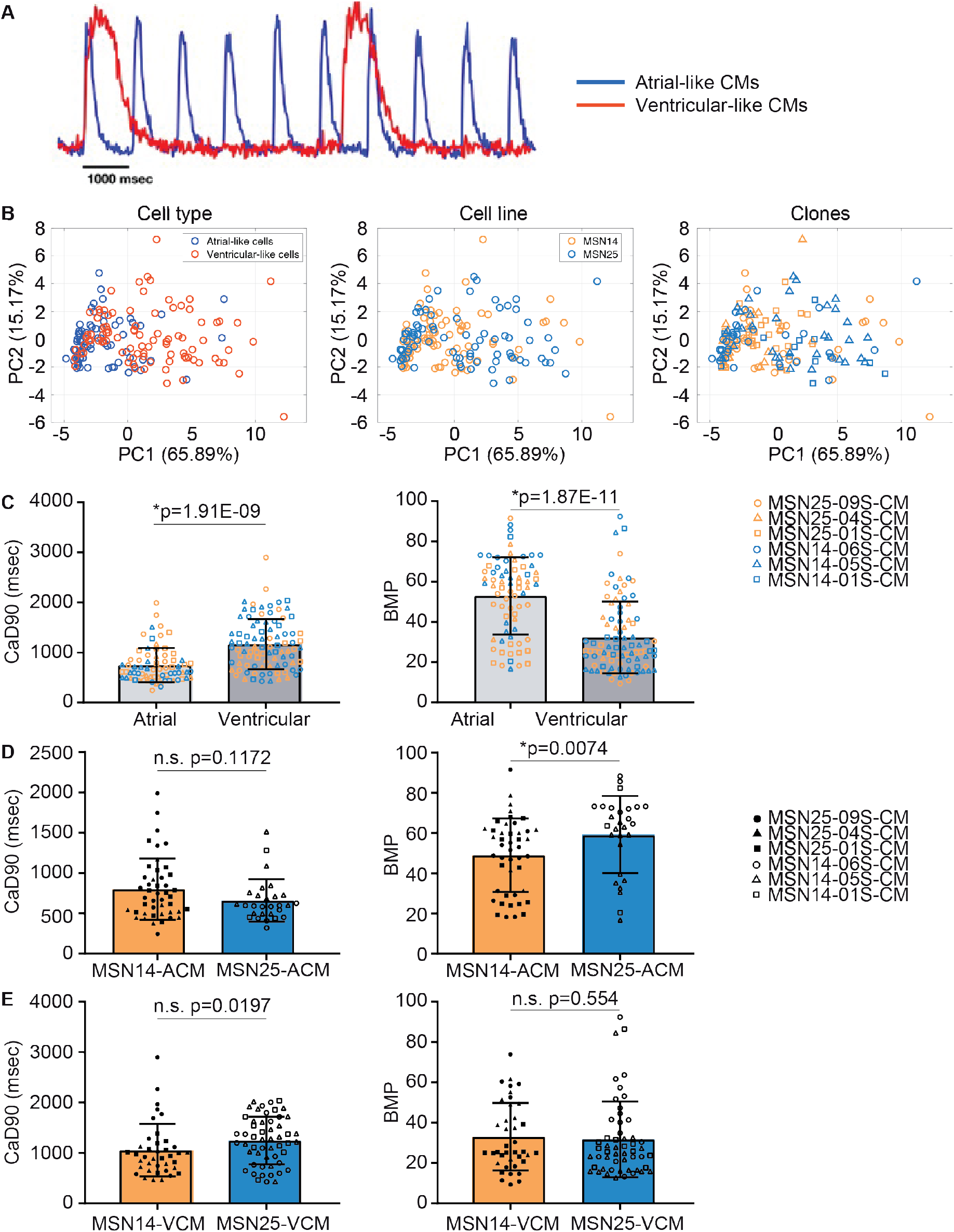
Calcium transient analysis of hiPSC-derived cardiomyocytes from two racially/ethnically distinct, age-matched males. (A) Representative calcium transient analysis traces of ventricular (red trace) and atrial (blue trace) hiPSC-derived cardiomyocytes. hiPSC-CMs were lactate-purified at day 20 of differentiation and analyzed at day 30 of differentiation. (B) Principal component analysis of multiple features measured from calcium transients recorded from atrial (blue) and ventricular (red) cell types differentiated from independent hiPSC clones established from each of two racially/ethnically distinct, age-matched male clinically healthy subjects (MSN14 - orange; MSN25 – blue) at day 30 of differentiation after lactate selection at day 20. The data are accrued from 2-3 independent differentiations for each hiPSC line and clone. (C) Comparison of calcium transient duration (left) and beat frequency (right) between VCM (light gray) and ACM (dark gray) differentiated from three hiPSC clones derived from two patients (MSN14, orange; MSN25, blue). The data are accrued from 2-3 independent differentiations. *, p<0.05 from unpaired Mann-Whitney test. (D) Comparison of calcium transient duration and frequency between two patient lines (MSN14 and MSN25) for atrial cells alone. The data are accrued from 2-3 independent differentiations for each hiPSC line/clone. *, p≤0.01 from unpaired Mann-Whitney test was considered significant; n.s. (p>0.01), non-significant. (E) Comparison of calcium transient duration and frequency between two patient lines from ventricular cells alone. The data are accrued from 2-3 independent differentiations for each hiPSC line/clone. *, p<0.05 from unpaired Mann-Whitney test; n.s. = non-significant.

## Discussion

Previous efforts to create libraries of hiPSCs have focused on diseases and associated controls that included disease-unaffected relatives or non-related individuals, self-declared healthy subjects, or persons with no medical disease record (Carcamo-Orive et al., 2017; Kilpinen et al., 2017; Panopoulos et al., 2017; Park et al., 2017; Streeter et al., 2017). To our knowledge, no gender-balanced, racially/ethnically diverse hiPSC library of well-characterized clinically screened healthy individuals exists. We have created such an hiPSC library consisting of well-characterized hiPSC lines that were established using integration-free reprogramming methods from forty healthy male and female individuals of diverse racial/ethnic backgrounds, who passed a rigorous health evaluation. As was done for other studies that generated “control” hiPSCs (Carcamo-Orive et al., 2017; Kilpinen et al., 2017; Panopoulos et al., 2017; Park et al., 2017; Streeter et al., 2017), our subjects had to complete a health questionnaire and had their medical histories evaluated. However, and in contrast to these other studies, the individuals who were selected for a skin biopsy in our study underwent a detailed health screen conducted by an internal medicine physician, and their results were evaluated by a three-member clinical panel. This screen included the measurement of general parameters, analysis of clinically relevant parameters present in the blood, a physical examination of the respiratory and gastrointestinal/abdominal systems, a neurological exam and a cardiac exam, including an EKG.

While all our hiPSC lines were distinct form human fibroblast lines in the RNA-seq based PluriTest assay (Figures 2D and S2), pluripotency and novelty scores of certain hiPSC lines and even of certain duplicate samples (Table S4) were inexplicably different and beyond the threshold set by Panopoulos and colleagues, who first applied the RNA-seq based PluriTest (Panopoulos et al., 2017). However, comparative whole transcriptome analysis of the 40 hiPSC lines (in duplicates) with 77 randomly-chosen hiPSC lines (Kilpinen et al., 2017)) and 66 human fibroblast lines (Hagai et al., 2018), and PCA clearly showed that the two sets of hiPSC lines are similar to each other and very distinct from the profiles of the fibroblast lines (Figure 3). This finding together with expression of 5 pluripotency markers for all hiPSC lines and clones (Figures 2 and S1) and demonstrated *in vitro* tri-lineage differentiation potential of a selection of lines/clones (Figure S3) provide confidence in the pluripotency of our hiPSC library.

It is well established that race and ethnicity, which are used as a way of categorizing people of shared ancestry and physical traits (Sankar et al., 2004), as well as sex (Soldin and Mattison, 2009) are contributing factors to interindividual differences in drug exposure and/or response (Ramamoorthy et al., 2015; Wilson et al., 2001). These are important factors to consider during drug development and evaluation of drug responses as they imply diverse risk-benefit ratios in specific populations. Therefore, any established hiPSC library, if it is not a library of hiPSC lines derived from individuals with a disorder that is preferentially associated with a particular racial/ethnic group, should be gender-balanced and racially/ethnically diverse. Our hiPSC library with 18 females and 22 males fulfills these criteria as it contains lines from individuals that self-identify as belonging to a wide range of ethnicities originating in multiple continents.

While our hiPSC library was derived from well-characterized clinically healthy individuals, whole-genome sequencing analysis revealed that several individuals were carriers of genetic variants associated with recessive and dominant pathogenic disease risk. This is important and useful information, particularly when using the hiPSC lines as controls for modeling a particular disease/disorder, or for evaluating the response to drugs. Several expression quantitative trait loci (eQTL) studies in hiPSCs have shown that genetic variations account for many of gene expression differences between hiPSC lines (Carcamo-Orive et al., 2017; DeBoever et al., 2017; Kilpinen et al., 2017; Panopoulos et al., 2017; Rouhani et al., 2014). While we did not investigate eQTL on global gene expression in hiPSCs, we found that age and sex had no impact (Figure 4B). This finding is in agreement with Kilpinen and colleagues’ study, which showed that neither sex nor cell passage number substantially influenced gene and protein expression in hiPSCs (Kilpinen et al., 2017).

If hiPSCs from healthy individuals are used as a control group for the study of a particular disease, then ideally the molecular and physiological functions of these differentiated cells should be similar. We tested this possibility using multiple age-matched, male and female hiPSC lines differentiated into cardiomyocytes. Physiological function and molecular profiles were assessed using calcium transient measurements, and RNA-seq (both bulk and single cell), respectively. Although small differences between groups could sometimes be seen when dozens of samples from each hiPSC line were assessed (e.g., Figure S4), which might be attributed to differences in proportions of cell sub-populations, there was always substantial overlap between the hiPSC lines in all the measurements we performed.

Further detailed analyses of three independent clones each from two separate hiPSC lines, under conditions where the atrial and ventricular cardiomyocytes showed clearly observable differences, did not reveal significant differences between either the independent clones or between the two hiPSC lines. (Figures 4 and 5). Although the general applicability of this finding needs to be confirmed, this may have practical and financial implications, especially for large-scale studies, where use of a single hiPSC clone per subject rather than multiple clones would curtail costs and reduce potential technical variability. Although our observations suggest that there are unlikely to be large differences between multiple hiPSC lines from healthy individuals, drawing firm generalized conclusions will require studies with tens or hundreds of independent hiPSC lines conducted under standardized conditions. Additionally, these conclusions from cardiomyocytes may not be applicable for these hiPSCs differentiated into other cell types or different organoids.

The possibility of differences between lines should be considered when planning to test drug responses across cell types differentiated from hiPSC derived from a range of racially and ethnically diverse, clinical well-characterized healthy male and female individuals. One could potentially expect different responses to the drugs investigated based on racial and ethnic background as well as genetic variants (pharmacogenomics). In this context, it is noteworthy that a recent report studying 10 control human pluripotent stem cell lines for drug screening in an engineered heart tissue format found that while there was great variability in baseline contractile parameters observed across the different lines, the variability appeared less relevant for drug screening (Mannhardt et al., 2020). A number of constraints precluded us from testing every healthy hiPSC line in our library for each of the conditions and assays described here. Hence, further studies will be required to unequivocally establish all of the conclusions of this study. These limitations notwithstanding, we provide a gender-balanced, racially/ethnically diverse library of hiPSCs from 40 clinically well-characterized healthy human individuals ranging in age from 22-61 years. This library is accompanied with comprehensive quality control characterization of karyotype, STR-matching to each individual (authentication), pluripotency analysis, whole-genome sequencing, ancestry determination, and disease gene and risk analysis. This hiPSC library can be useful to investigators looking for appropriate control hiPSCs to study normal human development, investigate drug responses/evaluation or as controls for their specific disease model.

### Experimental procedures

#### Recruitment and health screening of subjects

Female and male subjects of diverse racial and ethnic background (as per NIH standards described in NOT-OD-15-089) were recruited and consented under Mount Sinai Institutional Review Board approved protocol (HS# 14-00530) (Documents S1 and S2). Consented subjects were evaluated for their health status at the Mount Sinai Clinical Research Unit by internal medicine physicians. The evaluation involved completion of a health questionnaire, clinical history, and measurement of weight, height, waist and hip circumference, heart rate, blood pressure, respiratory rate, and oxygen saturation. Blood was drawn for analysis of clinically relevant parameters (note that determination of glucose levels was not limited to a specific state (fasting/non-fasting) or time of sampling), and the subjects underwent respiratory, gastrointestinal/abdominal, neurological and cardiovascular exams including an EKG (Document S3). A pregnancy test was included for female participants. Subjects with no clinical history of disease, normal physical exam, normal EKG and evaluated parameters within the normal range (Document S3) were classified as healthy by a medical panel consisting of two internal medicine physicians and an interventional cardiologist. Clinical health information remained within Mount Sinai’s protected electronic medical records system.

Clinical data of all consented subjects were extracted from the individual’s electronic medical record through the Electronic Privacy Information Center server of the Mount Sinai Data Warehouse. Clinical data for each participant were downloaded manually. Subjects’ age, sex, race/ethnicity (as per NIH standards) and selected healthy/passed or unselected/unpassed status were transcribed to a database and healthy subjects were de-identified with a unique sample ID (MSNxx, where xx is a two-digit number ranging from 01 to 40) (Tables S1 and S2). The primary reason for exclusion of forty-two individuals from the final pool of eligible healthy subjects was plotted as a pie-chart (Figure 1B–E). To visualize the distribution of eligible healthy subjects vs. excluded subjects, we plotted violin swarm plots of (i) body mass index (BMI) with median value for eligible healthy subjects of 24.15 vs. excluded of 24.4 kg/m^2^ (ii) Hemoglobin with median value for healthy of 14.5 vs excluded of 13.5 g/dL, (iii) Red Blood Cells with median value for healthy of 4.7 vs excluded of 4.6×10^6^/uL by using Seaborn python data visualization library. All analyses were done using Python programing.

#### Biopsy punch, derivation and culture of fibroblasts

Our detailed SOP can be found in Supplementary Experimental Procedures. Briefly, a skin sample was taken from each of 40 clinically healthy subjects using a 3 mm sterile disposable biopsy punch (Integra Miltex, Cat# 98PUN3-11) during a second visit at the Clinical Research Unit. Each sample was cut into smaller pieces and placed into gelatin-coated tissue culture dishes with DMEM supplemented with 20% FBS, Penicillin/Streptomycin, non-essential amino, acids, 2mM L-glutamine, 2 mM sodium pyruvate (all from Thermo Fisher Scientific) and 100 μM 2-mercaptoethanol (MP Biomedicals, Cat# 194705) to establish fibroblast lines. Fibroblasts were harvested using TrypLE Express (Thermo Fisher Scientific, Cat# 12605010) and passaged at a 1 to 4 split ratio. Fibroblasts were cryopreserved in 40% DMEM, 50% FBS and 10% DMSO (Millipore Sigma, Cat# D2438).

#### Reprogramming fibroblasts to hiPSCs

Mycoplasma-free fibroblasts at passage number 3-5 were reprogrammed using the mRNA reprogramming Kit (Stemgent, Cat # 00-0071) in combination with the microRNA booster kit (Stemgent, Cat# 00-0073) and/or the CytoTune™-iPS 2.0 Sendai Reprogramming Kit (Thermo Fisher Scientific, Cat# A16517) according to our SOPs that can be found in Supplementary Experimental Procedures. hiPSC clones will be deposited with, and made available through, WiCell.

#### Human iPSC culture

Established hiPSCs were cultured on plates coated with Matrigel (Corning, Cat# 254248) in mTeSR medium (STEMCELL Technologies, Cat# 05850) in a humidified incubator at 5% CO2, 37°C. Cells were passaged using ReLeSR (STEMCELL Technologies, Cat# 05872) according to the manufacturer’s instructions and grown for 16-24 hrs in mTeSR medium supplemented with 2 μM Thiazovivin (Millipore Sigma, Cat# 420220).

#### Human iPSC-cardiomyocyte (CM) differentiation

Human iPSCs were maintained in E8 media and passaged every 4 days onto Matrigel-coated plates (Roche) before differentiation. On Day 0 (start of differentiation) iPSCs were treated with 1mg/mL Collagenase B (Roche, Cat# 11088807001) for one hour, or until cells dissociated from plates, to generate embryoid bodies (EBs). Cells were collected and centrifuged at 300 rcf for 3 min, and resuspended as small clusters of 50–100 cells by gentle pipetting in CM differentiation media composed of RPMI 1640 (Thermo Fisher Scientific, Cat# 11875085) containing 2 mM/L L-glutamine (Thermo Fisher Scientific, Cat# 25030149), 4×10^−4^ M monothioglycerol (Millipore Sigma, Cat# M6145), and 50 μg/mL ascorbic acid (Millipore Sigma, Cat# A4403). Differentiation media was supplemented with 3ng/ml BMP4 (Biotechne-R&D Systems) and 3 μM Thiazovivin (Millipore Sigma, Cat# 420220) and EBs were cultured in 6-cm dishes (USA Scientific, Cat# 8609-0160) at 37°C in 5% CO_2_, 5% O2, and 90% N2. On Day 1, the media was changed to differentiation media supplemented with 20 ng/mL BMP4 (Biotechne-R&D Systems) and 20 ng/mL Activin A (Biotechne-R&D Systems), 5ng/mL bFGF (Biotechne-R&D Systems) and 1 μM Thiazovivin (Millipore Sigma, Cat# 420220). On Day 3, EBs were harvested and washed once with DMEM (Gibco). Media was changed to differentiation media supplemented with 5 ng/ml VEGF (Biotechne-R&D Systems) and 5 μM/L XAV939 (Reprocell-Stemgent, Cat# 04-0046). To generate atrial CMs, retinoic acid (RA) was added to the differentiation media at 0.5 μM (Devalla et al., 2015; Lee et al., 2017). On Day 5, media was changed to CM differentiation media supplemented with 5 ng/mL VEGF (Biotechne-R&D Systems). After Day 8, media was changed every 3-4 days to differentiation media without supplements (Dubois et al., 2011).

#### Lactate metabolic selection

EBs were dissociated on day 20 of CM differentiation and replated on Matrigel-coated 6 well plates at 1×10^6^ cells/well in CM differentiation media supplemented with 1 μM Thiazovivin (Millipore Sigma, Cat# 420220). Media was removed the following day and replaced with CM differentiation media. After 3 days, CM differentiation media was replaced with lactate media (stock solution: 1M lactate, 1M Na-Hepes in distilled water; working solution: 4mM lactate in DMEM-no Glucose) for 4 days. From days 5-8, lactate media was titrated down in the following lactate-media: CM differentiation-media ratios: Day 5: 3:1; Day 6: 1:1; Day 7: 1:3; Day 8: 0:4 (Tohyama et al., 2013).

## Supporting information

Supplemental Tabels S1-S8

## Author Contributions

C.S. recruited, enrolled, scheduled health evaluation and biopsies of study participants and de-identified and coded samples; C.L., K. M., N.T. H. W. conducting the clinical health evaluations and biopsy punches of study participants; J.C.K. conceived the clinical health evaluation; D.T. and J.C.K. supervised the clinical health evaluation, evaluated all clinical data and made final determination on health-pass fail decision supervised and oversaw all clinical aspects of the study; A.S.Y. performed clinical health data extraction, processing and analysis; C.S., P.D., and T.R. established fibroblast lines; C.S., P.D., T.R., B.H., and S.L.D. generated hiPSCs; B.H. performed the tri-lineage differentiation; P.D., D.M.G. and N.C.D. differentiated hiPSCs to cardiomyocytes and performed the physiology experiments; D.M.G., R.D., E.A.S. and N.C.D. analyzed the physiology experiments; C.S., P.D., T.R. and G.J. performed immunostaining and imaging; B.H. and G.J. isolated mRNA and genomic DNA; J.T. and V.N. supervised and interpreted Mount Sinai karyotype analysis; K.C.M., C.S.L., H.W and M.T. conducted the clinical health evaluations; J.G. and M.A.M. read and interpreted ECG data; J.C.K. and D.T supervised clinical health evaluations and made final determination on subject eligibility; K.G.B. and R.S. supervised bulk and scRNA-seq library preparations, sequencing, and quality control; Y.X. and J.H. performed RNA-seq data processing and computational analysis; B.M., M.M., M.M. E.S. performed whole genome-sequencing data processing, ancestry, variant and recessive disorder carrier mutation analysis, and interpretation; S.I.B. performed direct dominant genetic clinical risk analysis; J.G. provided quality assurance and control; D.V. and S.C.S. coordinated and integrated data into the online LINSC data portal; C.S. and A.E. managed the project; C.S., M.B., E.A.S., N.C.D. and R.I. conceived, designed and supervised the study; C.S. and R.I. wrote the manuscript with input from all authors.

## Acknowledgements

This work was supported by NIH Common Fund LINCS program awards U54HG008098 (R.I.) and U54HL127624 (S.C.S). We thank the Mount Sinai Clinical Research Unit for providing exam rooms and nurses, and the Mount Sinai Genomics Core for mRNA sequencing.

## References

Carcamo-Orive, I., Hoffman, G.E., Cundiff, P., Beckmann, N.D., D’Souza, S.L., Knowles, J.W., Patel, A., Papatsenko, D., Abbasi, F., Reaven, G.M., et al. (2017). Analysis of Transcriptional Variability in a Large Human iPSC Library Reveals Genetic and Non-genetic Determinants of Heterogeneity. Cell Stem Cell 20, 518–532 e519.

D’Antonio-Chronowska, A., Donovan, M.K.R., Young Greenwald, W.W., Nguyen, J.P., Fujita, K., Hashem, S., Matsui, H., Soncin, F., Parast, M., Ward, M.C., et al. (2019). Association of Human iPSC Gene Signatures and X Chromosome Dosage with Two Distinct Cardiac Differentiation Trajectories. Stem Cell Reports 13, 924–938.

DeBoever, C., Li, H., Jakubosky, D., Benaglio, P., Reyna, J., Olson, K.M., Huang, H., Biggs, W., Sandoval, E., D’Antonio, M., et al. (2017). Large-Scale Profiling Reveals the Influence of Genetic Variation on Gene Expression in Human Induced Pluripotent Stem Cells. Cell Stem Cell 20, 533–546 e537.

Devalla, H.D., Schwach, V., Ford, J.W., Milnes, J.T., El-Haou, S., Jackson, C., Gkatzis, K., Elliott, D.A., Chuva de Sousa Lopes, S.M., Mummery, C.L., et al. (2015). Atrial-like cardiomyocytes from human pluripotent stem cells are a robust preclinical model for assessing atrial-selective pharmacology. EMBO Mol Med 7, 394–410.

Dubois, N.C., Craft, A.M., Sharma, P., Elliott, D.A., Stanley, E.G., Elefanty, A.G., Gramolini, A., and Keller, G. (2011). SIRPA is a specific cell-surface marker for isolating cardiomyocytes derived from human pluripotent stem cells. Nat Biotechnol 29, 1011–1018.

Fusaki, N., Ban, H., Nishiyama, A., Saeki, K., and Hasegawa, M. (2009). Efficient induction of transgene-free human pluripotent stem cells using a vector based on Sendai virus, an RNA virus that does not integrate into the host genome. Proc Jpn Acad Ser B Phys Biol Sci 85, 348–362.

Hagai, T., Chen, X., Miragaia, R.J., Rostom, R., Gomes, T., Kunowska, N., Henriksson, J., Park, J.E., Proserpio, V., Donati, G., et al. (2018). Gene expression variability across cells and species shapes innate immunity. Nature 563, 197–202.

Hu, S., Zhao, M.T., Jahanbani, F., Shao, N.Y., Lee, W.H., Chen, H., Snyder, M.P., and Wu, J.C. (2016). Effects of cellular origin on differentiation of human induced pluripotent stem cell-derived endothelial cells. JCI Insight 1.

International Stem Cell, I., Amps, K., Andrews, P.W., Anyfantis, G., Armstrong, L., Avery, S., Baharvand, H., Baker, J., Baker, D., Munoz, M.B., et al. (2011). Screening ethnically diverse human embryonic stem cells identifies a chromosome 20 minimal amplicon conferring growth advantage. Nat Biotechnol 29, 1132–1144.

Karch, C.M., Kao, A.W., Karydas, A., Onanuga, K., Martinez, R., Argouarch, A., Wang, C., Huang, C., Sohn, P.D., Bowles, K.R., et al. (2019). A Comprehensive Resource for Induced Pluripotent Stem Cells from Patients with Primary Tauopathies. Stem Cell Reports 13, 939–955.

Kaserman, J.E., Hurley, K., Dodge, M., Villacorta-Martin, C., Vedaie, M., Jean, J.C., Liberti, D.C., James, M.F., Higgins, M.I., Lee, N.J., et al. (2020). A Highly Phenotyped Open Access Repository of Alpha-1 Antitrypsin Deficiency Pluripotent Stem Cells. Stem Cell Reports 15, 242–255.

Kattman, S.J., Witty, A.D., Gagliardi, M., Dubois, N.C., Niapour, M., Hotta, A., Ellis, J., and Keller, G. (2011). Stage-specific optimization of activin/nodal and BMP signaling promotes cardiac differentiation of mouse and human pluripotent stem cell lines. Cell Stem Cell 8, 228–240.

Keenan, A.B., Jenkins, S.L., Jagodnik, K.M., Koplev, S., He, E., Torre, D., Wang, Z., Dohlman, A.B., Silverstein, M.C., Lachmann, A., et al. (2018). The Library of Integrated Network-Based Cellular Signatures NIH Program: System-Level Cataloging of Human Cells Response to Perturbations. Cell Syst 6, 13–24.

Kilpinen, H., Goncalves, A., Leha, A., Afzal, V., Alasoo, K., Ashford, S., Bala, S., Bensaddek, D., Casale, F.P., Culley, O.J., et al. (2017). Common genetic variation drives molecular heterogeneity in human iPSCs. Nature 546, 370–375.

Koleti, A., Terryn, R., Stathias, V., Chung, C., Cooper, D.J., Turner, J.P., Vidovic, D., Forlin, M., Kelley, T.T., D’Urso, A., et al. (2018). Data Portal for the Library of Integrated Network-based Cellular Signatures (LINCS) program: integrated access to diverse large-scale cellular perturbation response data. Nucleic Acids Res 46, D558–D566.

Landrum, M.J., Lee, J.M., Riley, G.R., Jang, W., Rubinstein, W.S., Church, D.M., and Maglott, D.R. (2014). ClinVar: public archive of relationships among sequence variation and human phenotype. Nucleic Acids Res 42, D980–985.

Lee, J.H., Protze, S.I., Laksman, Z., Backx, P.H., and Keller, G.M. (2017). Human Pluripotent Stem Cell-Derived Atrial and Ventricular Cardiomyocytes Develop from Distinct Mesoderm Populations. Cell Stem Cell 21, 179–194 e174.

Mannhardt, I., Saleem, U., Mosqueira, D., Loos, M.F., Ulmer, B.M., Lemoine, M.D., Larsson, C., Ameen, C., de Korte, T., Vlaming, M.L.H., et al. (2020). Comparison of 10 Control hPSC Lines for Drug Screening in an Engineered Heart Tissue Format. Stem Cell Reports 15, 983–998.

Mayshar, Y., Ben-David, U., Lavon, N., Biancotti, J.C., Yakir, B., Clark, A.T., Plath, K., Lowry, W.E., and Benvenisty, N. (2010). Identification and classification of chromosomal aberrations in human induced pluripotent stem cells. Cell Stem Cell 7, 521–531.

Nazor, K.L., Altun, G., Lynch, C., Tran, H., Harness, J.V., Slavin, I., Garitaonandia, I., Muller, F.J., Wang, Y.C., Boscolo, F.S., et al. (2012). Recurrent variations in DNA methylation in human pluripotent stem cells and their differentiated derivatives. Cell Stem Cell 10, 620–634.

Panopoulos, A.D., D’Antonio, M., Benaglio, P., Williams, R., Hashem, S.I., Schuldt, B.M., DeBoever, C., Arias, A.D., Garcia, M., Nelson, B.C., et al. (2017). iPSCORE: A Resource of 222 iPSC Lines Enabling Functional Characterization of Genetic Variation across a Variety of Cell Types. Stem Cell Reports 8, 1086–1100.

Park, I.H., Zhao, R., West, J.A., Yabuuchi, A., Huo, H., Ince, T.A., Lerou, P.H., Lensch, M.W., and Daley, G.Q. (2008). Reprogramming of human somatic cells to pluripotency with defined factors. Nature 451, 141–146.

Park, S., Gianotti-Sommer, A., Molina-Estevez, F.J., Vanuytsel, K., Skvir, N., Leung, A., Rozelle, S.S., Shaikho, E.M., Weir, I., Jiang, Z., et al. (2017). A Comprehensive, Ethnically Diverse Library of Sickle Cell Disease-Specific Induced Pluripotent Stem Cells. Stem Cell Reports 8, 1076–1085.

Peterson, S.E., and Loring, J.F. (2014). Genomic instability in pluripotent stem cells: implications for clinical applications. J Biol Chem 289, 4578–4584.

Pianezzi, E., Altomare, C., Bolis, S., Balbi, C., Torre, T., Rinaldi, A., Camici, G.G., Barile, L., and Vassalli, G. (2020). Role of somatic cell sources in the maturation degree of human induced pluripotent stem cell-derived cardiomyocytes. Biochim Biophys Acta Mol Cell Res 1867, 118538.

Ramamoorthy, A., Pacanowski, M.A., Bull, J., and Zhang, L. (2015). Racial/ethnic differences in drug disposition and response: review of recently approved drugs. Clin Pharmacol Ther 97, 263–273.

Rouhani, F., Kumasaka, N., de Brito, M.C., Bradley, A., Vallier, L., and Gaffney, D. (2014). Genetic background drives transcriptional variation in human induced pluripotent stem cells. PLoS Genet 10, e1004432.

Sanchez-Freire, V., Lee, A.S., Hu, S., Abilez, O.J., Liang, P., Lan, F., Huber, B.C., Ong, S.G., Hong, W.X., Huang, M., et al. (2014). Effect of human donor cell source on differentiation and function of cardiac induced pluripotent stem cells. J Am Coll Cardiol 64, 436–448.

Sankar, P., Cho, M.K., Condit, C.M., Hunt, L.M., Koenig, B., Marshall, P., Lee, S.S., and Spicer, P. (2004). Genetic research and health disparities. JAMA 291, 2985–2989.

Soldin, O.P., and Mattison, D.R. (2009). Sex differences in pharmacokinetics and pharmacodynamics. Clin Pharmacokinet 48, 143–157.

Stathias, V., Turner, J., Koleti, A., Vidovic, D., Cooper, D., Fazel-Najafabadi, M., Pilarczyk, M., Terryn, R., Chung, C., Umeano, A., et al. (2020). LINCS Data Portal 2.0: next generation access point for perturbation-response signatures. Nucleic Acids Res 48, D431–D439.

Streeter, I., Harrison, P.W., Faulconbridge, A., The HipSci, C., Flicek, P., Parkinson, H., and Clarke, L. (2017). The human-induced pluripotent stem cell initiative-data resources for cellular genetics. Nucleic Acids Res 45, D691–D697.

Taapken, S.M., Nisler, B.S., Newton, M.A., Sampsell-Barron, T.L., Leonhard, K.A., McIntire, E.M., and Montgomery, K.D. (2011). Karotypic abnormalities in human induced pluripotent stem cells and embryonic stem cells. Nat Biotechnol 29, 313–314.

Takahashi, K., Tanabe, K., Ohnuki, M., Narita, M., Ichisaka, T., Tomoda, K., and Yamanaka, S. (2007). Induction of pluripotent stem cells from adult human fibroblasts by defined factors. Cell 131, 861–872.

Tohyama, S., Hattori, F., Sano, M., Hishiki, T., Nagahata, Y., Matsuura, T., Hashimoto, H., Suzuki, T., Yamashita, H., Satoh, Y., et al. (2013). Distinct metabolic flow enables large-scale purification of mouse and human pluripotent stem cell-derived cardiomyocytes. Cell Stem Cell 12, 127–137.

Warren, L., Manos, P.D., Ahfeldt, T., Loh, Y.H., Li, H., Lau, F., Ebina, W., Mandal, P.K., Smith, Z.D., Meissner, A., et al. (2010). Highly efficient reprogramming to pluripotency and directed differentiation of human cells with synthetic modified mRNA. Cell Stem Cell 7, 618–630.

Wilson, J.F., Weale, M.E., Smith, A.C., Gratrix, F., Fletcher, B., Thomas, M.G., Bradman, N., and Goldstein, D.B. (2001). Population genetic structure of variable drug response. Nat Genet 29, 265–269.

Witty, A.D., Mihic, A., Tam, R.Y., Fisher, S.A., Mikryukov, A., Shoichet, M.S., Li, R.K., Kattman, S.J., and Keller, G. (2014). Generation of the epicardial lineage from human pluripotent stem cells. Nat Biotechnol 32, 1026–1035.

Yu, J., Vodyanik, M.A., Smuga-Otto, K., Antosiewicz-Bourget, J., Frane, J.L., Tian, S., Nie, J., Jonsdottir, G.A., Ruotti, V., Stewart, R., et al. (2007). Induced pluripotent stem cell lines derived from human somatic cells. Science 318, 1917–1920.

Zhang, Q., Jiang, J., Han, P., Yuan, Q., Zhang, J., Zhang, X., Xu, Y., Cao, H., Meng, Q., Chen, L., et al. (2011). Direct differentiation of atrial and ventricular myocytes from human embryonic stem cells by alternating retinoid signals. Cell Res 21, 579–587.

