## Supplemental Information and Figures for "A Library of Induced Pluripotent Stem Cells from Clinically Well-Characterized, Diverse Healthy Human Individuals"

**Supplemental Figures**
Figure S1. Immunocytochemistry of pluripotency markers for all 40 iPSC lines. *Related to Figure 2*.
Figure S2. PluriTest plots of 40 iPSC lines. *Related to Figure 2*.
Figure S3. Tri-lineage differentiation of iPSC lines. *Related to Figure 2*.
Figure S4. Correlation of cardiomyocyte characteristics between different hiPSC lines. *Related to Figure 6.*

**Supplemental Tables** (provided as separate sheets in a single Supplemental Tables excel file)
Table S1. Clinical characteristics of all 40 selected, clinically healthy subjects (Supplied as an excel-file only). *Related to Figure 1.*
Table S2. Summary characterization of all 40 selected, clinically healthy subjects and derived iPSCs. *Related to Figures 1, 2 and S1, and Tables S1, S3, S4 and S6* (Supplied also as an excel-file).
Table S3. Cytogenetics of 40 iPSC lines. *Related to Figures 1 and 2* (Supplied as an excel-file only).
Table S4. RNA-seq read counts. *Related to Figures 1, 2, 3, S2 and S4B* (Supplied also as an excel-file).
Table S5. PCA projections, standard deviation, variance and percent of variance of each principal component, and PCA variable loadings of hiPSC and hiPSC-derived cardiomyocyte RNA-seq data. *Related to Figure 3B and S4B* (Supplied as an excel-file only).
Table S6. Ancestry analysis. *Related to Figure 1* (Supplied as an excel-file only).
Table S7. Pathogenic disease variants. *Related to Figures 1 and 3A, and Tables S1 and S2* (Supplied as an excel-file only).
Table S8. Calcium transient data and analysis of hiPSC-derived cardiomyocytes. *Related to Figure S4A* and S6 (Supplied as an excel-file only).

**Supplemental Documents**
Document S1. General Study Consent. *Related to Figure 1.*
Document S2. HIV Consent. *Related to Figure 1*.
Document S3. Clinical Report Form. *Related to Figure 1 and Table S1*.

**Supplemental Experimental Procedures**
Describes the following procedures:

- Karyotyping
- Short-tandem repeat (STR) analysis
- Immunocytochemistry
- RNA isolation, library preparation and RNA-seq
- RNA-seq data analysis
- RNA-seq-based PluriTest analysis
- RNA-seq-based comparative transcriptome analysis
- Detection of Sendai Virus genomic RNA sequence
- WGS and Mendelian disease/disorder analysis
- Ancestry analysis
- Calcium transient recording and analysis

And also includes:

- Quantification and statistical analysis
- Data and software availability
- Additional resources
- DToxS_SOP_CE-6.0-Biopsy and Derivation of Fibroblasts
- DToxS_SOP_CE-7.0-mRNA/miRNA Reprogramming of Human Fibroblasts to Human Induced Pluripotent Stem Cells
- DToxS_SOP_CE-7.1-SeV Reprogramming of Human Fibroblasts to Human Induced Pluripotent Stem Cells

#### Schaniel et al. Figure S1

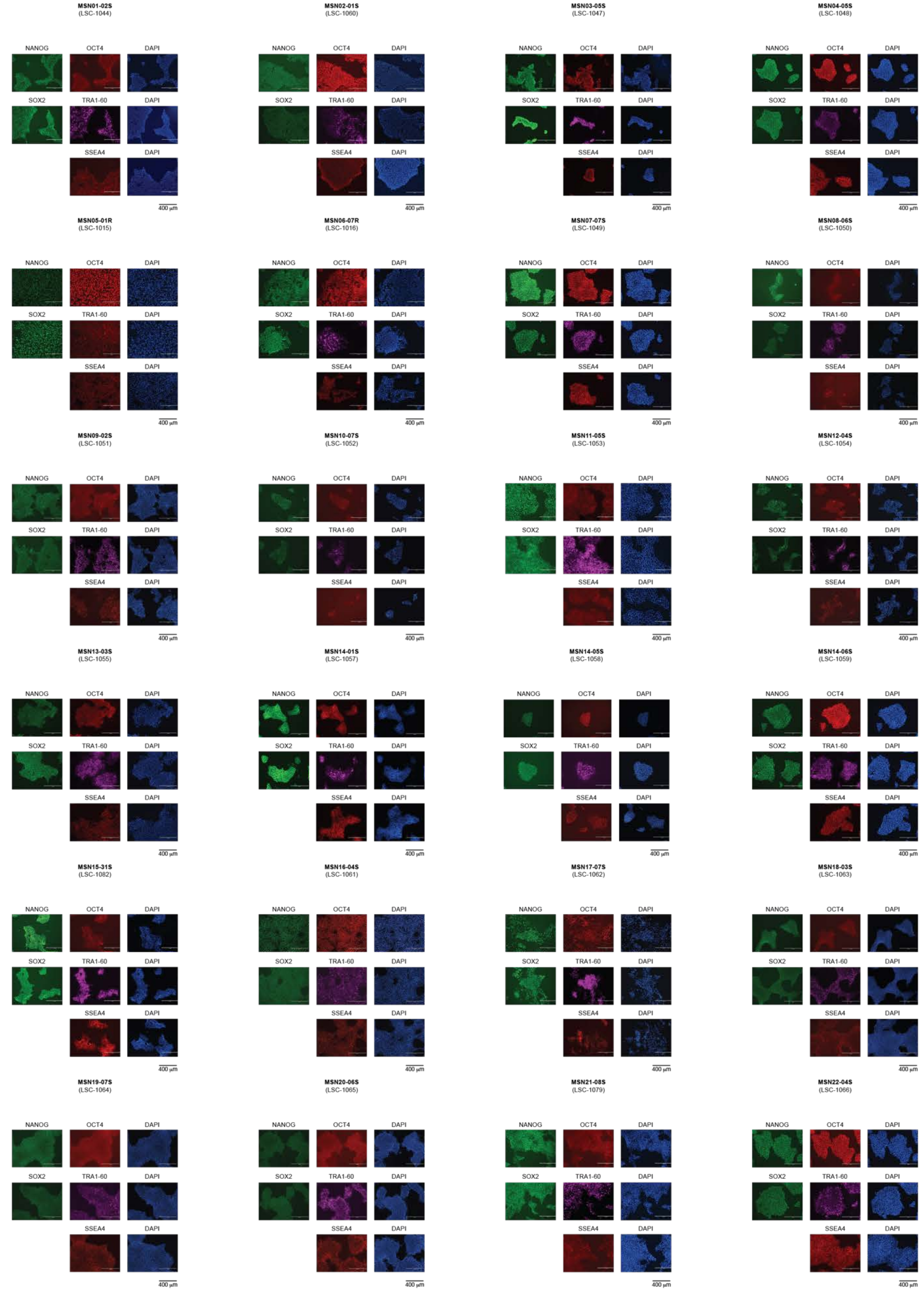

#### Schaniel et al. Figure S1 (continued)

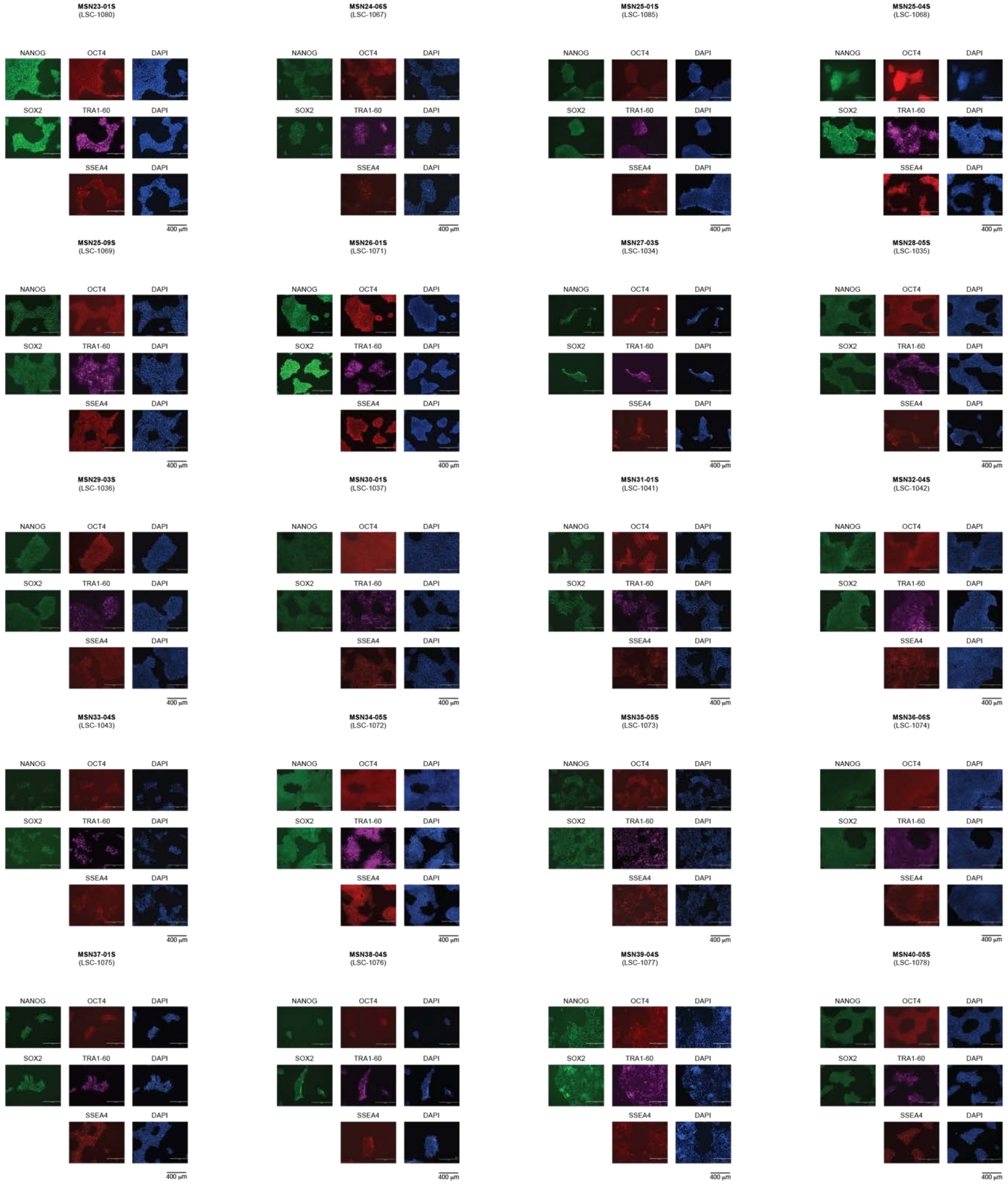

**Figure S1. Immunocytochemistry of pluripotency markers for all 40 iPSC lines.** ***Related to Figure 2.*** Immunocytochemistry of pluripotency markers, NANOG, OCT4, SOX2, TRA-1-60 and SSEA4 of one representative iPSC clone for each of the 40 selected, clinically healthy subjects. DAPI is used to stain nuclei. Bar, 400 µm.

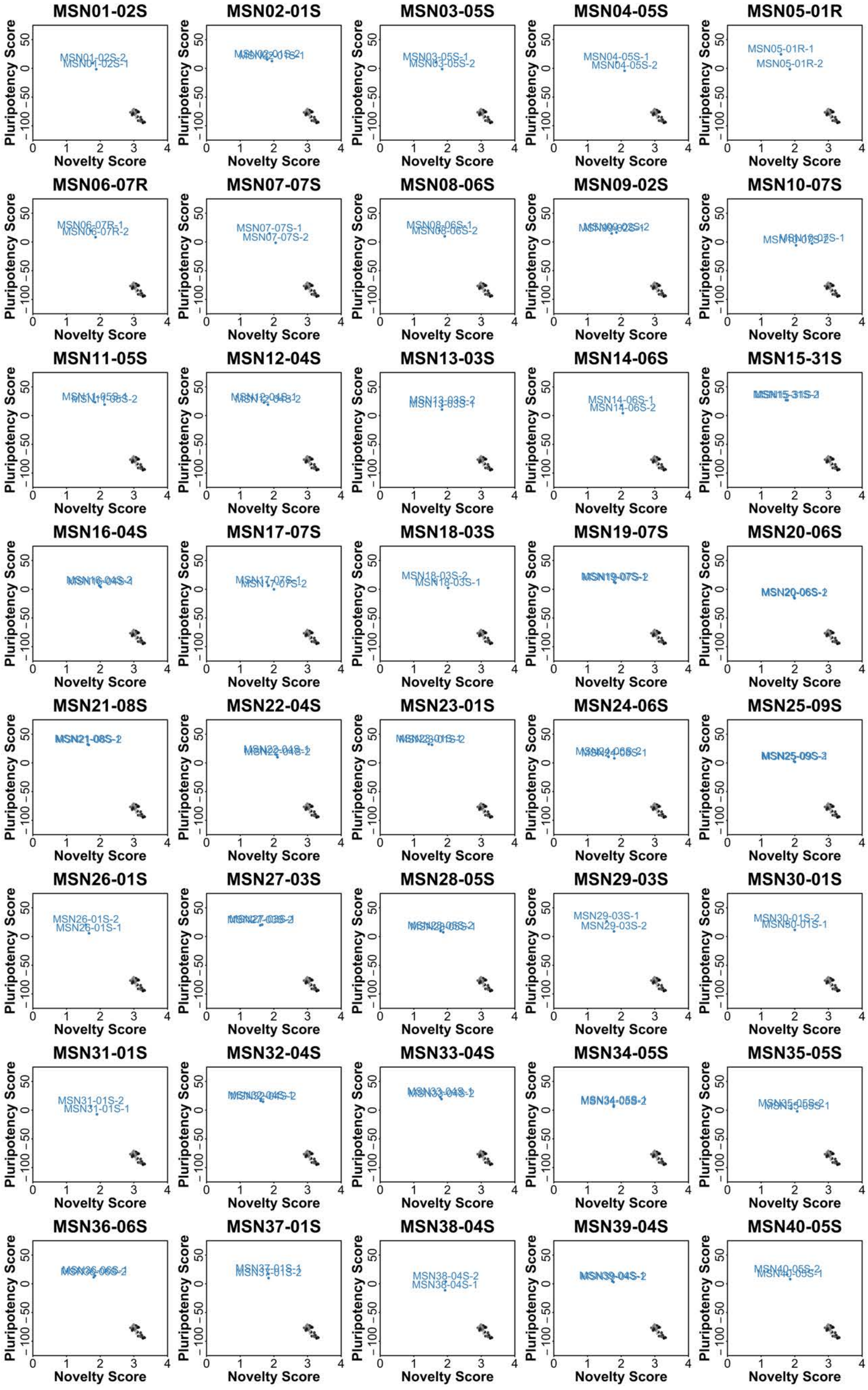

**Figure S2. PluriTest plots of 40 iPSC lines.** ***Related to Figure 2.*** PluriTest plots based on mRNA-seq transcriptomic analyses performed in duplicate for one representative iPSC clone for each of the 40 selected, clinically healthy subjects (colored circles). As a reference, the PluriTest results of transcriptomic data of fibroblast from 66 individuals are plotted in black/grey.

Schaniel et al. Figure S3

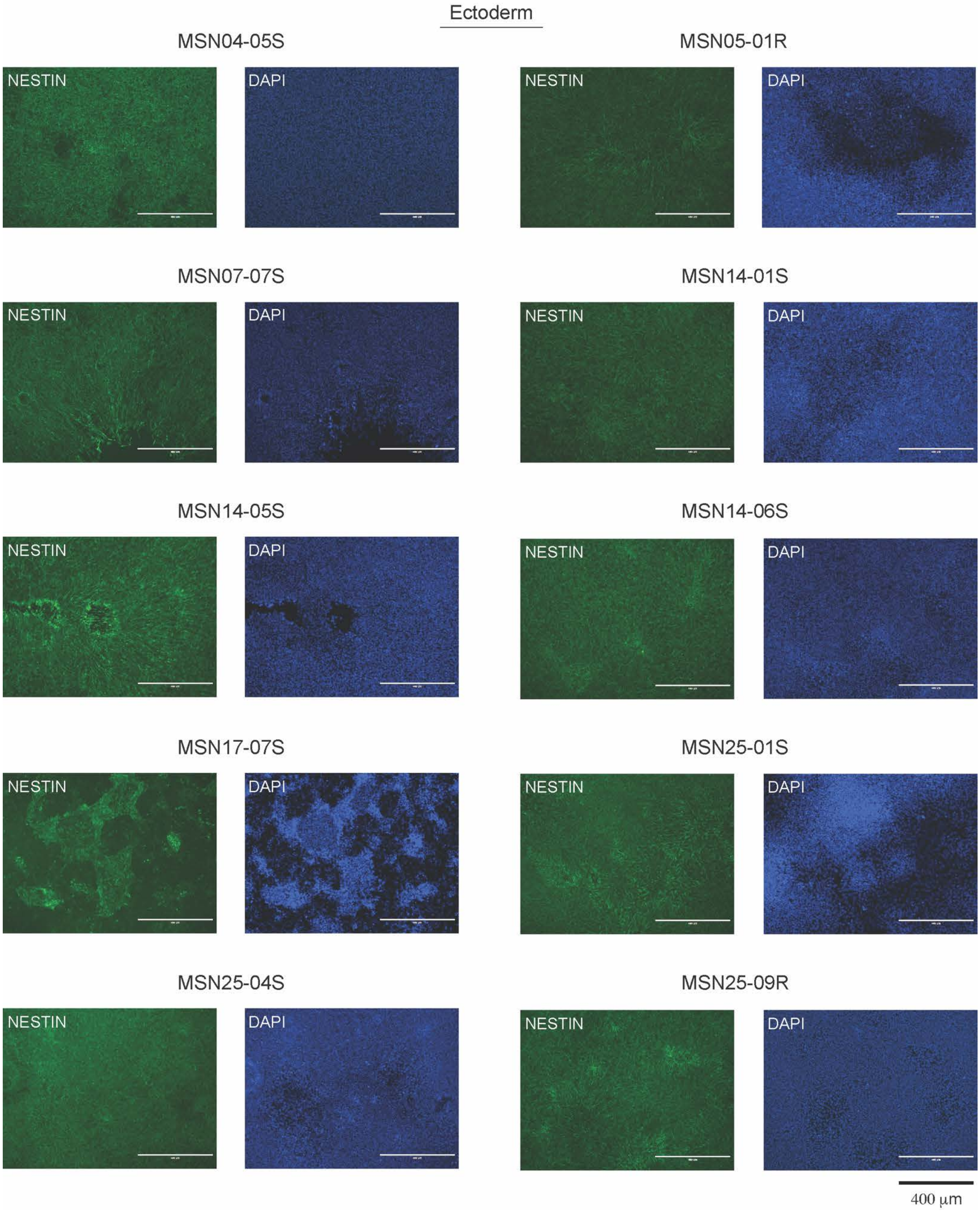

**Schaniel et al. Figure S3 (continued)**

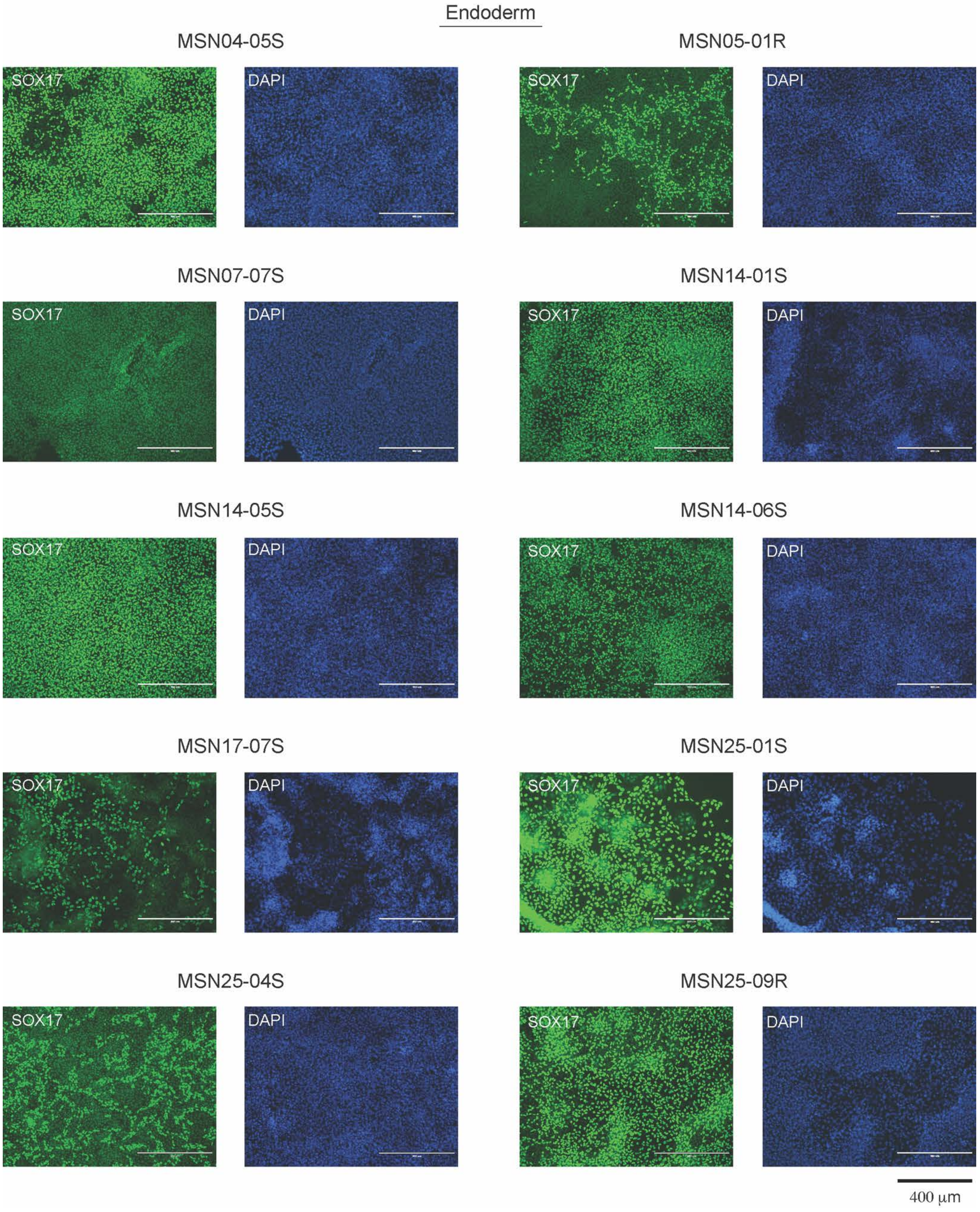

**Schaniel et al. Figure S3 (continued)**

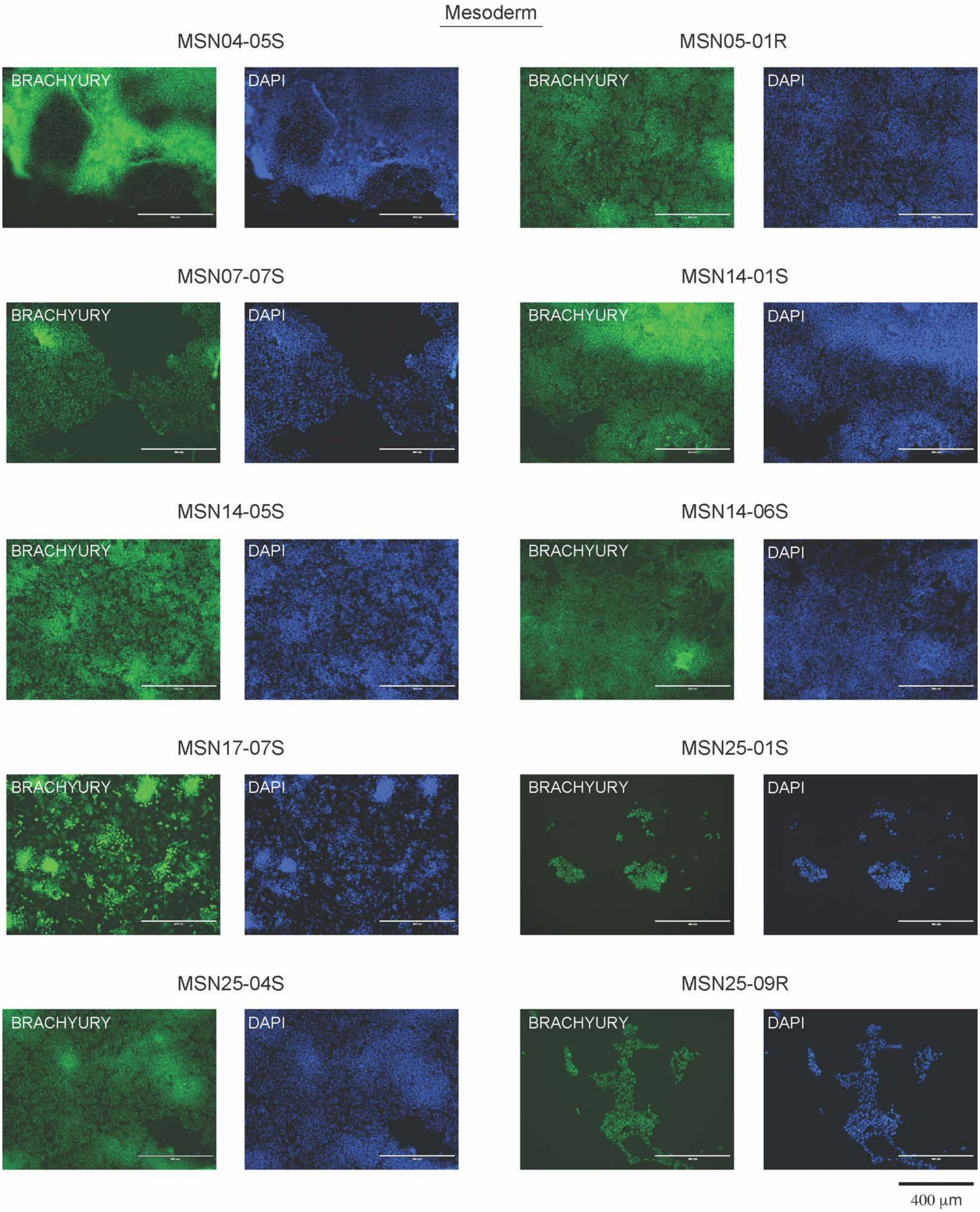

**Figure S3. Tri-lineage differentiation of iPSC lines.** ***Related to Figure 2.*** Immunocytochemistry of a selection of hiPSC lines and clones differentiated independently to ectoderm, endoderm and mesoderm. NESTIN, SOX17 and BRACHYURY expression is used to represent cells of ectodermal, endodermal and mesodermal origin, respectively. DAPI is used to stain nuclei. Bar, 400 µm.

Schaniel et al. Figure S4

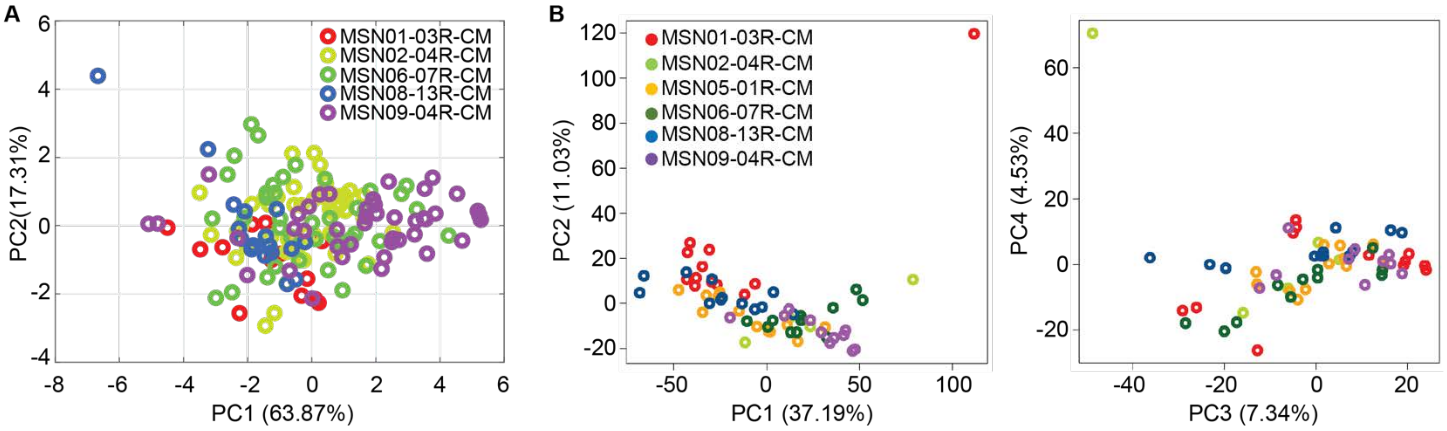

C

| | Cell counts | Mean number of unique reads per cell | Mean number of expressed genes per cells | Mean percent of mitochondrial reads per cells |
|---|---|---|---|---|
| MSN01-03R-CM | 3,281 | 25,354 | 4,836 | 17.28 |
| MSN02-04R-CM | 2,748 | 24,871 | 5,100 | 9.70 |
| MSN08-13R-CM | 3,996 | 25,354 | 4,699 | 7.72 |
| MSN09-04R-CM | 4,465 | 25,354 | 4,650 | 14.92 |

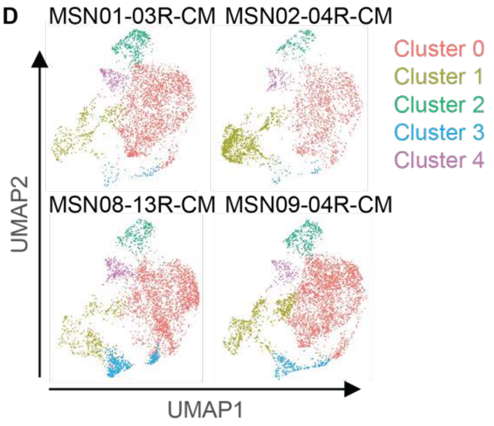

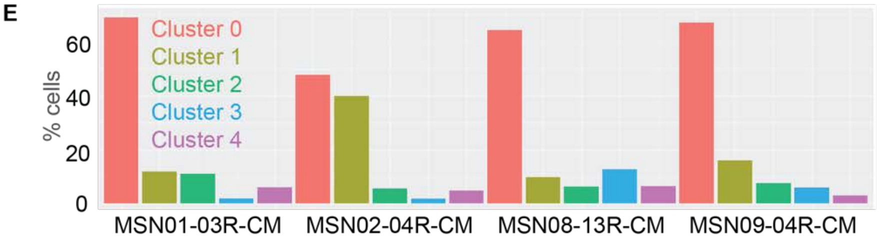

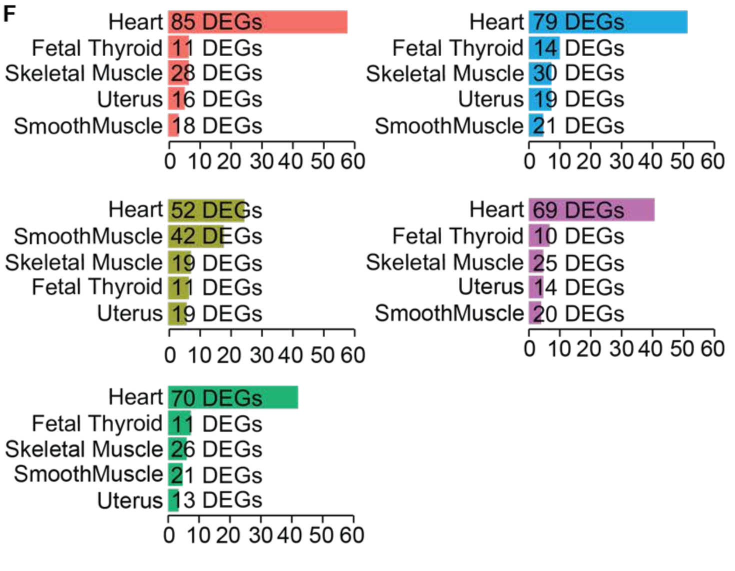

G

| Cell type | Gene | Cluster 0 | Cluster 1 | Cluster 2 | Cluster 3 | Cluster 4 |
|---|---|---|---|---|---|---|
| CM | ACTN2 | 0.83 | | | | 0.39 |
| CM | GATA4 | 0.19 | | | 0.13 | 0.41 |
| CM | MEF2C | | 0.02 | | 0.52 | 0.23 |
| CM | MYL7 | 0.49 | | | 0.66 | 0.53 |
| CM | MKX2-5 | 0.65 | | | | 0.57 |
| CM | SIRPA | 0.13 | | | | 0.29 |
| CM | TNNT2 | 0.79 | | | 0.24 | |
| EPDC-SMC | ACTA2 | | 1.66 | | 0.97 | 0.04 |
| EPDC-SMC | CNN1 | | 0.12 | | | 0.41 |
| EPDC-Fib | DDR2 | | 0.94 | | 0.33 | |
| EPDC-SMC | PDGFRB | | 2.13 | | | |
| EPDC-SMC | SNAI2 | | 1.15 | | | |
| EPDC-SMC | TAGLN | | 1.80 | | | |
| EPDC-SMC | TAGLN2 | | 1.64 | | | 0.29 |
| EPDC-Fib | THY1/CD90 | | 2.46 | | | |
| EPDC-SMC | VIM | | 2.01 | 0.22 | | 0.51 |

**Figure S4. Correlation of cardiomyocyte characteristics between different hiPSC lines.** ***Related to Figure 6*****. A.** PCA of 7 features ($Ca^{2+}$-amplitude; $Ca^{2+}$D50; $Ca^{2+}$D90; Time to peak; decay; transient duration; and frequency) from calcium transient measurements on 162 single ventricular cardiomyocytes derived from the five indicated hiPSC lines (MSN01-03R, 14 cells; MSN02-04R, 48 cells; MSN06-07R, 40 cells; MSN08-13R, 17 cells; and MSN09-04R, 43 cells, from 2-3 independent differentiations per line). **B.** PCA of bulk RNA-seq data obtained from ventricular cardiomyocytes differentiated from the indicated hiPSC lines after 2-day treatment with DMSO at 14.1 mM. Reads were mapped to the reference genome, followed by counting the reads per gene. Genes that were not detected in at least 80% of all cells were removed, $log_{10}$ transformed and centered the read count matrix and subjected it to PCA. Replicates (5-12 per hiPSC line, see also Table S4) are visualized in dependence of the first and second principal components (left panel), and the third and fourth principal components (right panel). **C.** Metrics of scRNA-seq of cardiomyocytes differentiated from the indicated four human iPSC lines after quality control. **D.** UMAP of the hiSPC-derived cardiomyocyte scRNA-seq data identifies five clusters. **E.** Relative distribution of the cells of each sample over the five clusters. **F.** Enrichment analysis of the top 500 most abundantly expressed genes in each cluster using Fisher's Exact Test and a library that contains human tissue and cell-type specific genes. DEGs, differentially expressed genes; x-axis, -$Log_{10}$ p-values. **G.** Cluster marker genes were obtained by calculation of differentially expressed genes in each cluster compared to all other clusters. $Log_2$ fold changes of cell type specific genes that allow differentiation of cardiomyocytes (CM) from epicardial derived cells (EPDC) (D'Antonio-Chronowska et al., 2019) are shown. SMC, smooth muscle cell; Fib, Fibroblast.

**Document S1. General Study Consent.** ***Related to Figure 1.***

IRB Approved

**ICAHN SCHOOL OF MEDICINE AT MOUNT SINAI AND THE MOUNT SINAI HOSPITAL**
**CONSENT FORM TO VOLUNTEER IN A RESEARCH STUDY**
**AND AUTHORIZATION FOR USE AND DISCLOSURE OF MEDICAL INFORMATION**

**Study ID #: HSM14-00530** **Form Version Date: February 11, 2015**

#### TITLE OF RESEARCH STUDY:

Title: **Drug Combination Signatures for Prediction and Mitigation of Toxicity**

#### PRINCIPAL INVESTIGATOR (HEAD RESEARCHER) NAME AND CONTACT INFORMATION:

Name: Ravi Iyengar, PhD

Physical Address: Icahn Medical Institute building, 12-70C

Mailing Address: One Gustave L. Levy Place
Box 1215
New York, NY 10029

#### WHAT IS A RESEARCH STUDY?

A research study is when scientists try to answer a question about something that we don't know enough about. Participating may not help you or others.

People volunteer to be in a research study. The decision about whether or not to take part is totally up to you. You can also agree to take part now and later change your mind. Whatever you decide is okay. It will not affect your ability to get medical care at Mount Sinai.

Someone will explain this research study to you. Feel free to ask all the questions you want before you decide. Any new information that develops during this research study that might make you change your mind about participating will be given to you promptly.

Basic information about this study will appear on the website http://www.ClinicalTrials.gov. There are a few reasons for this: the National Institutes of Health (NIH) encourages all researchers to post their research; some medical journals only accept articles if the research was posted on the website; and, for research studies the U.S. Food and Drug Administration (FDA) calls "applicable clinical trials" a description of this clinical trial will be available on http://www.ClinicalTrials.gov, as required by U.S. Law. This Web site will not include information that can identify you. At most, the Web site will include a summary of the results. You can search this Web site at any time.

#### PURPOSE OF THIS RESEARCH STUDY:

The purpose of this study is to understand adverse events caused by useful drugs in normal tissues of humans. For this we plan to obtain skin samples of healthy human subjects. Using this skin sample, we intend to establish fibroblast lines that will subsequently be reprogrammed to cells with the potential to give rise to any cell of the body and then differentiated to cells of the heart, liver and peripheral sensory (neurons) system for the purpose of the generation of molecular signatures for prediction and reduction of drug side effects caused by an specific drug and drug combinations.

Another purpose of this research project is to collect and store human samples (such as skin samples) and health information. Researchers can then use the stored materials in future studies. Through such studies, they hope to find new ways to detect, treat, and maybe even prevent or cure

IRB Approved

health problems. Some of these studies may be about how genes affect health, or how genes affect response to treatment. (Genes, which are made up of DNA, have the information needed to build and operate a human body.) Some of the studies may lead to new products, such as drugs or tests for diseases.

You may qualify to take part in this research study because your examination by the clinical research center indicates that you are a healthy person between the ages of 18-65.

Funds for conducting this research are provided by NIH.

#### LENGTH OF TIME AND NUMBER OF PEOPLE EXPECTED TO PARTICIPATE

Your participation in this research study is limited to a maximum of two visits. During the first visit to the clinical research center you will give informed consent and only then undergo a physical exam to establish your healthy status. Once your healthy status has been determined you will be contacted again to schedule an appointment during which we will obtain a skin sample. This visit will will take approximately 10 to 15 minutes to complete the skin biopsy. The tissue sample will be collected during a clinic visit arranged at your convenience and will require no extra time involvement after obtaining the tissue specimen.

The total number of people expected to take part in this research study is 30 individuals between 18-65 with no gender bias but about 30% ethnic minorities (African American, Asians and Hispanics).

#### DESCRIPTION OF WHAT'S INVOLVED:

If you agree to participate in this research study, the following information describes what may be involved.

During a visit to the clinical research center you will give informed consent prior to undergoing a physical exam to establish your healthy status. The exam will involve assessment of your clinically history, taking of vital signs and major system examinations (including pulmonary, cardiovascular, GI), an electrocardiogram (ECG; describe in the next paragraph) and having blood drawn (18 mL) to test heart, liver and sensory/touch system function parameters. Your blood will also be tested for HIV. HIV is the term used for the virus that produces HIV infection and may ultimately lead to AIDS. You must be told that you are being tested for HIV and give consent. You will sign an additional consent form prior to the HIV blood test. You have a right to know the results of the test. If you test positive you will be given more tests to confirm the results of the first test. If you are HIV positive, the study doctor will also give you a list of referrals for further information and counseling. Female subjects only will have to provide a urine sample to determine their pregnancy status in order to rule out any eventual problems from the lidocaine (for local anesthesia) and the skin punch biopsy. Specifically, the physical exam will involve first checking of weight, height, waist and hip circumference, heart rate, blood pressure, rate of breathing (breaths per minute) and peripheral oxygen saturation by pulse oximeter placed on a finger (this is a non-invasive measurement that takes only 10-20 seconds and is not painful). The doctor will then perform a complete physical examination that will include examination of the cardiovascular, respiratory, gastrointestinal and neurological systems. Particular aspects will involve listening to the heart and lungs with a stethoscope, palpation (feeling with the hands) of the abdomen, and examination of the sensori-motor system. A rectal exam will not be performed. This complete physical examination will take about 10-15 minutes.

Icahn School of Medicine at Mount Sinai
IRB Approved
Program for the Protection of Human Subjects

An ECG is the recording of the electrical activity of the heart captured over time by external electrodes attached to the skin. This involves placing 10 recording electrodes on the skin, with a gel-like substance contacting the skin (which can conduct the heart electricity). Six electrodes go across the left chest, while one electrode is attached to each limb (left and right arm and leg). For persons with a "hairy chest", shaving of some of the chest hair may be required to achieve good contact of the sensing electrodes and the skin. The entire process takes 5-10 minutes and is painless. Because an ECG is only recording the activity of the heart, there are no "electrical" or other sensations felt.
If you continue to qualify to take part in this study, you will have a skin sample taken at your convenience. If it is determined that you are not healthy and cannot continue in this study, you will be provided the results of the test by the study physician and be recommendation to see your personal physician. If you do not have a personal physician, we will refer you to the Mount Sinai Internal Medicine Associates.

After determination of your healthy status, a skin samples will be collected during a second visit arranged at your convenience. No further time involvement is required. We will obtain and save these samples and store the tissue in a bank with other samples that will be used for the research testing described as above and will be stored for a minimum of five years and for as long as deemed useful for research purposes. The skin sample will be assigned a 4-digit number (de-identification) for confidentiality during the research study.

The skin biopsy is called a punch biopsy and involves using a small 2 mm bore needle to obtain a sample of skin from either the forearm or behind the knee. The size of the sample taken will be approximately similar to the size of this typed letter "O", approximately 2 mm in size. A numbing anesthetic will be injected locally to numb the area before the sample is obtained. The area will then be covered with a small gauze or bandage. The risks involved are listed below. The punch biopsy will be obtained by a clinician who is a member of our research team.

We may wish to do future testing of your skin samples and derivatives, including but not limited to generated induced pluripotent stem cells and differentiated target cells such as heart, liver and peripheral nerve cells, based on potential future scientific developments and discoveries.

The researchers would like to ask your permission to keep specimens (like blood, tissue, hair, or any other body matter) collected or derived from you during this study to use them in future research studies. They would also like to know your wishes about how they might use your specimens in future research studies. You should also know that it is possible that products may someday be developed with the help of your specimens, and there are no plans to share any profits from such products with you.

***(1)** Will you allow the researchers to store your specimens to use in future research studies?*

*Yes________ No________ If no, please stop here. If yes, please continue to the next question.*

**(2)** The researchers can keep your specimens stored in one of two different ways: one way will store your specimens in a way that it is linked to your identity (through the use of a code that can indicate the information came from you personally) and the other way will store your specimens anonymously (no one will know who the information is from). It will not be stored both ways, so you must choose one of these two options. Please note that if you choose to have your specimens stored anonymously, you will not be able to change your mind to ask for your specimens to be destroyed at a future date.

How would you like your specimens stored? Please initial **ONE** choice:
I would like my specimens stored with a link to my identity__________
I would like my specimens stored anonymously __________

**(3)** Do you give the researchers permission to **contact you** in the future to collect additional information about you, discuss how your specimens might be used, or to discuss possible participation in another research project? Please initial your choice:

*Yes*__________ *No*__________

***(4)** Do you give the researchers permission to keep the specimens indefinitely and use them for future studies that are **directly related** to the purpose of the current study? Please initial your choice:*

*Yes*__________ *No*__________

***(5)** Do you give the researchers permission to keep the specimens indefinitely and use them for future studies that are **not related** to the purpose of the current study (for example, a different area of research)? Please initial your choice:*

*Yes*__________ *No*__________

**(a)** If the future research in a different area can be done without having to know that the specimens came from you personally, that will be done.

**(b)** If the future research in a different area requires that it is known specifically who the specimens came from, then one of the following will be done:

**(i)** If you allowed the researchers to contact you in the future, they will be able to contact you to explain why your specimen is needed and what will be done with it. Your permission will be asked to use your specimens in that research project.

**(ii)** If you do not give permission to be contacted in the future, or if it is found that contacting you is not practical, for example, because you have moved, your specimens may still be used. Either all links to your identity will be removed from the specimens, or an Institutional Review Board will be asked for permission to use the specimens linked to your identity. The Institutional Review Board (IRB) is a committee of doctors

Icahn School of Medicine at Mount Sinai Program for the Protection of Human Subjects IRB Approved

and scientists and non-scientists and people not associated with this hospital or medical school whose job it is to protect people who participate in research. The IRB can give permission for researchers to use and share health information connected to specimens that are linked to people's identities, but only if it determines that doing this will not be more than a minimal risk to people or their privacy.

**(6)** Do you give permission to have portions of the specimens **given to other researchers** at Mount Sinai or other institutions for use in research that is either related or not related to the purpose of this study? Please initial your choice:

*Yes*_________ *No*__________

As part of this study and do more powerful research (also in the future), it is helpful for researchers to share information they get from studying human samples. The scientific data collected in this study (but not your personal information such as name, address or social security number; see also below) will be deposited with the National Institutes of Health (an agency of the federal government) funded Data Coordination and Integration Center for the LINCS-BDK2 consortium, and possibly more scientific databases that may be maintained by Mount Sinai, by the federal government, or even by private companies, where it is stored along with information from other studies. Researchers can then study the combined information to learn even more about health and disease. A researcher who wants to study the information must adhere to rules set for by the Data Coordination and Integration Center for the LINCS-BDK2 consortium and conjunction with National Institutes of Health guidelines. Your name and other information that could directly identify you (such as address or social security number) will never be placed into a scientific database. However, because your genetic information is unique to you, there is a small chance that someone could trace it back to you. The risk of this happening is very small, but may grow in the future. Researchers will always have a duty to protect your privacy and to keep your information confidential.

In general, results from this research project will not be shared with you or put into your medical records. In some situations, the results (including genome sequencing results) might be important to your health or medical care. If this occurs, we will contact you to see if you want to learn more and if so refer you to your personal physician.

*No tests other than those authorized will be performed on the skin sample.*

#### YOUR RESPONSIBILITIES IF YOU TAKE PART IN THIS RESEARCH:

If you decide to take part in this research study you will be responsible for undergoing a physical exam, an electrocardiogram and provide a blood sample for lab tests to establish your healthy status and only if such is confirmed to provide a skin sample, which will be obtained at a separate visit arranged at your convenience.

#### COSTS OR PAYMENTS THAT MAY RESULT FROM PARTICIPATION:

Icahn School of Medicine at Mount Sinai
IRB Approved
Program for the Protection of Human Subjects

If you agree to take part in this research study, after completion of the skin punch biopsy, we will pay you $150 in cash as reimbursement for your time and effort.

Tax law may require the Mount Sinai Finance Department to report the amount of payment you receive from Mount Sinai to the Internal Revenue Service (IRS) or other agencies, as applicable. Generally this reporting would take place if you receive payments that equal $600 or more from Mount Sinai in a calendar year. You would be responsible for the payment of any tax that may be due.

#### POSSIBLE BENEFITS:

You are not expected to get any benefit from taking part in this research study. Others may not benefit either. However, possible benefits include gaining valuable knowledge about specific drug and/or drug combinations and why they may be toxic on a molecular level to cells of the heart, liver and peripheral sensory system (neurons). Even if such knowledge is gained it will not be shared with you.

#### REASONABLY FORESEEABLE RISKS AND DISCOMFORTS:

Possible risks and discomforts include those associated with obtaining the skin biopsy, which are bleeding, localized collection of blood outside the blood vessels, infection, and pain and/or physical discomfort at the time of the sampling. The sampling will be done during a routine clinic visit. Risks of the local anesthetic Lidocaine, used for skin biopsy, are pain at the site of sampling/injection. Additionally, lidocaine can pass very quickly through the placenta and reports exist of negative side effects on infants born to mothers using the anesthetic. Several lidocaine studies with human case reports have showed no negative side effects, pregnancy complications or birth defects. In addition, there is minimal risk for an allergic reaction to the local anesthetic. However, allergic reactions are immediate and since you will be watched by the study doctor during the procedure, the doctor can will watch for and treat any allergic reaction that may occur.

<u>Group Risks</u> □
Although we will not give researchers your name, we will give them basic information such as your race, ethnic group, and sex. This information helps researchers learn whether the factors that lead to health problems are the same in different groups of people. It is possible that such findings could one day help people of the same race, ethnic group, or sex as you. However, they could also be used to support harmful stereotypes or even promote discrimination. □

<u>Privacy Risks</u>
There always exists the potential for loss of private information; however, there are procedures in place to minimize this risk.

Your privacy is very important to us and we will use many safety measures to protect your privacy. However, in spite of all of the safety measures that we will use, we cannot guarantee that your identity will never become known. Although your genetic information is unique to you, you do share some genetic information with your children, parents, brothers, sisters, and other blood relatives. Consequently, it may be possible that genetic information from them could be used to help identify

IRB Approved

you. Similarly, it may be possible that genetic information from you could be used to help identify them. □

As part of this study and do more powerful research (also in the future), it is helpful for researchers to share information they get from studying human samples. The scientific data collected in this study (but not your personal information such as name, address or social security number; see also below) will be deposited with the National Institutes of Health funded Data Coordination and Integration Center for the LINCS-BDK2 consortium, and possibly more scientific databases, where it is stored along with information from other studies. Researchers can then study the combined information to learn even more about health and disease. A researcher who wants to study the information must adhere to rules set for by the Data Coordination and Integration Center for the LINCS-BDK2 consortium and conjunction with National Institutes of Health guidelines.

Your name and other information that could directly identify you (such as address or social security number) will never be placed into a scientific database. However, because your genetic information is unique to you, there is a small chance that someone could trace it back to you. The risk of this happening is very small, but may grow in the future. Since the database includes genetic information, a break in security may also pose a potential risk to blood relatives as well as yourself. For example, it could be used to make it harder for you (or a relative) to get or keep a job or insurance. If your private information was misused it is possible you would also experience other harms, such as stress, anxiety, stigmatization, or embarrassment from revealing information about your family relationships, ethnic heritage, or health conditions.

People may develop ways in the future that would allow someone to link your genetic or medical information in our databases back to you. For example, someone could compare information in our databases with information from you (or a blood relative) in another database and be able to identify you (or your blood relative). It also is possible that there could be violations to the security of the computer systems used to store the codes linking your genetic and medical information to you. □□

Since some genetic variations can help to predict the future health problems of you and your relatives, this information might be of interest to health providers, life insurance companies, and others. Patterns of genetic variation also can be used by law enforcement agencies to identify a person or his/her blood relatives. Therefore, your genetic information potentially could be used in ways that could cause you or your family distress, such as by revealing that you (or a blood relative) carry a genetic disease. □

There is a Federal law called the Genetic Information Nondiscrimination Act (GINA). In general, this law makes it illegal for health insurance companies, group health plans, and most large employers to discriminate against you based on your genetic information. However, it does not protect you against discrimination by companies that sell life insurance, disability insurance, or long-term care insurance.

There also may be other privacy risks that we have not foreseen.

#### OTHER POSSIBLE OPTIONS TO CONSIDER:

Icahn School of Medicine at Mount Sinai IRB Approved Program for the Protection of Human Subjects

You may decide not to take part in this research study without any penalty. The choice is totally up to you.

#### IN CASE OF INJURY DURING THIS RESEARCH STUDY:

If you believe that you have suffered an injury related to this research as a participant in this study, you should contact the Principal Investigator.

#### ENDING PARTICIPATION IN THE RESEARCH STUDY:

You may stop taking part in this research study at any time without any penalty. This will not affect your ability to receive medical care at Mount Sinai or to receive any benefits to which you are otherwise entitled.

If you decide to stop being in the research study, please contact the Principal Investigator or the research staff.

You may also withdraw your permission for the use and disclosure of any of your protected information for research, but you must do so in writing to the Principal Investigator at the address on the first page. Even if you withdraw your permission, the Principal Investigator for the research study may still use the information that was already collected if that information is necessary to complete the research study. Your health information may still be used or shared after you withdraw your authorization if you should have an adverse event (a bad effect) from participating in the research study.

Withdrawal without your consent: The study doctor, the sponsor or the institution may stop your involvement in this research study at any time without your consent. This may be because the research study is being stopped, the instructions of the study team have not been followed, the investigator believes it is in your best interest, or for any other reason. If specimens or data have been stored as part of the research study, they too can be destroyed without your consent.

If you chose to end your participation in this study, all specimens, i.e. fibroblasts derived from the biopsy (skin sample) as well as the induced pluripotent stem cells generated from those fibroblasts can easily be discarded/destroyed.

#### CONTACT PERSON(S):

If you have any questions, concerns, or complaints at any time about this research, or you think the research has hurt you, please contact the office of the research team and/or the Principal Investigator at phone number (212)-659-1707 or Christoph Schaniel, Ph.D., at (212)-659-8276.

If you experience an emergency during your participation in this research, contact Dr. Jason Kovacic at (212)-241-7300.

This research has been reviewed and approved by an Institutional Review Board. You may reach a representative of the Program for Protection of Human Subjects at the Icahn School of Medicine at Mount Sinai at telephone number (212) 824-8200 during standard work hours for any of the following reasons:

- Your questions, concerns, or complaints are not being answered by the research team.

Icahn School of Medicine at Mount Sinai
IRB Approved
Program for the Protection of Human Subjects

- You cannot reach the research team.
- You are not comfortable talking to the research team.
- You have questions about your rights as a research subject.

You want to get information or provide input about this research.

#### DISCLOSURE OF FINANCIAL INTERESTS:

None.

#### MAINTAINING CONFIDENTIALITY – HIPAA AUTHORIZATION:

As you take part in this research project it will be necessary for the research team and others to use and share some of your private protected health information. Consistent with the federal Health Insurance Portability and Accountability Act (HIPAA), we are asking your permission to receive, use and share that information.

What protected health information is collected and used in this study, and might also be disclosed (shared) with others?

As part of this research project, the researchers will collect your name, address, telephone number, and medical record number at Mount Sinai Hospital.

The researchers will also get information from your medical record from Mount Sinai Hospital.

During the study the researchers will gather information by:

- taking a medical history (includes current and past medications or therapies, illnesses, conditions or symptoms, family medical history, allergies, etc.)
- completing the tests, procedures, questionnaires and interviews explained in the description section of this consent.

Why is your protected health information being used?

Your personal contact information is important to be able to contact you during the study. Your health information and the results of any tests and procedures being collected as part of this research study will be used for the purpose of this study as explained earlier in this consent form. The results of this study could be published or presented at scientific meetings, lectures, or other events, but would not include any information that would let others know who you are, unless you give separate permission to do so.

The research team and other authorized members of The Mount Sinai Hospital and Mount Sinai School of Medicine (together, "Mount Sinai") workforce may use and share your information to ensure that the research meets legal, institutional or accreditation requirements. For example, the Mount Sinai School of Medicine Program for the Protection of Human Subjects is responsible for overseeing research on human subjects, and may need to see your information. If you receive any payments for taking part in this study, the Mount Sinai Medical Center Finance Department may need your name, address, social security number, payment amount, and related information for tax reporting purposes.

IRB Approved

If the research team uncovers abuse, neglect, or reportable diseases, this information may be disclosed to appropriate authorities.

Who, outside Mount Sinai, might receive your protected health information? None.

As part of the study, the Principal Investigator, study team and others in the Mount Sinai workforce may disclose your protected health information, including the results of the research study tests and procedures, to the following people or organizations:

- The United States Department of Health and Human Services and the Office of Human Research Protection.
- The sponsoring government agency and/or their representative who need to confirm the accuracy of the results submitted to the government or the use of government funds: The National Institutes of Health.

(It is possible that there may be changes to the list during this research study; you may request an up-to-date list at any time by contacting the Principal Investigator.)

In all disclosures outside of Mount Sinai, you will not be identified by name, social security number, address, telephone number or any other direct personal identifier unless disclosure of the direct identifier is required by law. Some records and information disclosed may be identified with a unique code number. The Principal Investigator will ensure that the key to the code will be kept in a locked file, or will be securely stored electronically. The code will not be used to link the information back to you without your permission, unless the law requires it, or rarely if the Institutional Review Board allows it after determining that there would be minimal risk to your privacy. It is possible that a sponsor or their representatives, a data coordinating office, a contract research organization, will come to inspect your records. Even if those records are identifiable when inspected, the information leaving the institution will be stripped of direct identifiers. Additionally, the monitors, auditors, the IRB, the Food and Drug Administration will be granted direct access to your medical records for verification of the research procedures and data. By signing this document you are authorizing this access. We may publish the results of this research. However, we will keep your name and other identifying information confidential.

For how long will Mount Sinai be able to use or disclose your protected health information? Your authorization for use of your protected health information for this specific study does not expire.

Will you be able to access your records?

During your participation in this study, you will have access to your medical record and any study information that is part of that record. The investigator is not required to release to you research information that is not part of your medical record.

Do you need to give us permission to obtain, use or share your health information?

NO! If you decide not to let us obtain, use or share your health information you should not sign this form, and you will not be allowed to volunteer in the research study. If you do not sign, it will not affect your treatment, payment or enrollment in any health plans or affect your eligibility for benefits.

Can you change your mind?

IRB Approved

**Study ID #: HSM14-00530** **Form Version Date: February 11, 2015**

You may withdraw your permission for the use and disclosure of any of your protected information for research, but you must do so in writing to the Principal Investigator at the address on the first page. Even if you withdraw your permission, the Principal Investigator for the research study may still use your protected information that was already collected if that information is necessary to complete the study. Your health information may still be used or shared after you withdraw your authorization if you should have an adverse event (a bad effect) from being in the study. If you withdraw your permission to use your protected health information for research that means you will also be withdrawn from the research study, but standard medical care and any other benefits to which you are entitled will not be affected. You can also tell us you want to withdraw from the research study at any time without canceling the Authorization to use your data.

If you have not already received it, you will also be given the Mount Sinai Hospital - Mount Sinai School of Medicine Notice of Privacy Practices that contains more information about how Mount Sinai uses and discloses your protected health information.

It is important for you to understand that once information is disclosed to others outside Mount Sinai, the information may be re-disclosed and will no longer be covered by the federal privacy protection regulations. However, even if your information will no longer be protected by federal regulations, where possible, Mount Sinai has entered into agreements with those who will receive your information to continue to protect your confidentiality.

If as part of this research project your medical records are being reviewed, or a medical history is being taken, it is possible that HIV-related information may be revealed to the researchers. If that is the case, the information in the following box concerns you. If this research does not involve any review of medical records or questions about your medical history or conditions, then the following section may be ignored.

---

**Notice Concerning HIV-Related Information**

If you are authorizing the release of HIV-related information, you should be aware that the recipient(s) is (are) prohibited from re-disclosing any HIV-related information without your authorization unless permitted to do so under federal or state law. You also have a right to request a list of people who may receive or use your HIV-related information without authorization. If you experience discrimination because of the release or disclosure of HIV-related information, you may contact the New York State Division of Human Rights at (888) 392-3644 or the New York City Commission on Human Rights at (212) 306-5070. These agencies are responsible for protecting your rights.

---

IRB Approved

**Study ID #: HSM14-00530** **Form Version Date: February 11, 2015**

**Signature Block for Capable Adult**

Your signature below documents your permission to take part in this research and to the use and disclosure of your protected health information. A signed and dated copy will be given to you.

**DO NOT SIGN THIS FORM AFTER THIS DATE →** **6/30/2015**

Signature of subject ______________________ Date ____________

Printed name of subject ______________________ Time ____________

**Person Explaining Study and Obtaining Consent**

Signature of person obtaining consent ______________________ Date ____________

Printed name of person obtaining consent ______________________ Time ____________

***Witness Section: For use when a witness is required to observe the consent process,, document below (for example, subject is illiterate or visually impaired, or this accompanies a short form consent):***

*My signature below documents that the information in the consent document and any other written information was accurately explained to, and apparently understood by, the subject, and that consent was freely given by the subject.*

*Signature of witness to consent process* ______________________ *Date* ____________

*Printed name of person witnessing consent process* ______________________ Time ____________

**Document S2. HIV Consent.** ***Related to Figure 1.***

IRB Approved

GCO #: 13-1953

**Study ID #: HSM14-00530** **Form Version Date: February 11, 2015**

#### TITLE OF RESEARCH STUDY:

Title: **Drug Combination Signatures for Prediction and Mitigation of Toxicity**

#### PRINCIPAL INVESTIGATOR (HEAD RESEARCHER) NAME AND CONTACT INFORMATION:

Name: Ravi Iyengar, PhD

Physical Address: Icahn Medical Institute building, 12-70C

Mailing Address: One Gustave L. Levy Place
Box 1215
New York, NY 10029

#### WHAT IS A RESEARCH STUDY?

A research study is when scientists try to answer a question about something that we don't know enough about. Participating may not help you or others.

People volunteer to be in a research study. The decision about whether or not to take part is totally up to you. You can also agree to take part now and later change your mind. Whatever you decide is okay. It will not affect your ability to get medical care at Mount Sinai.

Someone will explain this research study to you. Feel free to ask all the questions you want before you decide. Any new information that develops during this research study that might make you change your mind about participating will be given to you promptly.

Basic information about this study will appear on the website http://www.ClinicalTrials.gov. There are a few reasons for this: the National Institutes of Health (NIH) encourages all researchers to post their research; some medical journals only accept articles if the research was posted on the website; and, for research studies the U.S. Food and Drug Administration (FDA) calls "applicable clinical trials" a description of this clinical trial will be available on http://www.ClinicalTrials.gov, as required by U.S. Law. This Web site will not include information that can identify you. At most, the Web site will include a summary of the results. You can search this Web site at any time.

#### PURPOSE OF THIS RESEARCH STUDY:

The purpose of this study is to understand adverse events caused by useful drugs in normal tissues of humans. For this we plan to obtain skin samples of healthy human subjects. Using this skin sample, we intend to establish fibroblast lines that will subsequently be reprogrammed to cells with the potential to give rise to any cell of the body and then differentiated to cells of the heart, liver and peripheral sensory (neurons) system for the purpose of the generation of molecular signatures for prediction and reduction of drug side effects caused by an specific drug and drug combinations.

Another purpose of this research project is to collect and store human samples (such as skin samples) and health information. Researchers can then use the stored materials in future studies. Through such studies, they hope to find new ways to detect, treat, and maybe even prevent or cure

IRB Approved

health problems. Some of these studies may be about how genes affect health, or how genes affect response to treatment. (Genes, which are made up of DNA, have the information needed to build and operate a human body.) Some of the studies may lead to new products, such as drugs or tests for diseases.

You may qualify to take part in this research study because your examination by the clinical research center indicates that you are a healthy person between the ages of 18-65.

Funds for conducting this research are provided by NIH.

#### LENGTH OF TIME AND NUMBER OF PEOPLE EXPECTED TO PARTICIPATE

Your participation in this research study is limited to a maximum of two visits. During the first visit to the clinical research center you will give informed consent and only then undergo a physical exam to establish your healthy status. Once your healthy status has been determined you will be contacted again to schedule an appointment during which we will obtain a skin sample. This visit will will take approximately 10 to 15 minutes to complete the skin biopsy. The tissue sample will be collected during a clinic visit arranged at your convenience and will require no extra time involvement after obtaining the tissue specimen.

The total number of people expected to take part in this research study is 30 individuals between 18-65 with no gender bias but about 30% ethnic minorities (African American, Asians and Hispanics).

#### DESCRIPTION OF WHAT'S INVOLVED:

If you agree to participate in this research study, the following information describes what may be involved.

During a visit to the clinical research center you will give informed consent prior to undergoing a physical exam to establish your healthy status. The exam will involve assessment of your clinically history, taking of vital signs and major system examinations (including pulmonary, cardiovascular, GI), an electrocardiogram (ECG; describe in the next paragraph) and having blood drawn (18 mL) to test heart, liver and sensory/touch system function parameters. Your blood will also be tested for HIV. HIV is the term used for the virus that produces HIV infection and may ultimately lead to AIDS. You must be told that you are being tested for HIV and give consent. You will sign an additional consent form prior to the HIV blood test. You have a right to know the results of the test. If you test positive you will be given more tests to confirm the results of the first test. If you are HIV positive, the study doctor will also give you a list of referrals for further information and counseling. Female subjects only will have to provide a urine sample to determine their pregnancy status in order to rule out any eventual problems from the lidocaine (for local anesthesia) and the skin punch biopsy. Specifically, the physical exam will involve first checking of weight, height, waist and hip circumference, heart rate, blood pressure, rate of breathing (breaths per minute) and peripheral oxygen saturation by pulse oximeter placed on a finger (this is a non-invasive measurement that takes only 10-20 seconds and is not painful). The doctor will then perform a complete physical examination that will include examination of the cardiovascular, respiratory, gastrointestinal and neurological systems. Particular aspects will involve listening to the heart and lungs with a stethoscope, palpation (feeling with the hands) of the abdomen, and examination of the sensori-motor system. A rectal exam will not be performed. This complete physical examination will take about 10-15 minutes.

IRB Approved

An ECG is the recording of the electrical activity of the heart captured over time by external electrodes attached to the skin. This involves placing 10 recording electrodes on the skin, with a gel-like substance contacting the skin (which can conduct the heart electricity). Six electrodes go across the left chest, while one electrode is attached to each limb (left and right arm and leg). For persons with a "hairy chest", shaving of some of the chest hair may be required to achieve good contact of the sensing electrodes and the skin. The entire process takes 5-10 minutes and is painless. Because an ECG is only recording the activity of the heart, there are no "electrical" or other sensations felt.
If you continue to qualify to take part in this study, you will have a skin sample taken at your convenience. If it is determined that you are not healthy and cannot continue in this study, you will be provided the results of the test by the study physician and be recommendation to see your personal physician. If you do not have a personal physician, we will refer you to the Mount Sinai Internal Medicine Associates.

After determination of your healthy status, a skin samples will be collected during a second visit arranged at your convenience. No further time involvement is required. We will obtain and save these samples and store the tissue in a bank with other samples that will be used for the research testing described as above and will be stored for a minimum of five years and for as long as deemed useful for research purposes. The skin sample will be assigned a 4-digit number (de-identification) for confidentiality during the research study.

The skin biopsy is called a punch biopsy and involves using a small 2 mm bore needle to obtain a sample of skin from either the forearm or behind the knee. The size of the sample taken will be approximately similar to the size of this typed letter "O", approximately 2 mm in size. A numbing anesthetic will be injected locally to numb the area before the sample is obtained. The area will then be covered with a small gauze or bandage. The risks involved are listed below. The punch biopsy will be obtained by a clinician who is a member of our research team.

We may wish to do future testing of your skin samples and derivatives, including but not limited to generated induced pluripotent stem cells and differentiated target cells such as heart, liver and peripheral nerve cells, based on potential future scientific developments and discoveries.

The researchers would like to ask your permission to keep specimens (like blood, tissue, hair, or any other body matter) collected or derived from you during this study to use them in future research studies. They would also like to know your wishes about how they might use your specimens in future research studies. You should also know that it is possible that products may someday be developed with the help of your specimens, and there are no plans to share any profits from such products with you.

***(1)** Will you allow the researchers to store your specimens to use in future research studies?*

*Yes________ No_________ If no, please stop here. If yes, please continue to the next question.*

**(2)** The researchers can keep your specimens stored in one of two different ways: one way will store your specimens in a way that it is linked to your identity (through the use of a code that can indicate

IRB Approved

the information came from you personally) and the other way will store your specimens anonymously (no one will know who the information is from). It will not be stored both ways, so you must choose one of these two options. Please note that if you choose to have your specimens stored anonymously, you will not be able to change your mind to ask for your specimens to be destroyed at a future date.

How would you like your specimens stored? Please initial **ONE** choice:
I would like my specimens stored with a link to my identity__________
I would like my specimens stored anonymously __________

**(3)** Do you give the researchers permission to **contact you** in the future to collect additional information about you, discuss how your specimens might be used, or to discuss possible participation in another research project? Please initial your choice:

*Yes*________ *No*_________

*(4) Do you give the researchers permission to keep the specimens indefinitely and use them for future studies that are* ***directly related*** *to the purpose of the current study? Please initial your choice:*

*Yes*________ *No*_________

*(5) Do you give the researchers permission to keep the specimens indefinitely and use them for future studies that are* ***not related*** *to the purpose of the current study (for example, a different area of research)? Please initial your choice:*

*Yes*________ *No*_________

**(a)** If the future research in a different area can be done without having to know that the specimens came from you personally, that will be done.

**(b)** If the future research in a different area requires that it is known specifically who the specimens came from, then one of the following will be done:

**(i)** If you allowed the researchers to contact you in the future, they will be able to contact you to explain why your specimen is needed and what will be done with it. Your permission will be asked to use your specimens in that research project.

**(ii)** If you do not give permission to be contacted in the future, or if it is found that contacting you is not practical, for example, because you have moved, your specimens may still be used. Either all links to your identity will be removed from the specimens, or an Institutional Review Board will be asked for permission to use the specimens linked to your identity. The Institutional Review Board (IRB) is a committee of doctors

IRB Approved

and scientists and non-scientists and people not associated with this hospital or medical school whose job it is to protect people who participate in research. The IRB can give permission for researchers to use and share health information connected to specimens that are linked to people's identities, but only if it determines that doing this will not be more than a minimal risk to people or their privacy.

**(6)** Do you give permission to have portions of the specimens **given to other researchers** at Mount Sinai or other institutions for use in research that is either related or not related to the purpose of this study? Please initial your choice:

*Yes*_________ *No*__________

As part of this study and do more powerful research (also in the future), it is helpful for researchers to share information they get from studying human samples. The scientific data collected in this study (but not your personal information such as name, address or social security number; see also below) will be deposited with the National Institutes of Health (an agency of the federal government) funded Data Coordination and Integration Center for the LINCS-BDK2 consortium, and possibly more scientific databases that may be maintained by Mount Sinai, by the federal government, or even by private companies, where it is stored along with information from other studies. Researchers can then study the combined information to learn even more about health and disease. A researcher who wants to study the information must adhere to rules set for by the Data Coordination and Integration Center for the LINCS-BDK2 consortium and conjunction with National Institutes of Health guidelines. Your name and other information that could directly identify you (such as address or social security number) will never be placed into a scientific database. However, because your genetic information is unique to you, there is a small chance that someone could trace it back to you. The risk of this happening is very small, but may grow in the future. Researchers will always have a duty to protect your privacy and to keep your information confidential.

In general, results from this research project will not be shared with you or put into your medical records. In some situations, the results (including genome sequencing results) might be important to your health or medical care. If this occurs, we will contact you to see if you want to learn more and if so refer you to your personal physician.

*No tests other than those authorized will be performed on the skin sample.*

#### YOUR RESPONSIBILITIES IF YOU TAKE PART IN THIS RESEARCH:

If you decide to take part in this research study you will be responsible for undergoing a physical exam, an electrocardiogram and provide a blood sample for lab tests to establish your healthy status and only if such is confirmed to provide a skin sample, which will be obtained at a separate visit arranged at your convenience.

#### COSTS OR PAYMENTS THAT MAY RESULT FROM PARTICIPATION:

IRB Approved

If you agree to take part in this research study, after completion of the skin punch biopsy, we will pay you $150 in cash as reimbursement for your time and effort.

Tax law may require the Mount Sinai Finance Department to report the amount of payment you receive from Mount Sinai to the Internal Revenue Service (IRS) or other agencies, as applicable. Generally this reporting would take place if you receive payments that equal $600 or more from Mount Sinai in a calendar year. You would be responsible for the payment of any tax that may be due.

#### POSSIBLE BENEFITS:

You are not expected to get any benefit from taking part in this research study. Others may not benefit either. However, possible benefits include gaining valuable knowledge about specific drug and/or drug combinations and why they may be toxic on a molecular level to cells of the heart, liver and peripheral sensory system (neurons). Even if such knowledge is gained it will not be shared with you.

#### REASONABLY FORESEEABLE RISKS AND DISCOMFORTS:

Possible risks and discomforts include those associated with obtaining the skin biopsy, which are bleeding, localized collection of blood outside the blood vessels, infection, and pain and/or physical discomfort at the time of the sampling. The sampling will be done during a routine clinic visit. Risks of the local anesthetic Lidocaine, used for skin biopsy, are pain at the site of sampling/injection. Additionally, lidocaine can pass very quickly through the placenta and reports exist of negative side effects on infants born to mothers using the anesthetic. Several lidocaine studies with human case reports have showed no negative side effects, pregnancy complications or birth defects. In addition, there is minimal risk for an allergic reaction to the local anesthetic. However, allergic reactions are immediate and since you will be watched by the study doctor during the procedure, the doctor can will watch for and treat any allergic reaction that may occur.

Group Risks □
Although we will not give researchers your name, we will give them basic information such as your race, ethnic group, and sex. This information helps researchers learn whether the factors that lead to health problems are the same in different groups of people. It is possible that such findings could one day help people of the same race, ethnic group, or sex as you. However, they could also be used to support harmful stereotypes or even promote discrimination. □

Privacy Risks
There always exists the potential for loss of private information; however, there are procedures in place to minimize this risk.

Your privacy is very important to us and we will use many safety measures to protect your privacy. However, in spite of all of the safety measures that we will use, we cannot guarantee that your identity will never become known. Although your genetic information is unique to you, you do share some genetic information with your children, parents, brothers, sisters, and other blood relatives. Consequently, it may be possible that genetic information from them could be used to help identify

IRB Approved

you. Similarly, it may be possible that genetic information from you could be used to help identify them. □

As part of this study and do more powerful research (also in the future), it is helpful for researchers to share information they get from studying human samples. The scientific data collected in this study (but not your personal information such as name, address or social security number; see also below) will be deposited with the National Institutes of Health funded Data Coordination and Integration Center for the LINCS-BDK2 consortium, and possibly more scientific databases, where it is stored along with information from other studies. Researchers can then study the combined information to learn even more about health and disease. A researcher who wants to study the information must adhere to rules set for by the Data Coordination and Integration Center for the LINCS-BDK2 consortium and conjunction with National Institutes of Health guidelines.

Your name and other information that could directly identify you (such as address or social security number) will never be placed into a scientific database. However, because your genetic information is unique to you, there is a small chance that someone could trace it back to you. The risk of this happening is very small, but may grow in the future. Since the database includes genetic information, a break in security may also pose a potential risk to blood relatives as well as yourself. For example, it could be used to make it harder for you (or a relative) to get or keep a job or insurance. If your private information was misused it is possible you would also experience other harms, such as stress, anxiety, stigmatization, or embarrassment from revealing information about your family relationships, ethnic heritage, or health conditions.

People may develop ways in the future that would allow someone to link your genetic or medical information in our databases back to you. For example, someone could compare information in our databases with information from you (or a blood relative) in another database and be able to identify you (or your blood relative). It also is possible that there could be violations to the security of the computer systems used to store the codes linking your genetic and medical information to you. □□

Since some genetic variations can help to predict the future health problems of you and your relatives, this information might be of interest to health providers, life insurance companies, and others. Patterns of genetic variation also can be used by law enforcement agencies to identify a person or his/her blood relatives. Therefore, your genetic information potentially could be used in ways that could cause you or your family distress, such as by revealing that you (or a blood relative) carry a genetic disease. □

There is a Federal law called the Genetic Information Nondiscrimination Act (GINA). In general, this law makes it illegal for health insurance companies, group health plans, and most large employers to discriminate against you based on your genetic information. However, it does not protect you against discrimination by companies that sell life insurance, disability insurance, or long-term care insurance.

There also may be other privacy risks that we have not foreseen.

**OTHER POSSIBLE OPTIONS TO CONSIDER:**

IRB Approved

You may decide not to take part in this research study without any penalty. The choice is totally up to you.

**IN CASE OF INJURY DURING THIS RESEARCH STUDY:**

If you believe that you have suffered an injury related to this research as a participant in this study, you should contact the Principal Investigator.

**ENDING PARTICIPATION IN THE RESEARCH STUDY:**

You may stop taking part in this research study at any time without any penalty. This will not affect your ability to receive medical care at Mount Sinai or to receive any benefits to which you are otherwise entitled.

If you decide to stop being in the research study, please contact the Principal Investigator or the research staff.

You may also withdraw your permission for the use and disclosure of any of your protected information for research, but you must do so in writing to the Principal Investigator at the address on the first page. Even if you withdraw your permission, the Principal Investigator for the research study may still use the information that was already collected if that information is necessary to complete the research study. Your health information may still be used or shared after you withdraw your authorization if you should have an adverse event (a bad effect) from participating in the research study.

Withdrawal without your consent: The study doctor, the sponsor or the institution may stop your involvement in this research study at any time without your consent. This may be because the research study is being stopped, the instructions of the study team have not been followed, the investigator believes it is in your best interest, or for any other reason. If specimens or data have been stored as part of the research study, they too can be destroyed without your consent.

If you chose to end your participation in this study, all specimens, i.e. fibroblasts derived from the biopsy (skin sample) as well as the induced pluripotent stem cells generated from those fibroblasts can easily be discarded/destroyed.

**CONTACT PERSON(S):**

If you have any questions, concerns, or complaints at any time about this research, or you think the research has hurt you, please contact the office of the research team and/or the Principal Investigator at phone number (212)-659-1707 or Christoph Schaniel, Ph.D., at (212)-659-8276.

If you experience an emergency during your participation in this research, contact Dr. Jason Kovacic at (212)-241-7300.

This research has been reviewed and approved by an Institutional Review Board. You may reach a representative of the Program for Protection of Human Subjects at the Icahn School of Medicine at Mount Sinai at telephone number (212) 824-8200 during standard work hours for any of the following reasons:

- Your questions, concerns, or complaints are not being answered by the research team.

IRB Approved

- You cannot reach the research team.
- You are not comfortable talking to the research team.
- You have questions about your rights as a research subject.

You want to get information or provide input about this research.

**DISCLOSURE OF FINANCIAL INTERESTS:**

None.

**MAINTAINING CONFIDENTIALITY – HIPAA AUTHORIZATION:**

As you take part in this research project it will be necessary for the research team and others to use and share some of your private protected health information. Consistent with the federal Health Insurance Portability and Accountability Act (HIPAA), we are asking your permission to receive, use and share that information.

What protected health information is collected and used in this study, and might also be disclosed (shared) with others?

As part of this research project, the researchers will collect your name, address, telephone number, and medical record number at Mount Sinai Hospital.

The researchers will also get information from your medical record from Mount Sinai Hospital.

During the study the researchers will gather information by:

- taking a medical history (includes current and past medications or therapies, illnesses, conditions or symptoms, family medical history, allergies, etc.)
- completing the tests, procedures, questionnaires and interviews explained in the description section of this consent.

Why is your protected health information being used?

Your personal contact information is important to be able to contact you during the study. Your health information and the results of any tests and procedures being collected as part of this research study will be used for the purpose of this study as explained earlier in this consent form. The results of this study could be published or presented at scientific meetings, lectures, or other events, but would not include any information that would let others know who you are, unless you give separate permission to do so.

The research team and other authorized members of The Mount Sinai Hospital and Mount Sinai School of Medicine (together, "Mount Sinai") workforce may use and share your information to ensure that the research meets legal, institutional or accreditation requirements. For example, the Mount Sinai School of Medicine Program for the Protection of Human Subjects is responsible for overseeing research on human subjects, and may need to see your information. If you receive any payments for taking part in this study, the Mount Sinai Medical Center Finance Department may need your name, address, social security number, payment amount, and related information for tax reporting purposes.

If the research team uncovers abuse, neglect, or reportable diseases, this information may be disclosed to appropriate authorities.

Who, outside Mount Sinai, might receive your protected health information? None.

As part of the study, the Principal Investigator, study team and others in the Mount Sinai workforce may disclose your protected health information, including the results of the research study tests and procedures, to the following people or organizations:

- The United States Department of Health and Human Services and the Office of Human Research Protection.
- The sponsoring government agency and/or their representative who need to confirm the accuracy of the results submitted to the government or the use of government funds: The National Institutes of Health.

(It is possible that there may be changes to the list during this research study; you may request an up-to-date list at any time by contacting the Principal Investigator.)

In all disclosures outside of Mount Sinai, you will not be identified by name, social security number, address, telephone number or any other direct personal identifier unless disclosure of the direct identifier is required by law. Some records and information disclosed may be identified with a unique code number. The Principal Investigator will ensure that the key to the code will be kept in a locked file, or will be securely stored electronically. The code will not be used to link the information back to you without your permission, unless the law requires it, or rarely if the Institutional Review Board allows it after determining that there would be minimal risk to your privacy. It is possible that a sponsor or their representatives, a data coordinating office, a contract research organization, will come to inspect your records. Even if those records are identifiable when inspected, the information leaving the institution will be stripped of direct identifiers. Additionally, the monitors, auditors, the IRB, the Food and Drug Administration will be granted direct access to your medical records for verification of the research procedures and data. By signing this document you are authorizing this access. We may publish the results of this research. However, we will keep your name and other identifying information confidential.

For how long will Mount Sinai be able to use or disclose your protected health information? Your authorization for use of your protected health information for this specific study does not expire.

Will you be able to access your records?

During your participation in this study, you will have access to your medical record and any study information that is part of that record. The investigator is not required to release to you research information that is not part of your medical record.

Do you need to give us permission to obtain, use or share your health information?

NO! If you decide not to let us obtain, use or share your health information you should not sign this form, and you will not be allowed to volunteer in the research study. If you do not sign, it will not affect your treatment, payment or enrollment in any health plans or affect your eligibility for benefits.

Can you change your mind?

IRB Approved

You may withdraw your permission for the use and disclosure of any of your protected information for research, but you must do so in writing to the Principal Investigator at the address on the first page. Even if you withdraw your permission, the Principal Investigator for the research study may still use your protected information that was already collected if that information is necessary to complete the study. Your health information may still be used or shared after you withdraw your authorization if you should have an adverse event (a bad effect) from being in the study. If you withdraw your permission to use your protected health information for research that means you will also be withdrawn from the research study, but standard medical care and any other benefits to which you are entitled will not be affected. You can also tell us you want to withdraw from the research study at any time without canceling the Authorization to use your data.

If you have not already received it, you will also be given the Mount Sinai Hospital - Mount Sinai School of Medicine Notice of Privacy Practices that contains more information about how Mount Sinai uses and discloses your protected health information.

It is important for you to understand that once information is disclosed to others outside Mount Sinai, the information may be re-disclosed and will no longer be covered by the federal privacy protection regulations. However, even if your information will no longer be protected by federal regulations, where possible, Mount Sinai has entered into agreements with those who will receive your information to continue to protect your confidentiality.

If as part of this research project your medical records are being reviewed, or a medical history is being taken, it is possible that HIV-related information may be revealed to the researchers. If that is the case, the information in the following box concerns you. If this research does not involve any review of medical records or questions about your medical history or conditions, then the following section may be ignored.

---

**Notice Concerning HIV-Related Information**

If you are authorizing the release of HIV-related information, you should be aware that the recipient(s) is (are) prohibited from re-disclosing any HIV-related information without your authorization unless permitted to do so under federal or state law. You also have a right to request a list of people who may receive or use your HIV-related information without authorization. If you experience discrimination because of the release or disclosure of HIV-related information, you may contact the New York State Division of Human Rights at (888) 392-3644 or the New York City Commission on Human Rights at (212) 306-5070. These agencies are responsible for protecting your rights.

---

IRB Approved

**Signature Block for Capable Adult**

Your signature below documents your permission to take part in this research and to the use and disclosure of your protected health information. A signed and dated copy will be given to you.

**DO NOT SIGN THIS FORM AFTER THIS DATE → 6/30/2015**

Signature of subject ______________________ Date ________

Printed name of subject ______________________ Time ________

**Person Explaining Study and Obtaining Consent**

Signature of person obtaining consent ______________________ Date ________

Printed name of person obtaining consent ______________________ Time ________

***Witness Section: For use when a witness is required to observe the consent process, document below (for example, subject is illiterate or visually impaired, or this accompanies a short form consent):***

*My signature below documents that the information in the consent document and any other written information was accurately explained to, and apparently understood by, the subject, and that consent was freely given by the subject.*

*Signature of witness to consent process* ______________________ *Date* ________

*Printed name of person witnessing consent process* ______________________ Time ________

IRB Approved

GCO #: 13-1953
**Study ID #:** HSM14-00530 **Form Version Date: February 11, 2015**

#### TITLE OF RESEARCH STUDY:

Title: **Drug Combination Signatures for Prediction and Mitigation of Toxicity**

#### PRINCIPAL INVESTIGATOR (HEAD RESEARCHER) NAME AND CONTACT INFORMATION:

Name: Ravi Iyengar

Physical Address: Icahn Medical Institute, building 12-70C

Mailing Address: 1 Gustave L Levy Place, Box 1215, NY, NY 10029]

#### INFORMATION ABOUT HIV ANTIBODY TESTING:

You have been asked to participate in a research study, for which you have signed a separate consent. As indicated on that consent, part of your participation in that study will involve undergoing an HIV antibody test. This consent is specifically for the HIV antibody test. The HIV antibody test is a blood test used to ascertain whether you have antibodies to the Human Immunodeficiency Virus (HIV), the virus which causes Acquired Immunodeficiency Syndrome (AIDS). Less than one teaspoon of blood will be drawn from a vein in your arm using a needle. This may cause some discomfort and you may develop a black and blue mark. It takes approximately one to two weeks between the time your blood is drawn and the time you are notified of the results.

Both before and after your blood is tested, you will receive counseling from trained HIV counselors involved in this research project about the implications of negative and positive results, how to prevent future transmission, and the options available to you. Your partners may be notified of the results of this test and urged to undergo testing as well. If you do not want to tell them, or you will not tell them, your doctor or local health official can inform them that a partner of theirs has been tested and what the results of the test were, but only if the doctor feels that telling them is medically appropriate. You will not incur any costs nor receive any payment for participating in this part of the study.

A positive HIV antibody test means that your body is making antibodies to HIV but it does not mean that you will necessarily develop AIDS in the future. A negative test means that you are probably not infected; however, it is possible that you may be infected but that your body has not produced antibodies to HIV. If your results are negative and you have been exposed to HIV recently, you should be retested in a few months to make sure you are not infected.

There are several possible benefits to taking the HIV antibody test. If your test results are negative, you can learn how to avoid becoming infected in the future. If your results are positive, you can learn how to avoid infecting other individuals, and if you are pregnant, or are thinking about having children, you can learn how being HIV positive will affect your decision to have children. Additionally, we can offer you enrollment in a wide variety of research projects for the treatment of AIDS or refer you to a doctor for non-experimental treatment.

This is a voluntary procedure, and all results, either positive or negative, are confidential. Under New York State law, information about your HIV antibody test can only be released to people who you designate by signing a release form, or to those people listed below:

a) You (or a person authorized by law who consented to the test for you);

IRB Approved

b) To a health care facility (such as a hospital, blood bank, or clinical laboratory) or a health care provider (such as a physician, nurse, or mental health counselor) providing care to you or your child, and anyone working for such a facility or provider who reasonably needs the information to supervise, monitor or administer health care;
c) To a person whom your doctor believes is at significant risk for HIV infection, if you do not notify that person after being counseled to do so;
d) To a committee or organization responsible for reviewing or monitoring a health facility;
e) To a federal, state, county, or local health officer when state or federal law requires disclosure;
f) To a government agency, when the agency needs the information to supervise, monitor, or administer a health or social service;
g) To an authorized foster care or adoption agency;
h) To insurance companies and other third party payers such as Medicaid necessary for the payment of services to you;
i) To any person whom a court orders disclosure under limited circumstances set forth by law. Except in an emergency situation, advance notice and an opportunity to oppose the release of such information would be given to you;
j) To the Division of Parole, the Division of Probation, the Commission of Correction, or a medical director of a local correctional facility, as permitted by HIV confidentiality regulations of such organization.
k) By a physician to someone who may consent to health care for you if you have been counseled and won't inform such person and disclosure is medically necessary to provide timely care and treatment. Disclosure must not be against your best interest.

If you do not want anybody to know your tests results or that you have been tested, you can go to an anonymous test site. This is a place where you can have your blood tested and receive counseling without having to tell anybody your name or address. You can find the nearest anonymous test site by calling the AIDS Hotline at **1-(800)-541-2437**.

If your results are positive, you should be very careful who you disclose this information to. Some HIV positive people have been discriminated against by landlords, employers, and the like. If you believe you have been discriminated against, you should call the New York State Division of Human Rights at (888) 392-3644 or the New York City Commission on Human Rights at (212) 306-5070. If you have any further questions regarding AIDS or HIV antibody testing, you can contact the New York State Department of Health AIDS Hotline at 1-(800) -TALK-HIV / 1-(800)-825-5448.

IRB Approved

GCO #: 13-1953

**Study ID #:** HSM14-00530 **Form Version Date: February 11, 2015**

**Signature Block for Capable Adult**

Your signature below documents your permission for HIV Antibody Testing, pre-test counseling, post-test counseling and for a blood draw of less than one teaspoon of blood, one time from me. A signed and dated copy will be given to you.

**DO NOT SIGN THIS FORM AFTER THIS DATE →** **6/30/2015**

Signature of subject ______ Date and Time ______

Printed name of subject ______

**Person Explaining Study and Obtaining Consent**

Signature of person obtaining consent ______ Date and Time ______

Printed name of person obtaining consent ______

***If the individual cannot read, a witness is required to observe the consent process and document below:***

*My signature below documents that the information in the consent document and any other written information was accurately explained to, and apparently understood by, the subject, and that consent was freely given by the subject.*

*Signature of witness to consent process* ______ *Date and Time* ______

*Printed name of person witnessing consent process* ______

### Drug Combination Signatures for Prediction and Mitigation of Toxicity

#### ENROLLMENT FORM

**Pt. Initials ___ ___ ___ Pt. ID Number___ ___ ___ Date:__/______/______(mm/dd/yyyy)**

| **INCLUSION CRITERIA** | **Yes** | **No** |
|---|---|---|
| Patient ≥18 and ≤65 years of age and freely willing to participate | ☐ | ☐ |
| Signed written Informed Consent | ☐ | ☐ |
| **EXCLUSION CRITERIA** | | |
| Abnormal EKG | ☐ | ☐ |
| Any family history of any cardiovascular disorder, excluding hypertension, in any first or second degree relative(s), at age < 50. This includes stroke, myocardial infarction, angina, peripheral vascular disease, aortic dissection, aortic aneurysm, vasculitis, vasculopathy | ☐ | ☐ |
| Any family history of non-ischemic cardiomyopathy in any first or second degree relative(s), at any age | ☐ | ☐ |
| More than 2 pack-year of lifetime smoking | ☐ | ☐ |
| Positive for: HIV, Hepatitis B, Hepatitis C | ☐ | ☐ |
| Pts. who are currently participating in another investigational drug/device study | ☐ | ☐ |
| Patients with liver disease | ☐ | ☐ |
| Diabetes | | |
| Pregnant women and/or nursing mothers | ☐ | ☐ |
| Patients having undergone heart transplantation or any other organ transplantation. | ☐ | ☐ |
| Personal or family history of neuropathy, at any age in a first or second degree relative, with the exception of diabetic neuropathy in another family member over the age of 60 at first diagnosis | ☐ | ☐ |
| Personal or family history of myopathy, at any age in a first or second degree relative | ☐ | ☐ |
| Body Mass Index ≥ 30 kg/$m^2$ | ☐ | ☐ |
| Creatinine ≥ 1.3 mg/dL or known renal disease | ☐ | ☐ |
| Abnormal Chem14 panel, renal function or liver function studies | ☐ | ☐ |
| Abnormal BNP (Brain natriuretic peptide) | ☐ | ☐ |
| Abnormal hemoglobin, platelet count or white cell count | ☐ | ☐ |
| Other disorders, clinically manifest or detected by screening labs, that are associated with cardiovascular disease, including but not limited to hemochromatosis, hyperthyroidism or hypothyroidism. | ☐ | ☐ |
| Active autoimmune disease | ☐ | ☐ |
| Taking any medications other than occasional aspirin, occasional NSAIDS, occasional acetaminophen or the oral contraceptive pill. | ☐ | ☐ |
| Asthma requiring any regularly inhaled therapy | ☐ | ☐ |
| Epilepsy requiring any form of ongoing therapy (medication or other) | ☐ | ☐ |

**DEMOGRAPHICS**

Date of Birth: _____/____/_________(mm/dd/yyyy) Gender: ☐ Male ☐ Female

Race: ☐ Caucasian ☐ Hispanic
☐ Asian ☐ Other (specify) ______________
☐ African-American

**Social History:**

Tobacco Use:

☐ Never
☐ Quit; total duration ________________
☐ Current; total duration ________________

**Other Clinical History:**

____________________________________________________________
____________________________________________________________
____________________________________________________________
____________________________________________________________
____________________________________________________________
____________________________________________________________
____________________________________________________________
____________________________________________________________
____________________________________________________________
____________________________________________________________
____________________________________________________________
____________________________________________________________
____________________________________________________________
____________________________________________________________
____________________________________________________________
____________________________________________________________
____________________________________________________________
____________________________________________________________
____________________________________________________________
____________________________________________________________
____________________________________________________________
____________________________________________________________

**EXAMINATION:**

Weight: ________ kg/lbs ; Height: ________cm/inches; Waist: ________cm/inches; Hip: ________cm/inches

Heart Rate: _______ /min Blood Pressure: _____/____mm of Hg

Respiratory Rate: _______ /min

Oxygen saturation (on room air): _______ %

**Cardiovascular Exam:**
JVP:
Carotid bruits:
Apex beat:
Auscultatory findings:
Palpable AAA:
Pedal pulses:
Pedal edema:

**Respiratory Exam:**
Clubbing:
Upper airway:
Trachea midline:
Chest expansion:
Auscultation:

**GI/Abdominal Exam:**
Sclerae (jaundice):
Spider naevi:
Abdominal distension:
General palpation:
Liver:
Spleen:
Auscultation:

**Neurological Exam:**
Sit to stand without using upper limbs:
Screening cranial nerve exam:
Screening upper limb (light touch, tone, strength flexion and extension):
Screening lower limb (light touch, tone, strength flexion and extension):
Neurofilament (great toe):

**LABORATORY RESULTS:**

**Date of collection:____/____/_______ (mm/dd/yyyy) Time: ____:____(24-hr clock)**

| | | |
|---|---|---|
| Glucose | | 65-139 mg/dL |
| Sodium | | 135-145 MEQ/L |
| Potassium | | 3.5 – 5.0 MEQ/L |
| Chloride | | 96-108 MEQ/L |
| CO2 total | | 22 – 32 MEQ/L |
| Urea Nitrogen | | 10 – 30 mg/dL |
| Creatinine | | 0.6 – 1.4 mg/dL |
| eGFR African Am | | >60 ml/min/1.73m2 |
| eGFR non African Am | | >60 ml/min/1.73m2 |
| Calcium | | 8.5 – 10.5 mg/dL |
| ALT | | 1 – 53 U/L |
| AST | | 1 – 50 U/L |
| Bilirubin total | | 0.1 – 1.2 mg/dL |
| Alk Phos | | 30 – 110 U/L |

| | | |
|---|---|---|
| Albumin | | 3.5 – 4.9 g/dL |
| Protein total | | 6.0 – 8.3 g/dL |

**CBC & Platelets Differential:**

| | | |
|---|---|---|
| WHITE BLOOD CELL | | 4.5-11.0 / x10 3/uL |
| RED BLOOD CELL | | 4.50-6.00 / x10 6/uL |
| HEMOGLOBIN | | 13.9-16.3 / G/DL |
| HEMATOCRIT | | 42.0-52.0 / % |
| MEAN CORP. VOLUME | | 80.0-98.0 / FL |
| MEAN CORP. HGB | | 27.0-32.0 / PG |
| MEAN CORP. HGB CONC. | | 32.0-35.0 / G/DL |
| RED DISTRIB. WIDTH | | 11.5-15.0 / % |
| PLATELET | | 150-450 / x10 3/uL |
| MEAN PLT VOLUME | | 7.4-12.0 / FL |
| NEUTROPHIL % | | 40.0-78.0 / % |
| LYMPHOCYTE % | | 15.0-50.0 / % |
| MONOCYTE % | | 2.0-11.0 / % |
| EOSINOPHIL % | | 0.0-5.0 / % |
| BASOPHIL % | | 0.0-1.0 / % |
| NEUTROPHIL # | | 1.9-8.0 / x10 3/uL |
| LYMPHOCYTE # | | 1.0-4.5 / x10 3/uL |
| MONOCYTE # | | 0.2-1.0 / x10 3/uL |
| EOSINOPHIL # | | 0.0-0.6 / x10 3/uL |
| BASOPHIL # | | 0.0-0.2 / x10 3/uL |
| NUCLEATE RBC% | | 0.0-0.0 / % |
| NRBC# | | 0.0-0.0 / x10 3/uL |

**Heart Failure, endocrine and other:**

| | | |
|---|---|---|
| BNP | | 0 – 100 pg/ml |
| Ferritin | | 30 – 400 ng/ml |
| Transferrin Sat | | 15 – 50% |
| HBA1C | | 4.0 – 6.0% |
| TSH | | 0.34 – 5.6 uIU/Ml |

**Infectious:**

| |
|---|
| HIV |
| HepC |
| HebB sAG |

#### Drug Combination Signatures for Prediction and Mitigation of Toxicity

| |
|---|
| HepB sAB |
| HepB cAB |

**Other Labs:**

**Date of collection:____/____/_______ (mm/dd/yyyy) Time:______________(24-hr clock)**

**Pregnancy Test:** ☐ Positive ☐ Negative ☐ Not Applicable [If male/Post menopausal/sterile female]

**ECG Date:___/____/______(mm/dd/yyyy) Time: _____:______(24-hr clock)**

ECGcomments:______________________________________________________________

Clinician Name: ____________________________

Clinician Signature: _________________________ Date: _____/_______/________
**(mm/dd/yyyy)**

Investigator's Signature: ________________________________ Date: _____/_______/________
**(mm/dd/yyyy)**

#### Supplemental Experimental Procedures

##### Karyotyping

G-banded karyotype analysis was performed at WiCell (Madison, WI) or the Tumor CytoGenomics Laboratory at the Icahn School of Medicine at Mount Sinai according to standard operating procedures. In general, at least 10 metaphases per iPSC clone were analyzed.

##### Short Tandem Repeat (STR) analysis

Genomic DNA was extracted from parental fibroblasts and derived iPSC clones using the PureLink™ Genomic DNA Mini Kit (Thermo Fisher Scientific, K1820-01) according to the manufacturer's instruction. STR polymorphism analysis for 15 loci plus Amelogenin (PowerPlex® 16 System, Promega DC6531) was performed using standard procedures at and interpreted by WiCell (Madison, WI).

##### Immunocytochemistry

For immunostaining of pluripotency markers, iPSCs were cultured in Matrigel-coated wells of 24-well plates (3 wells per clone) and when 50-75% confluent washed with PBS (Thermo Fisher Scientific, 14190250). For tri-lineage differentiation, iPSCs were differentiated in Matrigel-coated wells of 24-well plates (3 wells per clone) using the StemDiff Trilineage Differentiation kit (Stemcell Technologies; 05230) according to the manufacturer's instruction. Cells were fixed with 4% PFA (Electron Microscopy Sciences, 15710-S) in PBS, washed with PBS plus 0.1%Triton-X100 (Millipore Sigma, T8787) (PBS-T), washed 3 times with PBS and incubated with blocking buffer consisting of PBS-T supplemented with 10% donkey serum (Jackson ImmunoResearch, 017-000-121) and 0.25% BSA (Thermo Fisher Scientific, 15260037). Following blocking, primary antibodies used for detection of pluripotency markers were anti-OCT4 (Biovision, 3576, RRID:AB_2167563) plus anti-NANOG (Biotechne-R&D Systems, AF1997, RRID:AB_355097) in well 1, anti-SOX2 (Santa Cruz Biotechnology, sc-365823, RRID:AB_10842165) and anti-TRA-1-60 DyLight 650 (Thermo Fisher Scientific, MA1-023-D650, RRID:AB_2536702) for well 2; and anti-SSEA-4 PE (Biotechne-R&D Systems, FAB1435P) well 3, all diluted at 1:100 in blocking buffer overnight at 4˚C. Following primary antibody staining, cells were washed 3 times with PBS-T and stained with donkey polyclonal anti-rabbit IgG (H+L) Alexa Fluor 594 (Thermo Fisher Scientific, A-21207) plus donkey polyclonal anti-goat IgG (H+L) Alexa Fluor 488 secondary antibodies (Thermo Fisher Scientific, A-11055) diluted 1:1,000 in PBS-T in well 1, and donkey polyclonal anti-mouse IgG (H+L) Alexa Fluor 488 (Thermo Fisher Scientific, A-21202) in well 2. The primary antibodies used for detection of tri-lineage differentiation were anti-SOX17 (Biotechne-R&D Systems, AF1924, RRID:AB_355060) for endoderm in well 1, anti-Brachyury (Biotechne-R&D Systems, AF2085, RRID:AB_2200235) for mesoderm in well 2, and anti-NESTIN (Thermo Fisher Scientific-eBioscience, 14-9843-82, RRID:AB_1548837) for (neuro)ectoderm in well 3, all diluted at 1:25 in blocking buffer overnight at 4˚C. Following primary antibody staining, cells were washed 3 times with PBS-T and stained with donkey anti-goat Alexa Fluor 488 (Thermo Fisher Scientific, A-11055) in well 1 and well 2, and donkey anti-mouse Alexa Fluor 488 (Thermo Fisher Scientific, A-21202) in well 3, diluted 1:1,000 in PBS-T. Following secondary antibody staining, cells were washed 3 times with PBS and nuclei visualized by 4′,6-diamidino-2-phenylindole (DAPI; Thermo Fisher Scientific, 3570). Cells were imaged with an EVOS inverted fluorescence microscope (Thermo Fisher Scientific, 12-563-462). μ

##### RNA isolation, library preparation and RNA-seq

RNA from duplicate samples for each of the 40 iPSC lines or from various replicates (5-12) of cardiomyocyte differentiations (2-3) from 6 select hiPSC lines was isolated using TRIzol Reagent (Thermo Fisher Scientific, 15596018) according to the manufacturer's instruction. RNA was quantified using Qubit RNA BR Assay kit (Thermo Fisher Scientific, Q10211) on a Qubit 2.0 Fluorometer (Thermo Fisher Scientific, Q32866). RNA integrity was determined using the RNA 6000 Nano kit (Agilent, 5067-1511) on a 2100 Bioanalyzer Instrument (Agilent, G2939BA). Only RNA samples with an RNA Integrity Number (RIN) of >8 (our samples commonly had a RIN of 9-10) were used for library preparation. Library preparation was done using the TruSeq Stranded mRNA Library Prep (96 Samples) (Illumina, 20020595) on a minimum of 100 ng total RNA (ideally 300- 500 ng) with TruSeq RNA adapter plate (96 plex) (Illumina, 15016427) and IDT for Illumina-TruSeq RNA UD Indexes (96 Indexes, 96 Samples) (Illumina, 20022371) according to Illumina's TruSeq Stranded mRNA reference guide. The prepared libraries were quantified on a Qubit 2.0 Fluorometer using the Qubit dsDNA HS Assay Kit (Thermo Fisher Scientific, Q32851) and library integrity checked on a 2100 Bioanalyzer Instrument using the Agilent DNA 1000 Kit (Agilent, 5067-1505). Only libraries with a library peak around 250-300 bp and no to very low adapter contamination were sequenced.

Single cell (sc) RNA-seq of ventricular cardiomyocytes differentiated from 4 hiPSC lines was performed using the Chromium platform (10x Genomics) according to the manufacturer's instruction. An aliquot of the amplified cDNAs was run on a 2100 Bioanalyzer using the High Sensitivity DNA kit (Agilent, 5067-4626). cDNAs needed to have a size distribution of 300-10,000 bp to be considered for library generation. Integrity of scRNA-seq libraries was assessed on a 2100 Bioanalyzer Instrument using the Agilent DNA 1000 Kit (Agilent, 5067-1505). scRNA-seq libraries needed to be in the size range of 300-1,000 bp with the peak around 400 bp without visible adapter dimers to be sequenced.

Sequencing of the 80 hiPSC, the 63 hiPSC-derived cardiomyocyte (from 6 hiPSC lines), and 4 single-cell (sc) hiPSC-derived cardiomyocyte RNA libraries was performed at the Genomic Core at the Icahn School of Medicine at Mount Sinai. For the bulk RNA-seq, pooled libraries were clustered in either a S1 flowcell on a full NovaSeq 6000 (Illumina, 20012850) or in a Mid flowcell on a NextSeq 550 (Illumina, SY-420-1002) according to manufacturer's guidance. The samples were then sequenced using a 1x100bp Single End configuration, and 20 million read pairs were generated for each sample. scRNA-seq was performed in *an S4 flowcell on a full*

*NovaSeq 6000 (Illumina) according to manufacturer's guidance, using a 2x100bp Paired End configuration, targeting at least 50,000 read pairs per cell.*

**RNA-seq data analysis**

For the bulk RNA-seq data, raw base call data (.bcl files) generated from the run were converted into FASTQ sequence files and de-multiplexed into individual samples using Illumina's bcl2fastq 2.20 software. For sequence alignment, the RNA-seq FASTQ file of each sample was aligned to the reference human genome library of the GRCh38 (hg38) version (downloaded from the UCSC Genome Browser at http://hgdownload.soe.ucsc.edu/downloads.html) using the STAR aligner (version 2.5.3a). Subsequently, the aligned sequence reads in the alignment BAM file for each sample was looked up in the human genome annotation using the featureCounts program from Subread (version 1.6.2) to sum up the counts of all aligned sequence reads that can be uniquely mapped to each annotated reference gene for the sample. As a result, a list containing such read counts of all annotated genes is obtained for each sample and a sum of these read counts over all annotated genes determined the percentage of uniquely mapped reads of each sample (Table S4). In average, the percentage reads mapped to transcriptome (mapping rate) was 69% for the 80 hiPSC samples and 67% for the 63 hiPSC-derived cardiomyocyte samples.

Gene expression profiles (read counts per sample and gene) were subjected to Pearson pairwise correlation analysis and Principal Component analysis (PCA). Pearson pairwise correlations were converted into a distance matrix (distance = (1-correlation)/2) that was subjected to hierarchical clustering (agglomeration method: average). PCA was done after removal of all genes that were not detected in at least 80% of all samples with at least one read count, converting all remaining zero read counts into 0.1, $\log_{10}$ transformation and centering of the read counts matrix.

The raw scRNA-seq data were processed using the Cell Ranger Single-Cell Software Suite (version 4.0, 10x Genomics) with default parameters to generate an expression data matrix from all annotated genes and barcoded cells. QC for each sample included evaluation of the percentage of valid barcodes (must be >95%), mapping rates to the transcriptome (must be >60%) and median unique molecular identifier (UMI) counts per cell (must be >1000). Only data that passed these initial quality thresholds, and barcoded cells with at least 2,500 detected genes and less than 50% mitochondrial UMI counts were considered for further analysis using Seurat version 4.0.1. Based on the top 2000 features we used the Seurat algorithm SCTransform to regress out UMI count variation and percentage of mitochondrial genes in each of the 4 subject specific datasets. Datasets were merged using the Seurat IntegrateData functionality. After dimensionality reduction using only the top 20 principal components, we used the Seurat functionality 'FindNeighbors', followed by unsupervised clustering with a resolution of 0.1045, to identify five different clusters. Results were visualized after UMAP dimensionality reduction. We used 'AverageExpression' to identify average gene expression in each cluster and 'FindMarkers' to calculate differentially expressed genes of each cluster compared to all other clusters (adj. p-value ≤0.05). In both cases, we used the 'RNA' assay and 'counts' slot. Top 500 most abundant genes (as documented by average expression values) of each cluster were subjected to enrichment analysis using Fisher's Exact Test and the 'Human Gene Atlas' library obtained from the enrichR website (Chen et al., 2013; Kuleshov et al., 2016). Cluster-specific marker gene expression (i.e., differentially expressed genes) of published marker genes for iPSC-derived cardiomyocytes, epicardial cells, smooth muscle cells and fibroblasts obtained from D'Antonio-Chronowska et al (D'Antonio-Chronowska et al., 2019) was documented.

**RNA-seq-based PluriTest analysis**

For pluripotency analysis, the RNA sequence file of each sample was analyzed using PluriTest, a bioinformatic assay for pluripotency, by uploading the files via the web interface at https://pluritest.org/. For the FASTQ files that exceeded the platform's file size limit of 1.5GB, reads contained in the FASTQ files were randomly removed in order for the files to be trimmed down to the size limit before uploading. PluriTest returns the calculated pluripotency and novelty scores. For comparison, the pluripotency and novelty scores of human dermal fibroblasts from 66 individuals (available at https://www.ebi.ac.uk/arrayexpress/experiments/E-MTAB-7032/) (Hagai et al., 2018) were plotted alongside each duplicate sample per iPSC clone.

**RNA-seq-based comparative transcriptome analysis**

Alignment files (BAM-files) of 77 randomly selected hiPSC samples (Kilpinen et al., 2017) were obtained from the European Nucleotide Archive (analysis accession IDs: ERZ123025, ERZ123026, ERZ123027, ERZ123028, ERZ123029, ERZ123030, ERZ123031, ERZ123032, ERZ123033, ERZ123034, ERZ123035, ERZ123036, ERZ123037, ERZ123038, ERZ123039, ERZ123040, ERZ123041, ERZ123042, ERZ123043, ERZ123044, ERZ123045, ERZ123046, ERZ123047, ERZ123048, ERZ123049, ERZ123050, ERZ123051, ERZ123052, ERZ123053, ERZ123054, ERZ123055, ERZ123056, ERZ123057, ERZ123058, ERZ123059, ERZ123060, ERZ123061, ERZ123062, ERZ123063, ERZ123064, ERZ123065, ERZ123066, ERZ123067, ERZ123068, ERZ123069, ERZ123070, ERZ123071, ERZ123073, ERZ123074, ERZ123076, ERZ123077, ERZ123078, ERZ123079, ERZ123080, ERZ123081, ERZ123082, ERZ123083, ERZ123084, ERZ123085, ERZ123086, ERZ123087, ERZ123088, ERZ123089, ERZ123090, ERZ123091, ERZ266941, ERZ266942, ERZ266943, ERZ266944, ERZ266946, ERZ266947, ERZ266948, ERZ266949, ERZ266950, ERZ266951, ERZ266952, ERZ266953, available at https://www.ebi.ac.uk/ena/browser/view/PRJEB7388) and subjected to featureCounts for identification of read counts per gene and sample. FASTQ-files of 66 fibroblast samples (Hagai et al., 2018), which are previously used to generated hiPSC lines by Kilpinen and colleagues were downloaded from ArrayExpress (available at https://www.ebi.ac.uk/arrayexpress/experiments/E-MTAB-7032/), aligned to the human reference genome hg19 using STAR, followed by identification of read counts per gene and sample, using featureCounts. Gene expression profiles (read counts per sample and gene) of the three different datasets were merged and subjected to pairwise correlation and PCA as described above.

**Sendai Virus genomic RNA sequence mapping to RNA-seq data**

The sequence of the SeV primer used for mapping was created by extracting the first 181-bp sequence from the 5'-end RNA genome of Sendai virus (GenBank ID: X03614.1). This sequence is amplified by the SeV-specific primers (fwd: 5'-GGATCACTAGGTGATATCGAGC-3'; rev: 5'-ACCAGACAAGAGTTTAAGAGATATGTATC-3') described in the Thermo Fisher Scientific's CytoTune®-iPS 2.0 Sendai Reprogramming Kit manual for detection of the presence/absence of SeV by RT-PCR. An index file of the SeV sequence was generated by the BWA alignment software (version 0.7.17-r1188) using the command `bwa index sev-primer.fasta`. Using the generated index data of the SeV primer, each raw RNA-seq data file was aligned to the SeV primer by the command `bwa mem -t 8 sev-primer.fasta sample-seq.fastq > sample-seq.sam`. After alignment, data files in SAM format were generated for all 80 iPSC samples by the BWA aligner. The unique mapping flag values representing unmapped reads, forward-mapped reads, and reverse-mapped reads, were extracted with the corresponding counts of sequence reads, and tabulated for each iPSC sample. Alternatively, we performed RT-PCR on cDNA from iPSC lines/clones using the SeV-specific primers mentioned above.

**WGS and Mendelian disease/disorder analysis**

Genomic DNA was isolated using the PureLink$^{TM}$ Genomic DNA Mini Kit (Thermo Fisher Scientific, K1820-01). Library preparation and whole genome sequencing with 30x coverage was performed by The New York Genome Center or BGI Americas Corp (Cambridge, MA). Reads for each sample were aligned independently to the human reference genome GRCh37/hg19. Mapped reads representing the Sema4 Expanded Carrier Screen (ECS; 283 genes) panel were extracted from the bam files and FASTQ files generated. These files were analyzed through the Sema4 expanded carrier screening pipeline to generate vcf files. The vcf files were then analyzed using VaRank software (http://www.lbgi.fr/VaRank/)(Geoffroy et al., 2015) to generate a pathogenic disease prediction report and annotated using the open source SnpEff software (https://pcingola.github.io/SnpEff/). Furthermore, in order to characterize direct genetic risks in our samples, we annotated the called variants using Annovar software release 2019-10-2 with clinvar version 20200316 for the hg19 reference genome. Annotated vcfs were processed to extract only variants with passed previous quality filtering steps, GQ scores greater than 20, and were listed in clinvar (Landrum et al., 2014) as Pathogenic or Likely Pathogenic. Variants with conflicting interpretations and no assertion criteria in the database were excluded. Genes were filtered to include only genes associated with dominant genetic disorders. For genes, associated with both dominant and recessive disorders, the specific variant was reviewed by an ABMGG certified clinical geneticist (S.B.) and variant was included only if the genetic change was associated with a dominant presentation.

**Ancestry analysis**

The genetic admixture proportions for each sample were obtained by Sema4 using its ancestry pipeline, which integrates the ancestry inference algorithm from Gencove, Inc (New York, NY) with other widely used open source software, BWA1 (Li and Durbin, 2009) and SAMtools2 (Li et al., 2009), and an in-house highly curated reference set of 3.3M+ SNPs in the Gencove-assembled worldwide ancestry database consisting of 7,345 individuals grouped together into 49 populations. Each of these individuals is annotated with information about the ethnic origin/geographic location of the individual. Some of these data were sourced from public resources, specifically Phase 3 of the 1,000 Genomes Project and Lazaridis and colleagues (Lazaridis et al., 2016). Global ethnicities covered in the Gencove data base can be found at: https://ancestry.gencove.com/. Each ancestry analysis used a random subset of 150K SNPs and a total of 10 bootstraps were performed. A single bootstrap generated a '.Q' file which contained the ancestry fractions inferred for the sample. An average of the ancestry proportion values from each of these 10 bootstraps was used as the final result.

**Calcium transient recording and analysis**

hiPSC-cardiomyocytes were harvested in single-cell conditions and seeded at 25,000 cells per Matrigel-coated plastic-round coverslip (25mm, Sarstedt) 3-5 days before the calcium transient recordings. On the day of the experiment, cells were loaded with Tyrode's solution (140mM of NaCl, 5.4mM of KCl, 10mM of HEPES, 1mM of $NaH_2PO_4$, 1mM of $MgCl_2$, 10mM of glucose and 1.8mM of $CaCl_2$, pH 7.4) supplemented with 10 μmol/L Fluo-3 AM (Biotium, 5014) for 30 min at 37°C. After incubation, the cardiomyocytes were washed and superfused with fresh Tyrode's solution at 37°C. Fluo-3-AM-loaded cells were excited at 488 nm, and fluorescence above 505 nm was recorded by a confocal microscope (LSM 5 Exciter Carl Zeiss AG, Jena, Germany) at oil-40x magnification, using the line-scan mode. Calcium transients were recorded from 5 to 10 cells per coverslip under spontaneous beating behavior (20 to 30 seconds per cell). The dynamic images were stored as LSM files for further analysis.

Calcium transient recordings were assessed by an expert blinded to the experimental groups (R.D.). Data were processed and analyzed in MATLAB®2018a (MathWorks, Natick, MA) using customized scripts and functions. In brief, fluorescence signals were corrected by their background and expressed as F/F0 (the ratio of fluorescence during the calcium transient divided by fluorescence during diastole). The raw traces were filtered to reduce noise (*sgolayfilt* function in MATLAB). After filtering, representative peaks of each cell were selected. Different features regarding the transient duration and height were measured automatically using our functions. For the detailed comparison of recordings from 2 cell lines presented in Figure 5, we computed the following metrics from time courses: Ca peak, Ca baseline, Amplitude, Time-to-peak, AUC, CAD10, CAD15, CAD20, CAD30, CAD50, CAD70, CAD80, CAD90, Ca decay time, spontaneous beating rate. We expect that many of these features will be correlated, and the principal components analysis shown in Figure 5 is a way of adjusting for these correlations. For the less detailed comparison across 5 cell lines shown in Supplementary Figure S4A, the following features were calculated from each sample: Ca amplitude; CaD50; CaD90; time-to-peak; decay time; total calcium transient duration; spontaneous beating rate.

**Quantification and statistical analysis**

For the statistical analyses presented in Figure 5, the calcium transient recordings of a total of 164 cells were used. These consisted of 17, 12 and 16 atrial and 13, 13, and 13 ventricular cardiomyocytes for MSN14-01S, MSN14-05S and MSN14-06S, respectively, and 11, 14 and 3 atrial and 20, 20, and 12 ventricular cardiomyocytes for MSN25-01S, MSN25-05S and MSN25-09S, respectively.

For the analysis presented in Supplementary Figure S4A, the sample numbers were as follows: MSN01-03R, 14 cells; MSN02-04R, 48 cells; MSN06-07R, 40 cells; MSN08-13R, 17 cells; and MSN09-04R, 43 cells.

Principal Components Analysis (PCA) was performed on the calcium transient recordings and graphically plotted using MATLAB®2018a tools (Figures 6B and S4A).

The Kolmogorov-Smirnov test was used to analyze the statistical significance of the calcium transient duration and beat frequency between atrial and ventricular cardiomyocytes differentiated from hiPSC clones from the same individual and from two racially/ethnically diverse, age-matched males (Figure 6C-E).

**Data and software availability**

All hiPSC RNA-seq data files have been deposited at the Gene Expression Omnibus (GEO) under accession number GSE156384. The bulk and single-cell RNA-seq data files of cardiomyocytes differentiated form hiPSC lines have been deposited at GEO under accession numbers GSE174773 and GSE175761, respectively. The whole genome sequencing (WGS) data, enhanced carrier screen gene variant vcf files, subjects' clinical exam parameters and STR data have been deposited at the database of Genotypes and Phenotypes (dbGAP) under accession number phs002088.v2.p1. These data are restricted to requestors from not-for-profit organizations and is available through control-access requiring provision of the requester's local Institutional Review Board (IRB) approval documentation as required by the GDS Extramural Institutional Certification agreement between Mount Sinai and NIH (NHGRI). Requester must agree to make any results of studies using this data available to the larger scientific community.

**Additional resources**

The Mount Sinai Institute for Systems Biomedicine's page of our Library of Integrated Network-Based Cellular Signatures (LINCS) project, which was part of a consortium program of centers that was supported by the NIH Common Fund, contains additional resources and information such as data, metadata, expression signatures and standard operating procedures. The website can be accessed through https://icahn.mssm.edu/research/systems-biomedicine/resources/resources/lincs_project. Furthermore, all data and associated metadata, which are linked, can also be accessed through the searchable LINCS data portal at http://lincsportal.ccs.miami.edu/dcic-portal/. Finally, all hiPSC lines and clones have been registered with the Human Pluripotent Stem Cell Registry (http://hpscreg.eu).

### Icahn School of Medicine at Mount Sinai LINCS Center for Drug Toxicity Signatures

#### Standard Operating Procedure:
#### Skin Biopsy Punch Explant Culture for Derivation of Primary Human Fibroblasts

DToxS SOP Index: CE-6.0
Last Revision: 01/08/2016
Written By: Christoph Schaniel

Approvals (Date): Joseph Goldfarb (12/14/15)
Marc Birtwistle(12/14/15)
Eric Sobie (12/13/2015)
Ravi Iyengar (01/08/2016)

Quality Assurance/Control (QA/QC) steps are indicated with green highlight.

Metadata recording is highlighted with yellow highlight and [superscript indices].

---

**Note: All materials/reagents are listed in a table in the Metadata[1] section of this SOP.**

1. Assemble media

   a. 0.1% Gelatin: Weigh 1 g of gelatin[1] and dissolve in 1L DPBS[1]. Autoclave and filter under sterile conditions[1]. Store at +4°C. Keep for up to 4 weeks.
   b. Complete DMEM/20% FBS (500 mL): Mix under sterile conditions 380 mL of DMEM[1], 5 mL Penicillin/Streptomycin (100x)[1], 5 mL Non-Essential Amino Acids (100x)[1], 5 mL L-Glutamine (200 mM)[1], 5 mL SodiumPyruvate (100 mM)[1], 3.5 µL 2-mercoptoethanol (14.3 M)[1], 100 mL FBS[1]. Store at +4°C. Keep for up to 4 weeks.
   c. Freezing medium (100 mL): Mix under sterile conditions 40 mL DMEM[1], 50 mL FBS[1], 10 mL DMSO[1]. Store at +4°C. Keep for up to 1 week.

**Critical! Each individual biopsy/subject is to be kept in a separate plate/dish and all steps described below are to be handled separately for each sample to avoid any sample mix-up!**

2. Perform Skin Biopsy Punch

   a. Have subject lie on exam table with arm for biopsy above head.
   b. Sterilize inner upper arm with ChloraPrep[1] 2 times.
   c. Lay drape over subject's arm with sterilized area for punch biopsy exposed.
   d. Inject 1-2 cc of lidocaine[1] to anesthetize area. Wait until a slight poke with a needle is not felt (usually 2-3 min).
   e. Take skin sample using punch biopsy[1].
   f. Place biopsy in a sterile 15 mL conical tube containing 5 mL, ice-cold, complete DMEM/20% FBS (see *1b.*) and prelabeled with an individual number.
   g. Place tube back on ice.

h. Apply gauze to subject's arm and wrap with 3M Coban[1] self-adherent wrap.
i. Instruct subject to remove dressing after 1-2 hours and replace with a small adhesive bandage. Subject can shower the next day.

Note: The physician taking the skin biopsy punch will communicate the subject name and the corresponding tube number to Christoph Schaniel, who will associate the number with the de-identification code[2] to ensure that no participant name remains directly associated with his/her sample during the course of the study. Dr. Schaniel will store the participant name and corresponding sample de-identification code in a password protected file on password protected computer in a locked office as per HIPAA regulation. He and Dr. Iyengar are the only persons who will know the identity of each sample and can link it to its corresponding participant in accordance with HIPPA guidelines and as approved by Mount Sinai's Institutional Review Board.

**All subsequent steps are to be performed aseptically inside a biosafety cabinet!**

3. Prepare the Skin Punch Biopsy

a. Prepare in advance using a separate 6-well plate[1] per biopsy/subject (This is critical!): Label each plate with the de-identified sample name, passage number 0, and date[2,3].
b. Add 1 mL of 0.1% Gelatin (see *1a.*) to 3 wells of a 6-well plate.
c. Set the plate aside for 30-60 min.
d. Aspirate gelatin solution and add 750 µL of complete DMEM/20% FBS (see *1b.*) media to each well. Ensure that the entire surface of the well is covered with media.
e. Invert the lid of a sterile 100-mm petri dish[1] and add 1.5 mL of DMEM/20% FBS (see *1b.*) to the middle of the lid with a 5 mL serological pipette[1].
f. Spread out the media "drop" with the tip of the serological pipette. The "drop" will not be able to cover the entire lid surface.
g. Using sterile forceps[1], place the skin biopsy piece in the media on the dish lid.

4. Dissect the Skin PunchBiopsy

a. On the dish lid, dissect the 3-mm round skin biopsy into about 9 evenly sized pieces with smooth, even edges by cutting pieces using one sterile size 10 scalpel[1] to hold the biopsy in place and the second sterile scalpel to cut with a rolling motion in one direction. (Note: This step can be performed under a dissecting microscope (remember to maintain a sterile environment.) in order to better visual the biopsy and ensure cutting of even sized pieces with smooth even edges.) We routinely do it without the help of a microscope.

5. Transfer the Dissected Skin Biopsy Pieces into Tissue Culture Plates

a. Using pointed forceps[1] place 3 biopsy pieces per well into the prepared 6-well plate.
b. Use a tapping or sliding motion to get the pieces to attach to the bottom of the well.
c. Use a scalpel as above to remove any biopsy pieces that remain stuck to the forceps.
d. Place a sterile 22 mm cover slip[1] on top of the biopsy pieces with light pressure to ensure proper cell attachment to the bottom of the well.
e. Place the 6-well plate in a humidified incubator at 37°C/5% $CO_2$.
f. Monitor wells daily to ensure that the pieces are attached and that there is enough media covering the well during the first week; add ~200 µL of complete DMEM/20% FBS (see *1b.*) every 2 days to replace any evaporated media.
g. After one week, increase amount of media to 2 mL of complete DMEM/20%FBS (see *1b.*) and change media every 2-3 days.
h. Passage cells when fibroblasts have grown out and the well is confluent to the point where the fibroblasts are reaching the edges of the well (about 4-8 weeks).

6. Passaging of Fibroblasts

a. Prepare in advance using separate 6-well plate(s) per sample (This is critical!)
    i. Label each plate with the sample name (de-identification code), incremented passage number, and date.
    ii. Add 1 mL of 0.1% Gelatin (see *1a.*) per well (splitting ratio is 1:4; for 1st passage prepare 2 full 6-well plates).
    iii. Set the plate aside for at least 30 min – 60 min.
    iv. Aspirate gelatin solution and add 500 µL of complete DMEM/20% FBS (see *1b.*) to each well.
b. Take a representative phase-contrast picture at 4x magnification per sample ([4] and QA/QC1).
c. Trypsinize fibroblasts
    i. Aspirate medium from well(s) containing fibroblasts at ~80% confluence (QA/QC1) and wash once with DPBS (~1 mL/well). If any well is at 70% confluence or lower continue incubation of that well until 80% confluence is reached and process separately at that time.
    ii. Remove cover slip with a sterile forceps and place coverslip into a fresh well of a 6-well plate, with the side having faced the tissue up.
    iii. Inspect the coverslip using a microscope to see whether fibroblasts have attached to the coverslip. This occurs in most cases, and these cells will also be harvested (see next step).
    iv. Add 500 µL TrypLE[1] per well (including the wells carrying the coverslips with attached fibroblasts) and incubate for 3-5 minutes. Use a microscope to check when fibroblasts are rounding up or starting to lift off the bottom of the well. Proceed immediately to step *6d*.
d. Add 500 µL complete DMEM/20% FBS (see *1b.*) to each well.
e. Harvest fibroblasts using a 1 mL micropipette and pool cells from all wells into a 15 mL conical tube pre-filled with 4 mL complete DMEM/20% FBS (see *1b.*).
f. Centrifuge tube at 300 g for 5 minutes.
g. Aspirate medium, add 6 mL of complete DMEM/20% FBS (see *1b.*) and resuspend fibroblasts by repeated pipetting.
h. Add 500 µL of fibroblasts to each well, populating 12 wells (2 six well plates) in total.
i. Place the plate(s) in a humidified incubator at 37°C/5% $CO_2$.
j. Next day add another 1 mL of complete DMEM/20% FBS (see *1b.*) to each well.
k. Replace medium with 2 mL fresh complete DMEM/20% FBS (see *1b.*) every 2-3 days until wells are ~80% confluent with fibroblasts
l. Repeat Step 6 for 3 ~80%-confluent wells at each passage until 3rd passage is reached. Proceed with step 7 for all other wells when ~80% confluence is reached, and for all 3rd passage wells.

7. Cryopreservation/Freezing of Fibroblasts

a. Prepare in advance:
    i. Fresh freezing medium (see *1c.*). Keep cooled at 4˚C.
    ii. Cryovials: label with "Fib" (for fibroblast) followed by sample name/de-identification code, passage number, and date of freezing[3].
    iii. Precool vials in an isopropanol freezing vessel[1] at -80°C for 10-15 minutes.
b. When fibroblasts reach ~80% confluence (see pictures under QA/QC1), aspirate medium from each well and wash once with DPBS[1]. (~1 mL/well).
c. Add 500 µL TrypLE[1] per well of a 6-well plate and incubate for 3-5 minutes. Use a microscope to check when fibroblasts are rounding up or starting to lift off the bottom of the well. Proceed immediately to step *7d*.
d. Add 500 µL complete DMEM/20% FBS (see *1b.*).

e. Harvest fibroblasts using a 1 mL micropipette and combine cells from all wells into a conical tube (15 mL for up to 6 wells or 50 mL for more than 6 wells) filled with 4 mL of complete DMEM/20% FBS (see *1b.*).
f. Use 10 µL to determine the total number of fibroblasts per mL with a hemocytometer.
g. Centrifuge tube at 300 g for 5 minutes.
h. Aspirate medium and add freezing medium (see *1c.*) so that fibroblasts are at ~$5x10^5$ cells/mL.
i. Add 1 mL of the cell solution per cryovial.
j. Freeze using a Cryo1°C freezing container[1] (filled with 250 mL 2-Propanol) at -80°C overnight.
k. Move vials to a liquid nitrogen tank for long-term storage the next day.[3]

#### Metadata

1. **Materials/Reagents:** Company name, Catalogue and lot numbers

| PRODUCT | COMPANY NAME | CAT # | Storage | Usable life |
|---|---|---|---|---|
| DMEM (500 mL) | Life Technologies | 11965-118 | +4°C | M |
| Penicillin/Streptomycin (100x) | Life Technologies | 15140-122 | +4°C | M |
| Sodium-Pyruvate (100 mM) | Life Technologies | 11360-070 | +4°C | M |
| L-Glutamine (200 mM) | Life Technologies | 25030-081 | +4°C | M |
| Non-Essential Amino Acids (100x) | Life Technologies | 11140-050 | +4°C | M |
| Fetal Bovine Serum (FBS; 500 mL) | Corning | 35-011-CV | +4°C | M |
| 2-Mercaptoethanol | MP Biomedicals | 194705 | RT | N/A |
| TrypLE Express (1x) | Life Technologies | 12605010 | +4°C | M |
| Gelatin | Sigma | G1890 | RT | M |
| Dulbecco's phosphate-buffered saline (DPBS) | Life Technologies | 14190-136 | +4°C | M |
| DMSO | Fisher Scientific | BP2311 | RT | M |
| Biopsy Punches | Integra Miltex | 33-32 or 33-22 | RT | N/A |
| Disposable scalpel size 10 (Sterile) | Exel International | 29550 | RT | N/A |
| Pointed forceps (sterile) | Fisher Scientific | 08-0887 | RT | N/A |
| ChloraPrep (3mL) | Carefusion | 260400 or [illegible] | RT | M |
| Lidocaine (1%) | HospiraInc | 0409-3178-03 | RT | M |
| 3M Coban self-adherent wrap | 3M | 1581 | RT | N/A |
| Gauze Sponges (4" x 4") | Medline | 60-21408C | RT | N/A |
| 1 L Sterile Disposable Filter | Fisher Scientific | 09-741-03 | RT | N/A |
| 5 mL tubes | Fisher Scientific | 14-956-3C | RT | N/A |
| 15 mL conical tube | BD FALCON | 352099 | RT | N/A |
| 100-mm petri dish (sterile) | BD FALCON | 351029 | RT | N/A |
| 6 well Tissue Culture (TC) plate | Corning | 353046 | RT | N/A |
| 2 mL Aspirating pipettes | BD FALCON | 357558 | RT | N/A |
| 5 mL Serological pipettes | BD FALCON | 357543 | RT | N/A |
| 10 mL Serological pipettes | BD FALCON | 357551 | RT | N/A |
| 1000 µL Blue graduated pipet tip | USA Scientific | 1111-2721 | RT | N/A |
| 200 µL Natural pipet tip | USA Scientific | 1111-0700 | RT | N/A |
| 0.1-10 µL Graduated pipet tip | USA Scientific | 1111-3700 | RT | N/A |
| 22 mm coverslip | Fisher Scientific | NC9304304 | RT | N/A |
| Hemocytomter | Fisher Scientific | 0267110 | RT | N/A |
| Cryovials | Nunc | 377367 | RT | N/A |
| Cryo 1°C freezing container | Nalgene | 5100-0001 | RT | N/A |

Continued - Materials/Reagents

| PRODUCT | COMPANY NAME | CAT # | Storage | Usable life |
|---|---|---|---|---|
| 2-Propanol | Fisher Scientific | A417-4 | RT | N/A |
| EVOS XL Core Imaging System # | Fisher Scientific | 12-562-751 | RT | N/A |

M; according to manufacturer's shelf-life information; RT, room temperature; #, or equivalent microscope

**2. Subject(s):** subject ID (de-identified), age, gender/sex, race/ethnicity

**3. Fibroblast(s):** subject ID (de-identified), passage number, dates of each passage, date of biopsy, date of initial plating, date of freezing.

**4. Microscopy pictures:** subject ID (de-identified), passage number, date, microscope name (company, catalogue number), magnification.

#### Quality Assurance/Control Steps (QA/QC)

**1: Microscopy pictures at 4x magnification.**

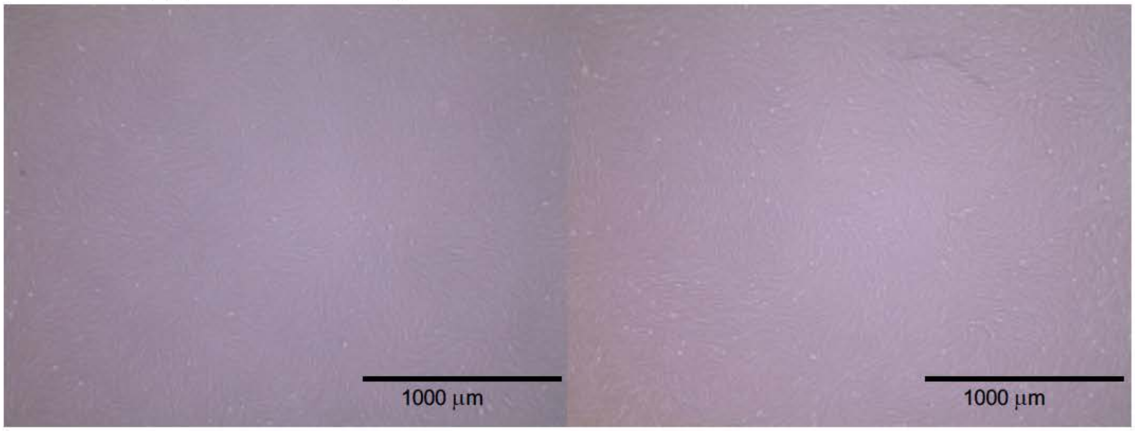

**Representative images of established fibroblast lines from two independent healthy subjects at passage 2 before splitting.**

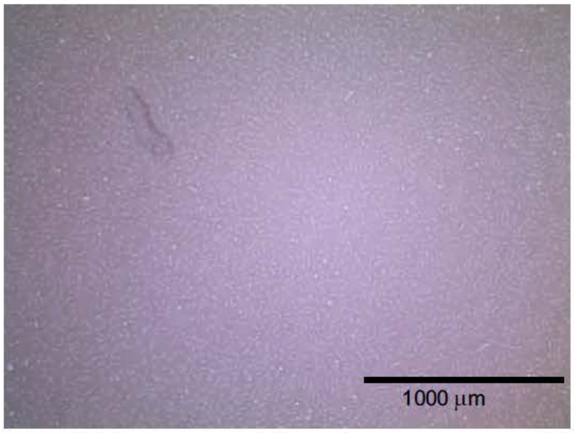

**The above image is representative of an established fibroblast line that is not quite yet ready for harvesting (would need another day or two).**

### Icahn School of Medicine at Mount Sinai LINCS Center for Drug Toxicity Signatures

#### Standard Operating Procedure: Reprogramming Human Fibroblasts to Human Induced Pluripotent Stem Cells (hiPSCs) using the mRNA with microRNA boost method

DToxS SOP Index: CE-7.0
Last Revision: 01/17/2018
Written By: Christoph Schaniel

Approvals (Date): Joseph Goldfarb ( 1/18/18)
Marc Birtwistle (1/17/18)
Eric Sobie ( 1/17/18)
Ravi Iyengar ( 1/17/18)

Quality Assurance/Control (QA/QC) steps are indicated with green highlight.

Metadata recording is highlighted with yellow highlight and superscript indices.

---

**Note: All materials/reagents are listed in a table in the Metadata[1] section of this SOP.**

1. Assemble media and reprogramming reagents

   a. 0.1% Gelatin: Weigh 1 g of gelatin[1] and dissolve in 1L DPBS[1]. Autoclave and sterile filter[1]. Store at +4°C. Keep for up to 4 weeks.
   b. Matrigel/DMEM: Thaw Matrigel on ice and at +4°C (cold room) overnight. Under sterile conditions and on ice, add equal volume of DMEM[1] to the thawed Matrigel solution[1]. Keep for up to 4 months at -20°C or up to 4 weeks at +4°C.
   c. Complete DMEM/20% FBS (500 mL): Mix under sterile conditions 380 mL of DMEM[1], 5 mL Penicillin/Streptomycin (100x)[1], 5 mL Non-Essential Amino Acids (100x)[1], 5 mL L-Glutamine (200 mM)[1], 5 mL Sodium Pyruvate (100 mM)[1], 3.5 µl 2-mercoptoethanol (14.3 M)[1], and 100 mL FBS[1]. Store at +4°C. Keep for up to 4 weeks.
   d. Complete DMEM/20% FBS without antibiotics (500 mL): Mix under sterile conditions 385 mL of DMEM[1], 5 mL Non-Essential Amino Acids (100x)[1], 5 mL L-Glutamine (200 mM)[1], 5 mL Sodium Pyruvate (100 mM)[1], 3.5 µl 2-mercoptoethanol (14.3 M)[1], 100 mL FBS[1]. Store at +4°C. Keep for up to 4 weeks.
   e. Complete DMEM/10% FBS (100 mL): Mix under sterile conditions 86 mL DMEM[1], 1 mL Penicillin/Streptomycin (100x)[1], 1 mL Non-Essential Amino Acids (100x)[1], 1 mL L-Glutamine (200 mM)[1], 1 mL Sodium-Pyruvate (100 mM)[1], 0.7 µL 2-mercoptoethanol[1] (14.3 M) in DMEM, 10 mL FBS[1]. Store at +4°C. Keep for up to 4 weeks.
   f. Complete mTeSR medium: Mix under sterile conditions 400 mL of mTeSR[1] with 100 ml mTeSR 1.5X Supplement[1] and 5 mL Penicillin/Streptomycin (100x)[1].
   g. Pluriton/Penicillin/Streptomycin (Pluriton/PS) medium: Under sterile conditions,
      i. Thaw frozen Pluriton Medium[1] at +4°C over 2 days.
      ii. To 495 mL Pluriton Medium add 5 mL of Penicillin/Streptomycin (100x)[1].
      iii. Transfer 250 mL into a separate sterile container and store at 4°C. This is used to generate MEF conditioned Pluriton/PS Medium (see *Step 2*).

iv. Store the remaining 250 mL Pluriton/PS medium at -20°C.

h. Pluripotent stem cell (PSC) medium (500 mL): Under sterile conditions
   i. Supplement DMEM/F12 (500 mL)[1] with 5 mL Penicillin/Streptomycin (100x)[1] and 5 mL L-Glutamine (200 mM)[1].
   ii. To 400 mL Supplemented DMEM/F12, add 100 mL KnockOut Serum Replacement,[1] 5 mL Non-Essential Amino Acids (100x)[1], 350 µL of 2-mercaptoethanol[1] that has been diluted 1:100 in DMEM/F12, and sterile filter.
   iii. Add 1 mL FGF2/BSA[1] [10 µg/mL] (final concentration: 20 ng/mL).
   iv. Wrap bottle in aluminum foil to protect from light at all times. Store at +4°C. Keep for up to 4 weeks.

i. Thiazovivin [10 mM]: Under sterile conditions
   i. Suspend 10 mg Thiazovivin powder in 3.212 mL of DMSO to obtain a 10 mM stock.
   ii. Place 50 µL aliquots into individual sterile, low protein binding 0.5 mL microcentrifuge tubes[1].
   iii. Freeze and store at -20°C until use. Keep for up to 6 months.

j. Pluriton Supplement (0.2ml): Under sterile conditions,
   i. Thaw the Pluriton Supplement[1] on ice. Once thawed, keep on ice at all times.
   ii. Centrifuge (4°C or room temperature) the vials to collect the contents at the bottom of the well at 13000 rpm for 10 sec.
   iii. Using RNAse-free filter tips[1] mix by pipetting up and down.
   iv. Place 1.7 µL aliquots of Pluriton Supplement into individual sterile, low protein binding 0.5 mL microcentrifuge tubes[1] (each is enough to supplement 4 mL Pluriton/PS medium at 0.4 µL/mL).
   v. Store at -80°C until use.

k B18R: Under sterile conditions,
   i. Thaw B18R[1] vials on ice. Once thawed, keep on ice at all times.
   ii. Centrifuge (4°C or room temperature) the vials to collect the contents at the bottom of the well at 13000 rpm for 10 sec.
   iii. Using RNAse-free filter tips[1], place 4.1 µL aliquots of B18R protein [500 µg/mL] into individual sterile, RNAse-free 0.5 mL tubes[1] (each is enough to supplement 4 mL Pluriton/PS medium at 1 µL/mL or 500 ng/mL).
   iv. Store at -80°C until use.

l. FGF2/BSA: Reconstitute FGF2[1] at 10 µg/mL in sterile PBS containing 0.1% (0.1 g / 100mL PBS) bovine serum albumin. Store at -20°C for up to 6 months. Store at +4°C and keep up to 4 weeks.

m. mRNA Reprogramming cocktail: Under sterile conditions,
   i. Thaw the OCT4, SOX2, KLF4, c-Myc, LIN28 and nucGFP mRNA vials[1] contained in the mRNA reprogramming kit on ice. Once thawed, keep on ice at all times.
   ii. Centrifuge (4°C or room temperature) the vials to collect the contents at the bottom of the well at 13,000 rpm for 10 sec.
   iii. Using RNAse-free filter tips[1] and a 200 uL micropipette, measure the volume of each mRNA solution.
   iv. Using RNAse-free filter tips[1], combine the mRNA factors at the ratio of volumes listed below in a sterile, RNAse-free 1.5 mL microcentrifuge tube[1] on ice. Scale up to maximize volume (OCT4 is usually the limiting component/volume).

| | |
|---|---|
| OCT4 mRNA | 400.0 µL |
| SOX2 mRNA | 123.8 µL |
| KLF4 mRNA | 162.0 µL |
| c-Myc mRNA | 153.5 µL |
| Lin28 mRNA | 85.7 µL |
| nGFP mRNA | 115.0 µL |

   v. Mix the content of the tube by pipetting up and down.
   vi. Place 40 µL aliquots of this mRNA reprogramming cocktail into individual sterile, RNAse-free 0.5 mL tubes[1] (each tube is sufficient for transfecting 4 wells).

vii. Store at -80°C until use

n. microRNA booster Kit:

    i. Thaw microRNA vials[1] on ice. Once thawed, keep on ice at all times.
    ii. Centrifuge (4°C or room temperature) vials to collect the content at the bottom of the well at 13,000 rpm for 10 sec.
    iii. Using RNAse-free filter tips[1] mix by pipetting up and down.
    iv. Aliquot 14 µL of this microRNA cocktail into individual sterile, RNAse-free 0.5 mL tubes[1] (each tube is sufficient for transfecting 4 wells).
    v. Freeze and store at -80°C until use.

o. hiPSC freezing medium (50 mL): 25 mL PSC medium, 20 mL FBS[1], 5 mL DMSO[1]. Store at +4°C. Keep for up to 1 week.

2. Generation of MEF-conditioned Pluriton medium

a. Remove 2 vials of irradiated MEFs[1] from liquid nitrogen storage and keep on dry ice.
b. Under sterile conditions, add 250 µL of pre-warmed DMEM/10% FBS (*Step 1.d.*) to each cryovial and then thaw each vial at 37°C for 1-3 minutes until only a tiny frozen cube remains.
c. Transfer the contents of the two vials to a single 50 mL conical tube containing 5 mL of pre-warmed complete DMEM/20% FBS (*Step 1.c.*)
d. Centrifuge the cells for 5min at 300 g at 4-8°C.
e. Aspirate the supernatant and resuspend each cell pellet in 40 mL of complete DMEM/20% FBS
f. Add 20 mL of each cell suspension to an individual 75cm$^2$ tissue culture flask[1].
g. Evenly distribute the MEF cell suspensions in the flasks.
h. Incubate the cells overnight at 37°C and 5% $CO_2$.
i. Replace medium with 20 mL of pre-warmed Pluriton/PS medium (*Step 1.g.*) and 16 µL FGF2/BSA (*Step 1.l.*) per flask.
j. Incubate the cells overnight at 37°C and 5% $CO_2$.
k. Collect medium, filter through a 50 ml rapid-flow conical tube filter with a 0.2 µm aPES membrane[1] and use this MEF-conditioned Pluriton medium in Step 6. Freeze leftover MEF-conditioned Pluriton/PS medium in a 50 mL conical tube at -20°C. Can be kept at -20°C up to 6 months.
l. Repeat *Steps 2.i.* through *Step 2.k.* daily until reprogramming is done (see *Step 6*).

3. Thawing of cryopreserved fibroblasts

**Critical! Cells from each individual subject (fibroblasts or induced pluripotent stem cells) are to be kept in a separate plate/dish and all steps described below are to be handled separately for each sample to avoid any sample mix-up and/or cross contamination.**

a. Prepare in advance using a separate 6-well plate per fibroblast line/subject (This is critical to minimize the risk of sample mix-up and cross-contamination!): Label each plate with the de-identified sample name, fibroblast passage number (passage number at time of freezing +1), and date (metadata also includes the date of freezing). Add 1 mL of 0.1% Gelatin (*Step 1.a.*) to 2 wells of a 6-well plate. Set the plate aside for 30-60 min. Aspirate gelatin solution and add 1 mL of complete DMEM/20% FBS (*Step 1.c.*) to one gelatin well (well 1) and 1 mL of complete DMEM/20% FBS without antibiotics (*Step 1.d.*) to the other gelatin well (well 2). Ensure that the entire surface of the well is covered with media. The antibiotic-free well will be used for mycoplasma testing.
b. Remove a cryovial with fibroblast passage number < 3 from the liquid nitrogen storage tank and keep on dry ice.
c. Add 250 µL of complete DMEM[1]/20% FBS[1] without antibiotics (*Step 1.d.*) to the cryovial and thaw at 37°C for 1-3 minutes until only a tiny frozen cube remains.
d. Under sterile conditions, transfer cell suspension into a 15 mL conical tube filled with 4 mL

complete DMEM[1]/20% FBS[1] without antibiotics (*Step 1.d.*).

e. Centrifuge tube at 300 g for 5 min at 4°C.
f. Aspirate medium, add 2 mL of complete DMEM[1]/20% FBS[1] without antibiotics (*Step 1.d.*), resuspend cells gently and add 1 mL per well (2 wells in total).
g. Place the plate(s) in a humidified incubator at 37°C / 5% $CO_2$.
h. Next day, replace medium with 2 mL fresh complete DMEM/20% FBS *with* antibiotics (*Step 1.c.*) for well 1 and 2 mL fresh complete DMEM/20% FBS *without* antibiotics for well 2.
i. Once cells are ~80% confluent (usually 2-3 days after thawing) proceed to *Step 4* for the well with antibiotic-free medium (well 2) and Step *5* for the well with antibiotic-containing medium.

4. QA/QC1Mycoplasma testing of fibroblasts

a. Add 500 µL TrypLE Express (1x)[1] to well 2 and incubate for 3-5 minutes. Use a microscope to check when fibroblasts are rounding up or starting to lift off the bottom of the well.
b. Add 500 µL complete DMEM/20% FBS without antibiotics (*Step 1.d.*).
c. Harvest fibroblasts using a 1 mL micropipette tip and add cells into a 15 mL conical tube filled with 9 mL sterile DPBS[1].
d. Centrifuge tubes at 300 g for 5 minutes at 4°C.
e. Aspirate medium and resuspend fibroblasts in 200 µl DPBS[1].
f. Proceed with DNA isolation using and following the Pure LINK™ Genomic DNA Mini Kit protocol (see Appendix A).
g. Proceed with Mycoplasma testing using and following the e-Myco™ plus Mycoplasma PCR Detection KIT according to the manufacturer's instructions (see Appendix B).
h. Discard any positive cultures and start over.

5. Plating of fibroblasts for reprogramming

a. Prepare in advance using a separate 6-well plate per fibroblast line/subject (This is critical to minimize the risk of sample mix-up and cross-contamination!): Label each plate with the de-identified sample name, fibroblast passage number, and date. Keeping plate on ice at all times, add 1 mL of Matrigel/DMEM further diluted 1:30 (v/v) in DMEM[1] (final dilution ratio is 1:60) to 2 wells of a 6-well plate. Set the plate aside for 30-60 min on ice. Aspirate Matrigel/DMEM and add 0.5 mL of complete DMEM/20% FBS (Step 1.c.) to each well. Ensure that the entire surface of the well is covered with media.
b. Aspirate medium from the remaining well containing fibroblasts at 80% confluence (*Step 3.i.*) and wash once with DPBS[1] (~1 mL/well).
c. Add 500 µL TrypLE[1] to the well and incubate for 3-5 minutes. Use a microscope to check when fibroblasts are rounding up and starting to lift off the bottom of the well.
d. Add 500 µL complete DMEM/20% FBS (*Step 1.c.*).
e. Harvest fibroblasts using a 1 mL micropipette tip and transfer to a 15 mL conical tube filled with 4 mL of complete DMEM/20% FBS (*Step 1.c.*).
f. Use 10 µL from the tube with cells in complete DMEM/20% FBS to determine the total number of fibroblasts with a hemocytometer.
g. Centrifuge tube at 300 g for 5 minutes at 4°C.
h. Aspirate medium and resuspend fibroblasts in complete DMEM/20% FBS (*Step 1.c.*) to 100,000 cells/mL.
i. Important note: Primary fibroblast cultures can exhibit different proliferation rates. As a result, faster growing fibroblast cultures will receive a lower microRNA/mRNA cocktail dose per cell then slower growing cells. Thus, plating fibroblasts may require adjusting the cell number depending on their proliferation index. Commonly, we
   i. Add 0.5 mL of cell suspension (50,000 cells) to one pre-coated well of a 6-well plate, and
   ii. Add fewer or more fibroblasts to another pre-coated well of a 6-well plate depending on whether they proliferate faster or more slowly.

j. Thaw 1 tube of B18R protein (*Step 1.k.iv.*) on ice, add 4 µL of B18R protein to 2 mL of DMEM/20% FBS (*Step 1.c.*) medium, and add 0.5 mL of DMEM/20%FBS/B18R medium per well.
k. Place the plate(s) in a humidified incubator at 37°C / 5% $CO_2$ (D0).

6. Reprogramming of fibroblasts

**Critical** **Reprogram only those fibroblasts/wells that are mycoplasma-free (step *4*)!**
**Reprogramming cocktail needs to be added every 24 hrs (± 30 min)**

**Note: The below amounts are for reprogramming 4 wells with fibroblasts of a 6-well plate. If fewer or more wells with fibroblasts are reprogrammed to tubes, volumes need to be scaled accordingly. The microRNA transfection is done first, followed by multiple rounds of mRNA transfection.**

a. QA/QC2 On day 1 (D1), take pictures of mycoplasma-free fibroblasts.
b. Thaw 2 aliquots of Pluriton Supplement[1] (*Step 1.j.v.*) on ice and add 3.2 µL per 8 mL of MEF-conditioned Pluriton medium[1] (see *Step 2*) in a 50 mL conical tube[1].
c. Thaw 2 aliquots of B18R[1] protein (*Step 1.k.iv.*) on ice and add 8 µL per 8 mL of Pluriton-supplemented, MEF-conditioned Pluriton medium[1] to the tube (Step *6b*).
d. Aspirate medium from fibroblast reprogramming wells and add 2 mL of supplemented, MEF-conditioned Pluriton medium with B18R protein per well. The volume of media added depends on the confluency of the cells; >85% confluence add only 1mL of supplemented, MEF-conditioned Pluriton medium with B18R protein, 70-85% add 1.5 mL, <70% add 2mL.
e. Place the plate(s) in a humidified incubator at 37°C / 5% $CO_2$ for at least 2 hrs.
f. Remove Stemfect RNA Transfection reagent[1] and Stemfect Buffer[1] (part of Stemfect Trasnfection Kit[1]) from +4°C refrigerator and equilibrate to room temp for 15-30 min.
g. Thaw 1 aliquot of microRNA cocktail (*Step 1.n.v.*) per 4 wells of fibroblasts on ice.
h. Prepare two 1.5 mL sterile, RNAse-free microfuge tubes[1] and using RNAse-free filter tips[1], add:
    i. To Tube 1: 14 µL of microRNA cocktail, followed by 86 µL of Stemfect Buffer[1] (sufficient for transfection of 4 wells). Mix well by pipetting up and down.
    ii. To Tube 2: 16 µL of Stemfect RNA Transfection reagent[1] followed by 84 µL of Stemfect Buffer[1] (sufficient for transfection of 4 wells). Mix well by pipetting up and down.
i. Using an RNAse-free filter tip[1], transfer the contents of Tube 2 to Tube 1 and mix by gently pipetting 3-5 times up and down. This is the microRNA transfection complex.
j. Incubate the microRNA transfection complex at room temperature for 15 min.
k. Holding the plate with the fibroblasts at a 45° angle, add 50 µL of microRNA transfection complex dropwise to each well to be transfected using a Pipette with a 200 µl RNAse-free filter tip[1].
l. Gently rock plate from front to back and side to side, to evenly distribute the microRNA transfection mixture.
m. Place the plate(s) in a humidified incubator at 37°C / 5% $CO_2$ and incubate for 24 hrs.
n. Next day (D2), check cells under a microscope. If there is a lot of cell death (i.e. rounded, non-attached cells), increase medium from 1 mL to 1.5. mL or 2 mL see guidelines in Step *6.d*.
o. Thaw on ice and add 2 aliquots of Pluriton Supplement[1] (*Step 1.j.v.*) and 2 aliquots of B18R protein[1] (*Step 1.k.iv.*) per 8 mL of MEF-conditioned Pluriton medium (see *Step* 2).
p. Aspirate medium from well(s) and add 2 mL, of supplemented, MEF-conditioned Pluriton medium with Pluriton Supplement and B18R protein per well.
q. Place the plate(s) in a humidified incubator at 37°C / 5% $CO_2$.
r. Remove Stemfect RNA Transfection reagent[1] and Stemfect Buffer[1] from +4°C refrigerator and equilibrate to room temp for 15-30 min.
s. Thaw, on ice, 1 aliquot of mRNA reprogramming cocktail (*Step 1.m.vii.*) per 4 wells of fibroblasts.

t. Prepare two 1.5 mL sterile, RNAse-free microfuge tubes and using RNAse-free filter tips, add:
    i. To Tube 1: 40 µL of mRNA reprogramming cocktail, followed by 60 µL of Stemfect Buffer (sufficient for transfection of 4 wells). Mix well by pipetting up and down.
    ii. To Tube 2: 16 µL of Stemfect RNA Transfection Reagent followed by 84 µL of Stemfect Buffer (sufficient for transfection of 4 wells). Mix well by pipetting up and down.
u. Using an RNAse-free filter tip, transfer the content of Tube 2 to Tube 1 and mix by gently pipetting 3-5 times up and down to produce the mRNA transfection complex.
v. Incubate the mRNA transfection complex at room temperature for 15 min.
w. Holding the plate with the fibroblasts at an 45° angle, add 50 µL of mRNA transfection complex dropwise to each well to be transfected using a Pipette with a 200 µl tip[1]
x. Gently rock plate from front to back and side to side, to evenly distribute the mRNA transfection mixture.
y. Place the plate(s) in a humidified incubator at 37°C / 5% $CO_2$ and incubate for 24 hrs.
z. Repeat *Steps 6n-6y* on D3 and D4.
aa. D5, check cells under a microscope and adjust medium volume according to amount of cell death as described in *6n*.
bb. D5 - Repeat *Steps 6b-6k* (microRNA transfection) and *6s-6y* (mRNA transfection) together.
cc. D6-D12 repeat *Steps 6n-6y*. QA/QC3 If by D12 the morphology of the reprogrammed fibroblasts not yet similar enough to establish human pluripotent stem cell lines, continue the mRNA transfection for an additional 1-2 days.
dd. Continue feeding cells daily with supplemented, MEF-conditioned Pluriton medium (*Step 2*) for an additional 1-3 days or until colonies are large enough to be picked and expanded (see *Step 7*).

7. Picking and replating induced pluripotent stem cells (feeder-free picking, preferred method)

a. When hiPSC colonies reach >1,000 cells prepare one 24-well plate per sample/subject. Label plate with the de-identified sample name and date. (METADATA also should include the fibroblast passage number, date of freezing & thawing, date of reprogramming, date of picking). Add 500 µL of Matrigel/DMEM (*Step 1.b.*) at a final 1:60 (v/v) dilution in DMEM[1] per well of a 24-well plate. Set the plates aside for 30-60 min at RT
b. Aspirate the Matrigel/DMEM from wells and replace with 100 µL per well of complete mTeSR medium (*Step 1.f.*) supplemented with Thiazovivin[1] (*Step 1.i.iii.*) 1:5,000 (v/v).
c. Pick individual colonies using physical scraping and sucking into a 20 ul filter tip with the help of an inverted microscope (in the hood) and place each picked colony into an individual well. This is considered passage 1. Note, sample name is now MSN_patient de-idenification code-N, where N is the colony number (e.g., MSN01-1).
d. The next day, and on subsequent days, replace medium with 250 µL fresh complete mTESR medium (*Step 1.f.*) per well until colonies reach a respectable size (at least 500-1000 cells) and retain an undifferentiated morphology (QA/QC4). (Go to *Step 8*)

.

8. Splitting/expansion of induced pluripotent stem cells (MEF/feeder-free)

**Note: This step details passaging the cells from the 24 well plate onto a 12 well plate and finally onto a 6 well plate**

a. At least one hour before passaging from a 24 well plate, coat the wells of a 12 well plate with 500 µL Matrigel/DMEM diluted at a final ratio of 1:60 v/v Matrigel/DMEM. If passaging from a 12 well plate use 1 mL for each well of a 6-well plate. Note: Label plate with the de-identified sample name and date. (METADATA also should include the fibroblast passage number, date of freezing & thawing, date of reprogramming, date of picking).
b. Wash cells in a well of a 24 well plate with 250 µL/well of calcium- and magnesium-free

phosphate-buffered saline (DPBS)[1] or 500 µL/well in the case of a 12 well plate and aspirate.

c. Add 250 µL/well (24 well format), or 500 µL/well (12 well format) of ReLeSR™[1] and aspirate ReLeSR™[1] within one minute, so that colonies are exposed to a thin film of liquid.
d. Incubate the culture at 37°C for 6 minutes.
e. Prepare complete mTESR medium (*Step 1.f.*) supplemented with Thiazovivin (*Step 1.i.iii.*) at a 1:5000 dilution. Add 500 µL/well (24 well format) or 1 mL/well (12 well format) of complete mTESR medium + Thiazovivin (only for day 1 of splitting).
f. Detach the colonies by holding the plate with one hand and use the other hand to firmly tap the side of the plate for approximately 30 - 60 seconds.
g. Once the cell aggregates start lifting off, remove cells and re-plate the cell aggregate mixture from the well of a 24 well plate into 1 well of the 12 well plate pre-coated with Matrigel (*Step 8.a.*) or from the well of a 12 well plate into 1 well of the 6-well plate pre-coated with Matrigel (*Step 8.a.*)
h. Place the plate in a 37°C incubator. Move the plate in several quick, short, back-and-forth and side-to-side motions to evenly distribute the cell aggregates. Do not disturb the plate for 24 hours.
i. Perform daily medium changes using complete mTESR medium (no Thiazovivin; 1 ml/well of a 12 well and 2 ml/well of a 6 well plate) and visually assess cultures to monitor growth until the next passaging time.

9. Expansion (6 well plate format) of about 80% confluent wells and cryopreservation/freezing of induced pluripotent stem cells

a. Prepare in advance:
    i. Fresh hiPSC freezing medium (*Step 1.o.*). Keep cooled at 4°C.
    ii. Cryovials: label 6 vials with "MSN_patient de-idenification code-N, where N is the picking/clone number passage number (p), and date of freezing (e.g., MSN01-1, p10, 04052016). Precool vials in an isopropanol freezing vessel at -80°C for 10-15 minutes.
b. Split 1/6 of a well of a 6 well plate into a fresh well of a 6 well plate following *Steps 8a-8i* but washing cells with 1mL/well of calcium- and magnesium-free phosphate-buffered saline (DPBS)[1] using 1 mL/well of ReLeSR™[1] , and instead of transferring 1 mL only transfer 166 µL to one well of a Matrigel/DMEM coated 6 well plate. These cells can continue to be expanded for subsequent experiments. Track passage number.
c. The remaining cells in the well are cryopreserved at this point as a precaution to preserve the cell line in the event of unexpected problems during the expansion phase.
    i. Transfer the 834 µL of hiPSCs into a 15 mL conical tube.
    ii. Centrifuge at 300 g for 5 min at 4°C.
    iii. Aspirate medium and resuspend cells in 1 ml of of hiPSC freezing medium (*Step 1.o.*).
    iv. Transfer cells to the cryovial and freeze using a Cryo 1°C freezing container[1] (filled with 250 mL 2-Propanol[1]) at -80°C overnight then move vials to a liquid nitrogen tank for long-term storage.

**Concluding remarks**

We had multiple reprogramming successes using this method. However, we subsequently ran into reprogramming failures for various fibroblast lines on multiple attempts. The failures are unlikely attributable to the subject's age, gender or race/ethnicity at the time of biopsy, the specimen handling and processing, fibroblast culture or staff. In the failed reprogramming attempts we observed increased fibroblast death during the reprogramming process. We attempted to control this by increasing the fibroblasts numbers as well as exchanging the reprogramming medium with fresh 2 mL of supplemented, MEF-conditioned Pluriton medium with B18R protein 6 hrs or more after transfection. Unfortunately, this did not resolve our problem. We subsequently observed that during the generation of MEF-conditioned Pluriton medium many MEFs died. We concluded that the Pluriton medium was cytotoxic (reason unknown). Stemgent, has discontinued to sell Pluriton medium. In addition, the RNA reprogramming technology has evolved to new, next generation non-modified and self-replicative RNA and microRNA technologies.

**Metadata**

1. **Materials/Reagents:** Company name, Catalogue and lot numbers

| PRODUCT | COMPANY NAME | CAT # | Storage | Usable life |
|---|---|---|---|---|
| DMEM (500 mL) | Life Technologies | 11965-118 | +4°C | M |
| IMDM | Life Technologies | 12440-053 | +4°C | M |
| DMEM/F12 | Life Technologies | 11330057 | +4°C | M |
| Penicillin/Streptomycin (100x) | Life Technologies | 15140-122 | +4°C | M |
| Sodium-Pyruvate (100 mM) | Life Technologies | 11360-070 | +4°C | M |
| L-Glutamine (200 mM) | Life Technologies | 25030-081 | +4°C | M |
| Non-Essential Amino Acids (100x) | Life Technologies | 11140-050 | +4°C | M |
| Fetal Bovine Serum (FBS; 500 mL) | Corning | 35-011-CV | +4°C | M |
| EDTA | Corning | 46-034-CI | RT | M |
| KnockOut Serum Replacement | Life Technologies | 10828-028 | -20°C<br>+4°C | M |
| mTeSR™:<br>1) mTeSR™ Basal medium<br>2) mTeSR™ 1.5X Supplement | STEMCELL Technologies | 05850 | +4°C<br>-20°C | M |
| 2-Mercaptoethanol | MP Biomedicals | 194705 | RT | N/A |
| TrypLE Express (1X) | Life Technologies | 12605010 | +4°C | M |
| ReLeSR™ | STEMCELL Technologies | 05872 | RT | M |
| Gelatin | Sigma | G1890 | RT | M |
| Matrigel | Corning | 254248 | -20°C<br>+4°C | M |
| Dulbecco's phosphate-buffered saline (DPBS) | Life Technologies | 14190-136 | +4°C | M |
| DMSO | Fisher Scientific | BP2311 | RT | M |
| FGF2 | R&D Systems | 223-FB-10 | -20°C<br>+4°C | M |
| Thiazovivin | Millipore | 420220 | -20°C<br>+4°C | M |
| Irradiated MEFs (Mouse Embryonic Fibroblasts) | Global Stem | GSC-6001G | Liquid Nitrogen | M |
| mRNA Reprogramming Kit | Stemgent | 00-0071 | -70°C | M |
| microRNA Booster Kit | Stemgent | 00-0073 | -70°C | M |
| B18R | Stemgent | 03-0071 | -70°C | M |
| Pluriton Supplement (2500x) | Stemgent | 01-0061 | -80°C | M |
| Pluriton Medium (500 mL) | Stemgent | 01-0015 | -20°C<br>+4°C | M |
| Stemfect RNA Transfection Kit:<br>1) Stemfect Transfection Buffer<br>2) Stemfect Transfection Reagent | Stemgent | 00-0069 | +4°C | M |

| | | | | |
|---|---|---|---|---|
| PureLink™ Genomic DNA Mini Kit | Life Technologies | K1820-001 | RT | N/A |
| e-Myco™ plus Mycoplasma PCR Detection KIT | iNtRON Biotechnology | 25235 | -20°C | M |
| 50 ml conical tube | BD FALCON | 352098 | RT | N/A |
| 15 ml conical tube | BD FALCON | 352099 | RT | N/A |
| 75 $cm^2$ Cell Culture Flask | Corning | 430641 | RT | N/A |
| 50 mL Rapid Flow Conical Filter with a 0.2 µm aPES membrane | Thermo Scientific | 564-0020 | RT | N/A |
| 6 well Tissue Culture (TC) plate | Corning | 353046 | RT | N/A |
| 2 ml Aspirating pipettes | BD FALCON | 357558 | RT | N/A |
| 5 ml Serological Pipettes | BD FALCON | 357543 | RT | N/A |
| 10 ml Serological Pipettes | BD FALCON | 357551 | RT | N/A |
| Hemocytomter | Thermo Scientific | 0267110 | RT | N/A |
| 1 mL filter tips | USA Scientific | 1126-7810 | RT | N/A |
| 200 µL filter tips | USA Scientific | 1120-8810 | RT | N/A |
| 20 µL filter tips | USA Scientific | 1123-1810 | RT | N/A |
| 0.1-10 µL filter tips | USA Scientific | 1121-3810 | RT | N/A |
| Cryovials | Nunc | 377367 | RT | N/A |
| Cryo 1°C freezing container | Nalgene | 5100-0001 | RT | N/A |
| 2-Propanol | Thermo Scientific | A417-4 | RT | N |

M; according to manufacture's shelf-life information; RT, room temperature

2. **Subject(s):** subject ID (de-identified), age, gender/sex, race/ethnicity

3. **Fibroblast(s):** subject ID (de-identified), passage number, dates of each passage, date of biopsy, date of initial plating, date of freezing.

4. **Microscopy pictures:** subject ID (de-identified), passage number, date, microscope name (company, catalogue number), magnification.

#### Quality Assurance/Control Steps (QA/QC)

QA/QC1: **Mycoplasma test** (relates to *Step 4*).

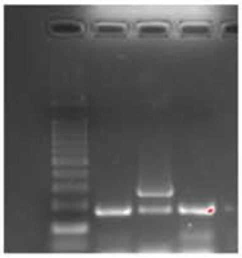

The above gel image. PCR products were separated on a 2% agarose gel. Lane 1, Bench Top 100 bp DNA ladder (Promega, Cat# G829B); Lane 2, negative/water control; Lane 3, positive control DNA (supplied with the e-Myco™ plus Mycoplasma PCR Detection KIT); Lane 4, DNA from actual fibroblast sample (100 ng of genomic DNA was used to run the mycoplasma PCR). The lower band is an internal control band indicating that the PCR worked.

QA/QC2: **Cell density of mycoplasma-free fibroblasts on day of reprogramming** (relates to *Step 6*).

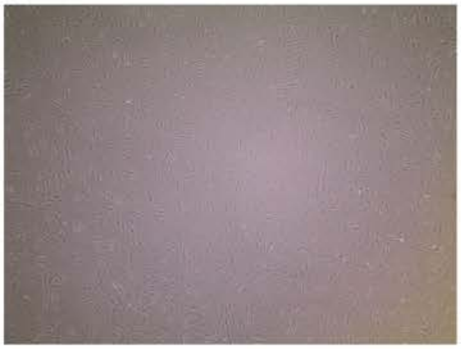

The above image is representative of the density of an established, mycoplasma-free fibroblast line taken on Day 1 just before start of reprogramming. Fifty thousand cells were plated in one well of a six-well plate the day before taking the picture.

QA/QC3: **Morphology of an emerging hiPSC colony** (relates to *Step 6*).

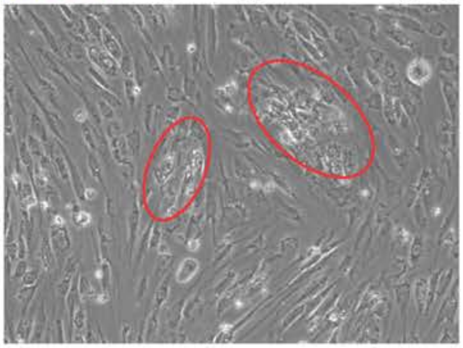

The above image represents the morphology of emerging hiPSC colonies (circles) that still require additional days of reprogramming.

**QA/QC4: Morphology of an established hiPSC line (relates to Step 7).**

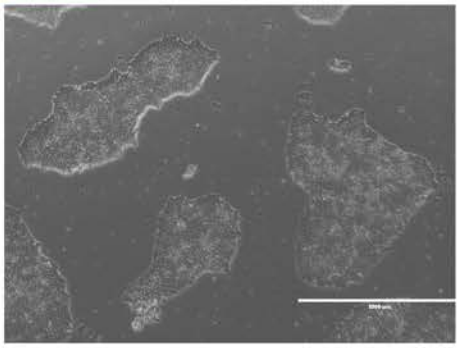

The above image represents the morphology of an established hiPSC line. Note the clear, focused edges of the colonies, which is typical for human pluripotent stem cells.

Icahn School of Medicine at Mount Sinai LINCS Center for Drug Toxicity Signatures

#### Standard Operating Procedure: Reprogramming Human Fibroblasts to Human Induced Pluripotent Stem Cells (hiPSCs) Using the Sendai Virus Method

DToxS SOP Index: CE-7.1
Last Revision: 01/07/2019
Written By: Christoph Schaniel and Sunita L. D'Souza

Approvals (Date): Joseph Goldfarb (01/07/19)
Marc Birtwistle (01/07/19)
Ravi Iyengar (10/27/20)

Quality Assurance/Control (QA/QC) steps are indicated with green highlight.

Metadata recording is highlighted with yellow highlight and superscript indices.

---

**Note: All materials/reagents are listed in a table in the Metadata[1] section of this SOP.**

1. Assemble media and reprogramming reagents
   Media and reagents are stored in original bottles of main ingredient, or if filtered, in bottles that are part of the filter set

   a. 0.1% Gelatin: Weigh 1 g of gelatin[1] and dissolve in 1L DPBS[1]. Autoclave and sterile filter using a 500 mL Rapid Flow Conical Filter with a 0.2 µm aPES membrane.[1]. Store at +4˚C. Keep for up to 4 weeks.
   b. Matrigel/DMEM: Thaw Matrigel on ice in a +4˚C cold room overnight. Under sterile conditions and on ice, add equal volume of DMEM[1] to the thawed Matrigel solution[1]. Pipette up and down several times to mix. Aliquot 2mL/15 mL conical tube[1]. Freeze aliquots at -20˚C. Keep for up to 4 months at -20˚C or up to 4 weeks at +4˚C.
   c. Complete DMEM/20% FBS (500 mL): Mix under sterile conditions 380 mL of DMEM[1], 5 mL Penicillin/Streptomycin (100x)[1], 5 mL Non-Essential Amino Acids (100x)[1], 5 mL L-Glutamine (200 mM)[1], 5 mL Sodium Pyruvate (100 mM)[1], 3.5 µL 2-mercaptoethanol (14.3 M)[1], and 100 mL FBS[1]. Store at +4˚C. Keep for up to 4 weeks.
   d. Complete DMEM/20% FBS without antibiotics (500 mL): Mix under sterile conditions 385 mL of DMEM[1], 5 mL Non-Essential Amino Acids (100x)[1], 5 mL L-Glutamine (200 mM)[1], 5 mL Sodium Pyruvate (100 mM)[1], 3.5 µL 2-mercaptoethanol (14.3 M)[1], 100 mL FBS[1]. Store at +4˚C. Keep for up to 4 weeks.
   e. Complete DMEM/10% FBS (100 mL): Mix under sterile conditions 86 mL DMEM[1], 1 mL Penicillin/Streptomycin (100x)[1], 1 mL Non-Essential Amino Acids (100x)[1], 1 mL L-Glutamine (200 mM)[1], 1 mL Sodium-Pyruvate (100 mM)[1], 0.7 µL 2-mercaptoethanol[1] (14.3 M) in DMEM, 10 mL FBS[1]. Store at +4˚C. Keep for up to 4 weeks.

f. Pluripotent stem cell (PSC) medium (500 mL): Under sterile conditions
   i. Supplement DMEM/F12 1:1 + L-glutamine +15 mM HEPES (henceforth referred to as DMEM/F12) (500 mL)[1] with 5 mL Penicillin/Streptomycin (100x)[1] and 5 mL L-Glutamine (200 mM)[1].
   ii. To 400 mL Supplemented DMEM/F12, add 100 mL KnockOut Serum Replacement,[1] 5 mL Non-Essential Amino Acids (100x)[1], 350 µL of a 1:100 dilution of 2-mercaptoethanol[1] in DMEM/F12, and sterile filter.[1]
   iii. Add 100 uL FGF2/BSA[1] [100 µg/mL] (final concentration: 20 ng/mL).
   iv. Wrap bottle in aluminum foil to protect from light at all times. Store at +4˚C. Keep for up to 4 weeks.
g. Complete mTeSR medium: Mix, under sterile conditions, 400 mL of mTeSR[1] with 100 mL mTeSR 1.5X Supplement[1] and 5 mL Penicillin/Streptomycin (100x)[1].
h. Thiazovivin [10 mM]: Under sterile conditions
   i. Suspend 10 mg Thiazovivin powder[1] in 3.212 mL of DMSO to obtain a 10 mM stock.
   ii. Place 50 µL aliquots into individual sterile, low protein binding 0.5 mL microcentrifuge tubes[1].
   iii. Freeze and store at -20˚C until use. Keep for up to 6 months.
i. FGF2/BSA: Reconstitute FGF2[1] at 100 µg/mL in sterile PBS containing 0.1% (0.1 g / 100mL PBS) bovine serum albumin. Aliquot at 100 µL per sterile 1.7 mL microtube[1] and store at -80˚C for up to 12 months. Store at +4˚C and keep up to 4 weeks.
j. hiPSC freezing medium A for cells on Mouse embryonic feeder cells (MEFs)[1] (50 mL): 25 mL PSC medium (*Step 1.f*), 20 mL FBS[1], 5 mL DMSO[1]. Store at +4˚C. Keep for up to 1 week.
k. hiPSC freezing medium B for cells on Matrigel (50 mL): 25 mL complete mTeSR medium (*Step 1.g*), 20 mL FBS[1], 5 mL DMSO[1]. Store at +4˚C. Keep for up to 1 week.
l. SeV Reprogramming cocktail:

The CytoTune™-iPS 2.0 Sendai Reprogramming Kit contains three CytoTune™ 2.0 reprogramming vectors that are used for delivering and expressing key genetic factors necessary for reprogramming somatic cells into iPSCs.

Contents and storage:

| Component[1] | Cap color | Amount[2] | | Storage |
|---|---|---|---|---|
| | | A16517 | A16518 | |
| CytoTune™ 2.0 KOS | clear | 100 µL | 3 × 100 µL | –80°C |
| CytoTune™ 2.0 hc-Myc | white | 100 µL | 3 × 100 µL | |
| CytoTune™ 2.0 hKlf4 | red | 100 µL | 3 × 100 µL | |

The titer of each CytoTune™ 2.0 reprogramming vector is lot-dependent. For the specific titer of your vectors, refer to the Certificate of Analysis (CoA) available at www.thermofisher.com/cytotune. Search for the CoA by product lot number, which is printed on the vial.

Each vial contains 100 µL of one of the CytoTune™ 2.0 reprogramming vectors at a concentration of ≥$8x10^7$ cell infectious units/mL (CIU/mL).

Under sterile condition,

   i. Prepare a master mix of SeV for 150,000 cells/subject calculated on the basis of the viral titer and number of subjects whose cells are to be infected

      Required volume of virus (µL) = multiplicity of infection (MOI) x number of cells x titer of virus x $10^{-3}$

**Note 1:** We recommend initially performing the transductions with MOIs of 5, 5, and 3 (i.e., KOS MOI=5, hc-Myc MOI=5, hKlf4 MOI=3). These MOIs can be optimized for your application.

**Note 2:** Avoid re-freezing and thawing of the reprogramming vectors since viral titers can decrease dramatically with each freeze/thaw cycle.

2. Thawing of cryopreserved fibroblasts

**Critical! Cells from each individual subject (fibroblasts or induced pluripotent stem cells) are to be kept in a separate plate/dish and all steps described below are to be handled separately for each sample to avoid any sample mix-up and/or cross contamination.**

a. Prepare in advance using a separate 6-well plate per fibroblast line/subject (This is critical to minimize the risk of sample mix-up and cross-contamination!): Label each plate with the de-identified sample name, fibroblast passage number (passage number at time of freezing +1), and date (metadata also includes the date of freezing). Add 1 mL of 0.1% Gelatin (*Step 1.a.*) to each of 2 wells of a 6-well plate. Set the plate aside for 30-60 min. Aspirate gelatin solution and add 1 mL of complete DMEM/20% FBS (*Step 1.c.*) to one gelatin well (well 1) and 1 mL of complete DMEM/20% FBS without antibiotics (*Step 1.d.*) to the other gelatin well (well 2). Ensure that the entire surface of the well is covered with media. The antibiotic-free well will be used for mycoplasma testing.
b. Remove a cryovial with fibroblast passage number < 3 from the liquid nitrogen storage tank and keep on dry ice.
c. Add 250 µL of complete DMEM[1]/20% FBS[1] without antibiotics (*Step 1.d.*) to the cryovial and thaw at 37˚C for 1-3 minutes until only a tiny frozen cube remains.
d. Under sterile conditions, transfer cell suspension into a 15 mL conical tube filled with 4 mL complete DMEM[1]/20% FBS[1] without antibiotics (*Step 1.d.*).
e. Centrifuge tube at 300 g for 5 min at room temperature.
f. Aspirate medium, add 2 mL of complete DMEM[1]/20% FBS[1] without antibiotics (*Step 1.d.*), resuspend cells gently and add 1 mL per well (2 wells in total).
g. Place the plate(s) in a humidified incubator at 37°C / 5% $CO_2$.
h. Next day, replace medium with 2 mL fresh complete DMEM/20% FBS *with* antibiotics (*Step 1.c.*) for well 1, and 2 mL fresh complete DMEM/20% FBS *without* antibiotics for well 2.
i. Once cells are up to 80% confluent proceed to *Step 3* for the well with antibiotic-free medium (well 2; cells need to be cultured in antibiotic-free medium for at least 5 days for the mycoplasma test to be meaningful) and *Step 4* for the well with antibiotic-containing medium, if mycoplasma testing is negative.

3A. QA/QC1Mycoplasma testing of fibroblasts (luminescence based, immediate testing, no DNA isolation required)

a. Collect supernatant into a tube and centrifuge at 300 g for 10 min at room temperature in order to remove any dead cells & cell debris.
b. Transfer supernatant to a fresh tube. (Note: supernatant can be stored at +4˚C for up to one week before testing)
c. Take "Myco R", "Myco S" and "Myco + Control" vials from the MycoAlert Mycoplasma Detection Kit[1] stored at -80˚C and thaw them at room temperature.
d. Pipet 50 µL of supernatant (*Step 3A.b.*) into one well of a 96-well assay plate[1]. Also make negative (-) and positive (+) control wells. Use 50 µL of DMEM[1]/20% FBS[1] without antibiotics for the (-) control and for the (+) control, 2 µL of the "Myco + Control" from the kit.
e. Add 50 µL "Myco R" to each test sample and the (+) and (-) controls.
f. Incubate at room temperature for 5 min.
g. Take a luminescence reading on a plate reader equipped for luminometry (we use an Enspire Multilabel Plate reader from ThermoFisher Scientific). This will be reading "A".
h. Add 50 µL "Myco S" to each sample and the controls.
i. Incubate at room temperature for 10 min.
j. Take a luminescence reading on a plate reader equipped for luminometry (we use an Enspire Multilabel Plate Reader from ThermoFisher Scientific). This will be reading "B".
k. Calculate mycoplasma" load" by dividing reading “B” by reading "A" (B/A) The sample is mycoplasma free if the “B”/”A” ratio is <1.25.

l. Discard any positive cultures and start over.

3B. QA/QC1 - Alternative Mycoplasma testing of fibroblasts (PCR-based, requires DNA isolation)

a. Add 500 µL TrypLE Express (1x)[1] to well 2 and incubate for 3-5 minutes. Use a microscope to check when fibroblasts are rounding up or starting to lift off the bottom of the well.
b. Add 500 µL complete DMEM/20% FBS without antibiotics (*Step 1.d.*).
c. Harvest fibroblasts using a 1 mL micropipette tip and add cells into a 15 mL conical tube filled with 9 mL sterile DPBS[1].
d. Centrifuge tubes at 300 g for 5 minutes at 4˚C.
e. Aspirate medium and resuspend fibroblasts in 200 µL DPBS[1].
f. Proceed with DNA isolation using and following the Pure LINK™ Genomic DNA Mini Kit protocol (see Appendix A).
g. Proceed with Mycoplasma testing using and following the e-Myco™ plus Mycoplasma PCR Detection KIT according to the manufacturer's instructions (see Appendix B).
h. Discard any positive cultures and start over.

4. Harvesting of fibroblasts for reprogramming

a. On day 0 (D0), inspect cell density and morphology of mycoplasma-free fibroblasts under a microscope.
b. Aspirate medium from the remaining well containing fibroblasts at up to 80% confluence (*Step 2.i.*) and wash once with DPBS[1] (~1 mL/well).
c. Add 500 µL TrypLE[1] to the well and incubate for 3-5 minutes. Use a microscope to check when fibroblasts are rounding up and starting to lift off the bottom of the well.
d. Add 500 µL complete DMEM/20% FBS (*Step 1.c.*).
e. Harvest fibroblasts using a 1 mL micropipette tip and transfer to a 15 mL conical tube filled with 4 mL of complete DMEM/20% FBS (*Step 1.c.*).
f. Use 10 µL from the tube with cells in complete DMEM/20% FBS to determine the total number of fibroblasts with a hemocytometer.

5. Reprogramming of fibroblasts

**Critical: Reprogram only those fibroblasts/wells that are mycoplasma-free (step 3)!**

**Note: The amounts are for reprogramming 1 well with fibroblasts in a 6-well plate.**
**All reprogramming and expansion steps that follow are performed in a sterile BSL-2 hood**
**Plates have lid on except when adding reagents or cells**

a. At least 30 minutes before reprogramming starts, coat one well of a 6-well plate per fibroblast line (separate plates per line to minimize contamination/mix-up) with Matrigel (*Step 1.b.*) diluted in DMEM to a final ratio of 1:100 v/v (1:50 dilution of Matrigel/DMEM mix from *Step 1.b.*). Use 1 mL for each well of a 6-well plate. Note: Label plate with the de-identified sample name and date.
b. Remove one CytoTune™-iPS2.0 kit[1] (*Step 1.k.*) from -80°C storage. (One kit can be used to reprogram 8-9 wells/fibroblast lines).
c. Thaw each tube one at a time by first immersing the bottom of the tube in a 37°C water/bead bath for 5–10 seconds followed by thaw at room temperature. Once thawed, briefly centrifuge the tube and place it immediately on ice.
d. Check certificate of analysis from manufacturer (Fisher Scientific-Life Technologies) for number of Sendai viral particles (see *Step 1.k.*) and calculate appropriate volume to be added in *Step 5.f.*
e. Warm up complete DMEM/20% FCS (*Step 1.d.*)(1 mL per fibroblast line/well to be infected) at 37˚C.
f. Mix 150,000 fibroblast cells (*Step 4.e.*) in 1mL DMEM/20% FCS media with an appropriate volume

of SeV particles and add to the well.

g. Swirl the 1mL around to make sure that the virus is well distributed. Discard all virus tips into a 10% bleach solution.
h. Place the plate(s) in a humidified incubator at 37°C / 5% $CO_2$. Note: This is D0 of reprogramming.
i. After 24 hours of virus incubation (D1), feed with 1mL fresh complete DMEM/20% FBS.
j. D1: Check cells under a microscope (QA/QC2) and replace medium with fresh 2 mL of complete DMEM/20% FBS.
k. D2: Look at cells. Replace medium with fresh 2 mL of complete DMEM/20% FBS (QA/QC3).
l. D3: Replace medium with fresh 2 mL of complete DMEM/20% FBS.
m. D4: Replace medium with fresh 2 mL of complete DMEM/20% FBS.
n. D5: Replace medium with fresh 2 mL of complete DMEM/20% FBS.
o. D6: Replace medium with fresh 2 mL of complete DMEM/20% FBS.
p. D7: Check cells under a microscope (QA/QC4) and replace medium with fresh 2 mL of complete DMEM/20% FBS. Go to Step 6.

6. Passaging of SeV-infected fibroblasts onto mouse embryonic feeders

a. D7: Prepare 3 100-mm standard tissue culture dishes of irradiated mouse embryonic fibroblasts[1] (MEFs), 1 million per dish per fibroblast line/subject.
   i. Add 10 mL of 0.1% Gelatin per dish. Set the dishes aside for 30-60 min.
   ii. Remove MEF cryovial from the liquid nitrogen storage tank and keep on dry ice.
   iii. Add 250-500 µL of complete DMEM/10% FBS(step 1.e) to the cryovial and thaw at 37˚C (do not thaw completely; leave a tiny frozen cube behind).
   iv. Transport cryovial to a sterile tissue culture hood and transfer MEFs into a 50 mL conical tube filled with 4 mL complete DMEM/10% FBS.
   v. Centrifuge tube at 1000 rpm for 5 min at room temperature.
   vi. Aspirate medium, add 40 mL of complete DMEM/10% FBS and resuspend MEF cells.
   vii. Aspirate 1% Gelatin from dishes and add 10 mL of MEFs per 100-mm dish.
   viii. Place the dishes in a humidified incubator at 37°C / 5% $CO_2$.
b. D8: Harvest "reprogrammed" fibroblasts from *Step 5.p.*:
   i. Aspirate medium from well(s) containing "reprogrammed" fibroblasts at 80-100% confluence and wash once with DPBS (~1 mL/well).
   ii. Add 500 µL TrypLE per well and incubate for 3-5 minutes. Use a microscope to check when "reprogrammed" fibroblasts are rounding up or starting to lift off the bottom of the well.
   iii. Add 500 µL of DMEM/10% FBS per well.
   iv. Harvest "reprogrammed" fibroblasts using a 1 mL pipette with 1 mL tip and add cells into a 15 mL conical tube filled with 4 mL of complete DMEM/10% FBS.
c. Centrifuge tubes at 1000 rpm for 5 minutes at room temperature
d. Aspirate medium and resuspend "reprogrammed" fibroblasts in 5 mL DMEM/10% FBS.
e. Use 10 µL of the cell suspension to determine the total number of "reprogrammed" fibroblasts with a hemocytometer. (Cells should have undergone a 7-10-fold expansion).
f. Plating of "reprogrammed" fibroblasts onto MEF dishes
   i. Label dishes (from *Step 6.a.viii*) with the de-identified sample name, and date (METADATA also includes the fibroblast passage number, date of freezing & thawing, date of reprogramming) of the "reprogrammed" fibroblasts to be plated.
   ii. Aspirate the medium from the 1$^{st}$ 100-mm dish (*Step 6a.viii.*) and plate 50,000 "reprogrammed" fibroblasts onto the dish in 10 mL DMEM/10% FBS.
   iii. Aspirate the medium from the 2$^{nd}$ 100-mm dish (*Step 6a.viii.*) and plate 100,000 "reprogrammed" fibroblasts onto the dish in 10 mL DMEM/10% FBS.
   iv. Aspirate the medium from the 3$^{rd}$ 100-mm dish (*Step 6a.viii.*) and plate 150,000 "reprogrammed" fibroblasts onto the dish in 10 mL DMEM/10% FBS.
   v. Transfer remaining "reprogrammed" fibroblasts to cryovials--500,000 cells/mL/vial in 50%DMEM/40% BFS/10%DMSO. Freeze using a Cryo 1˚C freezing container[1](filled with

250 mL 2-Propanol[1]) at -80˚C overnight then move vials to a liquid nitrogen tank for long-term storage.

g. D9: Replace 2.5 mL of medium in each dish with fresh PSC medium (*Step 1.f.*).
h. D10: Replace 5.0 mL of medium in each dish with fresh PSC medium.
i. D11: Replace 7.5 mL of medium in each dish with fresh PSC medium.
j. D12: Replace all medium in each dish with 10 mL fresh PSC medium.
k. D13: Replace all medium in each dish with 10 mL fresh PSC medium
l. D14: Replace all medium in each dish with 10 mL fresh PSC medium.
m. D15+: Replace all medium in each dish with 10 mL fresh PSC medium on a daily basis, until hiPSC colonies (clones) reach > 1,000 cells (QA/QC5). Then proceed to step 7.

7. Picking and replating induced pluripotent stem cells on MEFs

a. When hiPSC clones reach >1,000 cells (QA/QC5) prepare one 48-well plate of MEFs (1/48th of 1 million MEFs per well, 1 million MEFs per plate if all wells were used) per subject.
   i. Label plate with the de-identified subject ID and date. (METADATA also should include the fibroblast passage number, date of freezing & thawing, date of reprogramming, date of picking).
   ii. Add 250 µL of 0.1% Gelatin (*Step 1.a.*) per well and set the plate(s) aside for 30-60 min at room temperature. Coat only 16 wells / 48-well plate.
   iii. Remove MEF cryovial from the liquid nitrogen storage tank and keep on dry ice.
   iv. Add 250-500 µL of DMEM/10% FBS (*Step 1.e.*) to the cryovial and thaw at 37˚C (do not thaw completely; leave a tiny frozen cube behind).
   v. Transport cryovial to a sterile tissue culture hood and transfer MEFs into a 50 mL conical tube filled with 4 mL DMEM/10% FBS.
   vi. Centrifuge tube at 1000 rpm for 5 min at room temperature.
   vii. Aspirate medium, add 48 mL of DMEM/10% FBS and resuspend cells.
   viii. Aspirate 0.1% Gelatin from wells and add 250 µL of MEFs to each of the 16 wells in a 48-well plate
   ix. Place the plate(s) in a humidified incubator at 37°C / 5% $CO_2$.
b. The following day, aspirate medium from wells and replace with 250 µL of PSC medium (*Step 1.f.*) supplemented with Thiazovivin at a final concentration of 2 µM (*Step 1.h*; 1:5,000 dilution) per well.
c. Pick individual clones using physical scraping and sucking into a 20 uL filter tip with the help of an inverted microscope (in the hood) and place each picked clone into an individual well. This is considered passage 1. Note, sample name is now MSN_subject ID code-N, where N is the two-digit clone number followed by the letter S (S stands for Sendai Virus reprogramming) (e.g., MSN01-01S).
d. The next day, and on subsequent days, replace medium with 250 µL fresh PSC medium (*Step 1.f.*) without Thiazovivin per well until clones reach at least 500-1000 cells and retain an undifferentiated morphology (QA/QC5). (Go to *Step 8*).

8. Splitting/expansion of induced pluripotent stem cells on MEFs

a. Cells are expanded from 48-well format to 24-well format
   i. When hiPSC clones reach about 70% confluency, prepare one well of a 24-well plate of MEFs per iPSC clone of the 48-well plate that reaches criterion size and morphology, according to *Step 7.a.i.-ix*, but in step viii add 500 µL of MEFs to each well.
   ii. The following day, harvest hiPSCs from MEF plate by washing with 150 µL/well of calcium- and magnesium-free phosphate-buffered saline (DPBS)
   iii. Add 150 µL/well of ReLeSR™[1] and aspirate ReLeSR™[1] within one minute, so that colonies are exposed to a thin film of liquid.
   iv. Incubate the culture at 37°C for 5 minutes.

v. Prepare PSC medium (*Step 1.f.*) supplemented with Thiazovivin at a final concentration of 2 µM (*Step 1.h*; 1:5,000 dilution). Add 125 µL/well of complete PSC medium plus Thiazovivin (only for day 1 of splitting).
vi. Detach the colonies by holding the plate with one hand and use the other hand to firmly tap the side of the plate for approximately 30 - 60 seconds.
vii. Once the cell aggregates start lifting off, remove cells and re-plate the cell aggregate mixture from the harvest well into 1 well of the 24 well plate with MEFs .
    Note:,aspirate medium from MEF-containing wells prior to adding PSC medium with iPSCs.
viii. Place the plate in a 37°C incubator. Move the plate in several quick, short, back-and-forth and side-to-side motions to evenly distribute the cell aggregates. Do not disturb the plate for 12-24 hours.
ix. Perform daily medium changes, aspirating old medium and replacing with complete PSC medium (no Thiazovivin); 500 µL/well (and visually assess cultures to monitor growth until the next passaging time.

b. Cells are expanded from 24-well format to 12-well format
i. When hiPSC clones reach about 70% confluency, prepare one well of a 12-well plate of MEFs per iPSC clone of the 24-well plate that reaches criterion size and morphology, according to *Step 7.a.i.-ix*, but in step viii add 1 mL of MEFs to each well.
ii.The following day, harvest hiPSCs from MEF plate by washing with, 250 µL/well of calcium- and magnesium-free phosphate-buffered saline (DPBS)[1].
iii. Add 250 µL/well of ReLeSR™[1] and aspirate ReLeSR™[1] within one minute, so that colonies are exposed to a thin film of liquid.
iv. Incubate the culture at 37°C for 5 minutes.
v. Prepare PSC medium (*Step 1.f.*) supplemented with Thiazovivin at a final concentration of 2 µM (*Step 1.h*; 1:5,000 dilution). Add 250 µL/well of complete PSC medium plus Thiazovivin (only for day 1 of splitting).
vi. Detach the colonies by holding the plate with one hand and use the other hand to firmly tap the side of the plate for approximately 30 - 60 seconds.
vii. Once the cell aggregates start lifting off, remove cells and re-plate the cell aggregate mixture from the harvest well into 1 well of the 12 well plate with MEFs
    Note: aspirate medium from MEF-containing wells prior to adding PSC medium with iPSCs.
viii. Place the plate in a 37°C incubator. Move the plate in several quick, short, back-and-forth and side-to-side motions to evenly distribute the cell aggregates. Do not disturb the plate for 12-24 hours.
ix. Perform daily medium changes, aspirating old medium and replacing with complete PSC medium (no Thiazovivin) 1 mL/well and visually assess cultures to monitor growth until the next passaging time.

c. Cells are expanded from 12-well format to 6-well format
i. When hiPSC clones reach about 70% confluency, prepare one well of a 6-well plate of MEFs per iPSC clone of the 12-well plate that reaches criterion size and morphology, according to *Step 7.a.i.-ix*, but in step viii add 2 mL of MEFs to each well.
ii. The following day, harvest hiPSCs from MEF plate by washing with, 500 µL/well of calcium- and magnesium-free phosphate-buffered saline (DPBS)[1].
iii. Add 500 µL/well of ReLeSR™[1] and aspirate ReLeSR™[1] within one minute, so that colonies are exposed to a thin film of liquid.
iv. Incubate the culture at 37°C for 5 minutes.
v. Prepare PSC medium (*Step 1.f.*) supplemented with Thiazovivin at a final concentration of 2 µM (*Step 1.h*; 1:5,000 dilution). Add 500 µL/well of complete PSC medium plus Thiazovivin (only for day 1 of splitting).
vi. Detach the colonies by holding the plate with one hand and use the other hand to firmly tap

the side of the plate for approximately 30 - 60 seconds.

vii. Once the cell aggregates start lifting off, remove cells and re-plate the cell aggregate mixture from the harvest well into 1 well of the 6 well plate with MEFs

Note: aspirate medium form MEF-containing wells prior to adding PSC medium with iPSCs.

viii. Place the plate in a 37°C incubator. Move the plate in several quick, short, back-and-forth and side-to-side motions to evenly distribute the cell aggregates. Do not disturb the plate for 12-24 hours.

ix.Perform daily medium changes, aspirating old medium and replacing with complete PSC medium (no Thiazovivin); 2 mL/well and visually assess cultures to monitor growth until the hiPSCs reach 80% confluence.

d. Expand each well of the 6 well plate into 3 wells of a 6-well MEF plate.

i. When hiPSC clones reach about 70% confluency, prepare three wells of a 6-well plate of MEFs per well of the 6-well plate that reaches criterion size and morphology, according to *Step 7.a.i.-ix*, but in step viii add 2 mL of MEFs to each well.

ii. The following day, harvest hiPSCs from MEF plate by washing with, 500 µL/well of calcium- and magnesium-free phosphate-buffered saline (DPBS)[1].

iii. Add 500 µL/well of ReLeSR™[1] and aspirate ReLeSR™[1] within one minute, so that colonies are exposed to a thin film of liquid.

iv. Incubate the culture at 37°C for 5 minutes.

v. Prepare PSC medium (*Step 1.f.*) supplemented with Thiazovivin at a final concentration of 2 µM (*Step 1.h*; 1:5,000 dilution). Add 500 µL/well of complete PSC medium plus Thiazovivin (only for day 1 of splitting).

vi. Detach the colonies by holding the plate with one hand and use the other hand to firmly tap the side of the plate for approximately 30 - 60 seconds.

vii. Once the cell aggregates start lifting off, remove cells and re-plate the cell aggregate mixture from the harvest well into 3 wells of a 6 well plate with MEFs

Note: aspirate medium form MEF-containing wells prior to adding PSC medium with iPSCs.

viii. Place the plate in a 37°C incubator. Move the plate in several quick, short, back-and-forth and side-to-side motions to evenly distribute the cell aggregates. Do not disturb the plate for 12-24 hours.

ix.Perform daily medium changes, aspirating old medium and replacing with complete PSC medium (no Thiazovivin); 2 mL/well and visually assess cultures to monitor growth until the hiPSCs reach 80% confluence.

x. Once hiPSCs reach 70% confluence, split one well at a 1 to 6 split ratio (i.e., 1 well into 6 wells) onto Matrigel-coated plates (see *Step 9*) and cryopreserve the other two wells in separate cryovials (see *Step 10.*).

9. Feeder-free splitting/expansion of induced pluripotent stem cells onto Matrigel-coated plates)

a. At least 30 minutes before passaging hiPSCs from MEFs onto Matrigel-plates, dilute Matrigel (*Step 1.b.*) in DMEM at a final ratio of 1:60 v/v (1:30 dilution of Matrigel/DMEM mix from *Step 1.b.*). Use 1 mL for each well of a 6-well plate. Note: Label plate with the de-identified sample name and date. (METADATA also should include the passage number, date of freezing & thawing, date of reprogramming, date of picking).

b. Wash cells on MEF plate with 1 mL/well (6-well format) of calcium- and magnesium-free phosphate-buffered saline (DPBS)[1] and aspirate.

c. Add 1 mL/well of ReLeSR™[1] and aspirate ReLeSR™[1] within one minute, so that colonies are exposed to a thin film of liquid.

d. Incubate the culture at 37°C for 5 minutes.

e. Prepare complete mTeSR medium (*Step 1.g.*) supplemented with Thiazovivin (*Step 1.h.*) at a final concentration of 2 μM (*Step 1.h*; 1:5,000 dilution). . Add 1 mL/well of complete mTeSR medium + Thiazovivin (only for day 1 of splitting).

f. Detach the colonies by holding the plate with one hand and use the other hand to firmly tap the side of the plate for approximately 30 - 60 seconds.
g. Once the cell aggregates start lifting off, remove cells and re-plate the cell aggregate mixture from one MEF well into the 6 wells of a 6 well plate pre-coated with Matrigel (*Step 9.a.*).
h. Place the plate in a 37°C incubator. Move the plate in several quick, short, back-and-forth and side-to-side motions to evenly distribute the cell aggregates. Do not disturb the plate for 24 hours.
i. Perform daily medium changes, aspirating old medium and replacing with complete mTeSR medium (no Thiazovivin; 2 ml/well of a 6 well plate) and visually assess cultures to monitor growth until the next passaging time.

10. Expansion (6 well plate format) of about 80% confluent wells and cryopreservation/freezing of induced pluripotent stem cells

a. Prepare in advance:
   i. Fresh hiPSC freezing medium A or B depending on whether cells are frozen from plating on MEFs(A) or Matrigel (B) pre-warmed to 37˚C (*Steps 1.j, 1.k.*).
   ii. Cryovials: label 6 vials with "MSN_patient de-identification code-N, where N is the colony/clone number followed by the letter S (for Sendai virus reprogramming) (e.g., passage number (p), and date of freezing (e.g., MSN01-01S, p10, 04052016).
b. Split 1/6 of a well of a 6 well plate into a fresh well of a 6 well plate following *Steps 8a-8i,* by washing cells with 1mL/well of calcium- and magnesium-free phosphate-buffered saline (DPBS)[1] using 1 mL/well of ReLeSR™[1] , but instead of transferring 1 mL only transfer 166 μL to one well of a Matrigel/DMEM coated 6 well plate in 1 mL complete mTeSR medium (*Step 1.g.*) supplemented with Thiazovivin (*Step 1.h.*) at a final concentration of 2 μM (*Step 1.h*; 1:5,000 dilution).. These cells can continue to be expanded for subsequent experiments. Track passage number.
c. The remaining cells in the well, and the cells in the other 5 wells of the Matrigel plate are cryopreserved at this point as a precaution to preserve the cell line in the event of unexpected problems during the expansion phase.
   i. Transfer the 834 μL of hiPSCs into a 15 mL conical tube.
   ii. Centrifuge at 300 g for 5 min at 4˚C.
   iii. Aspirate medium and resuspend cells in 1 mL of of hiPSC freezing medium B (*Step 1.k.*).
   iv. Transfer cells to the cryovial and freeze using a Cryo 1˚C freezing container[1] (filled with 250 mL 2-Propanol[1]) at -80˚C overnight then move vials to a liquid nitrogen tank for long-term storage.

**Note: When iPSC passage number reaches 9 expand cells to at least 6 wells of 6-well plate format (iSPC characterization which includes karyotyping and STR analysis is done at passage 10+). Once karyotype is found normal, cells are further expanded for 1-2 rounds and cryopreserved/banked at ≥10 vials per clone.**

**Metadata**

1. **Materials/Reagents:** Company name, Catalogue and lot numbers

| PRODUCT | COMPANY NAME | CAT # | Storage | Usable life |
|---|---|---|---|---|
| DMEM (500 mL) | Thermo Fisher Scientific | 11965-118 | +4˚C | M |
| IMDM | Thermo Fisher Scientific | 12440-053 | +4˚C | M |
| DMEM/F12 (DMEM/F12 (1:1) + L-Glutamine + 15mM HEPES) | Thermo Fisher Scientific | 11330-032 | +4˚C | M |
| Penicillin/Streptomycin (100x) | Thermo Fisher Scientific | 15140-122 | +4˚C | M |
| Sodium-Pyruvate (100 mM) | Thermo Fisher Scientific | 11360-070 | +4˚C | M |
| L-Glutamine (200 mM) | Thermo Fisher Scientific | 25030-081 | +4˚C | M |
| Non-Essential AminoAcids (100x) | Thermo Fisher Scientific | 11140-050 | +4˚C | M |
| Fetal Bovine Serum (FBS; 500 mL) | Corning | 35-011-CV | +4˚C | M |
| EDTA | Corning | 46-034-CI | RT | M |
| KnockOut Serum Replacement | Thermo Fisher Scientific | 10828-028 | -20˚C<br>+4˚C | M |
| mTeSR™:<br>1) mTeSR™ Basal medium<br>2) mTeSR™ 1.5X Supplement | STEMCELL Technologies | 05850 | +4˚C<br>-20˚C | M |
| 2-Mercaptoethanol | MP Biomedicals | 194705 | RT | N/A |
| TrypLE Express (1X) | Thermo Fisher Scientific | 12605010 | +4˚C | M |
| ReLeSR™ | STEMCELL Technologies | 05872 | RT | M |
| Gelatin | Sigma | G1890 | RT | M |
| Matrigel | Corning | 254248 | -20˚C<br>+4˚C | M |
| Dulbecco's Phosphate-Buffered Saline (DPBS) | Thermo Fisher Scientific | 14190-136 | +4˚C | M |
| DMSO | Fisher Scientific | BP2311 | RT | M |
| FGF2 | R&D Systems | 223-FB-10 | -20˚C<br>+4˚C | M |
| Thiazovivin | Millipore | 420220 | -20˚C<br>+4˚C | M |
| Irradiated MEFs (Mouse Embryonic Fibroblasts) | Global Stem | GSC-6001G | Liquid Nitrogen | M |
| CytoTune™-iPS 2.0 Sendai Reprogramming Kit | Thermo Fisher Scientific | A16517 | -80˚C | M |
| MycoAlert Mycoplasma Detection Kit | Lonza Walkersville | LT07218 | -80˚C | M |
| PureLink™ Genomic DNA Mini Kit | Thermo Fisher Scientific | K1820-001 | RT | N/A |
| e-Myco™ plus Mycoplasma PCR Detection KIT | iNtRON Biotechnology | 25235 | -20˚C | M |
| 1.7 mL microtube | Thermo Fisher Scientific | 14-222-168 | RT | N/A |
| 50 ml conical tube | BD FALCON | 352098 | RT | N/A |
| 15 ml conical tube | BD FALCON | 352099 | RT | N/A |
| 500 mL Rapid Flow Conical Filter with a 0.2 µm aPES membrane | Thermo Fisher Scientific | 566-0020 | RT | N/A |

| | | | | |
|---|---|---|---|---|
| 100 mm standard tissue culture dish | Corning Falcon | 353003 | RT | N/A |
| 96 well assay plate, with lid | Costar | 3917 | RT | N/A |
| 6 well Tissue Culture (TC) plate | Corning Falcon | 353046 | RT | N/A |
| 2 ml Aspirating pipettes | BD FALCON | 357558 | RT | N/A |
| 5 ml Serological Pipettes | BD FALCON | 357543 | RT | N/A |
| 10 ml Serological Pipettes | BD FALCON | 357551 | RT | N/A |
| Hemocytomter | Thermo Fisher Scientific | 0267110 | RT | N/A |
| 1 mL filter tips | USA Scientific | 1126-7810 | RT | N/A |
| 200 µL filter tips | USA Scientific | 1120-8810 | RT | N/A |
| 20 µL filter tips | USA Scientific | 1123-1810 | RT | N/A |
| 0.1-10 µL filter tips | USA Scientific | 1121-3810 | RT | N/A |
| Cryovials | Nunc | 377367 | RT | N/A |
| Cryo 1˚C freezing container | Nalgene | 5100-0001 | RT | N/A |
| 2-Propanol | Thermo Fisher Scientific | A417-4 | RT | N |

M; according to manufacture's shelf-life information; RT, room temperature

2. **Subject(s):** subject ID (de-identified), age, gender/sex, race/ethnicity

3. **Fibroblast(s):** subject ID (de-identified), passage number, dates of each passage, date of biopsy, date of initial plating, date of freezing.

4. **Microscopy pictures:** subject ID (de-identified), passage number, date, microscope name (company, catalogue number), magnification.

#### Quality Assurance/Control Steps (QA/QC)

**QA/QC1: Mycoplasma test** (relates to *Step 3B*).

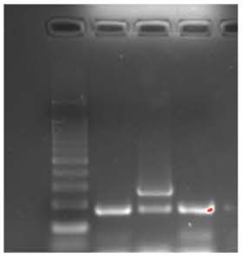

The above gel image. PCR products were separated on a 2% agarose gel. Lane 1, Bench Top 100 bp DNA ladder (Promega, Cat# G829B); Lane 2, negative/water control; Lane 3, positive control DNA (supplied with the e-Myco$^{TM}$ plus Mycoplasma PCR Detection KIT); Lane 4, DNA from actual fibroblast sample (100 ng of genomic DNA was used to run the mycoplasma PCR). The lower band is an internal control band indicating that the PCR worked.

**QA/QC2 (visual, not recorded): Cell density of mycoplasma-free fibroblasts one day after reprogramming** (relates to *Step 5.j*).

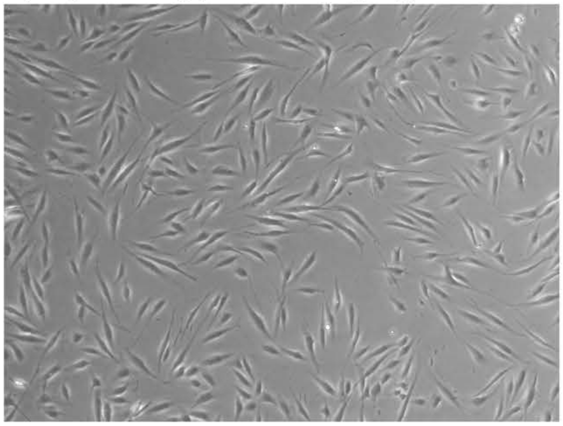

The above image is representative of the density of an established, mycoplasma-free fibroblast line taken on Day 1 of reprogramming. One hundred fifty thousand cells were plated in one well of a six-well plate the day before taking the picture.

**QA/QC3 (visual, not recorded): Morphology on day 2 of reprogramming** (relates to *Step 5.k.*)

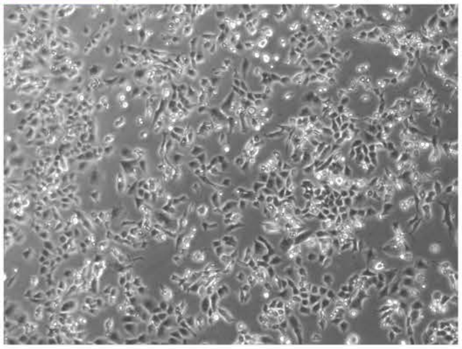

**QA/QC4 (visual, not recorded): Morphology before splitting reprogrammed fibroblasts onto MEFs** (relates to *Step 5.o*).

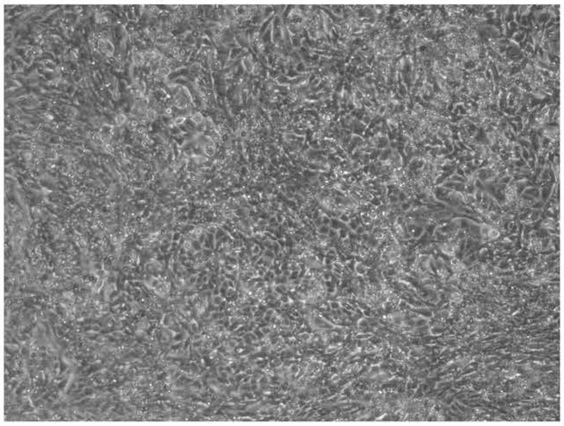

**QA/QC5 (visual, not recorded): Morphology at picking** (relates to *Step 7*).

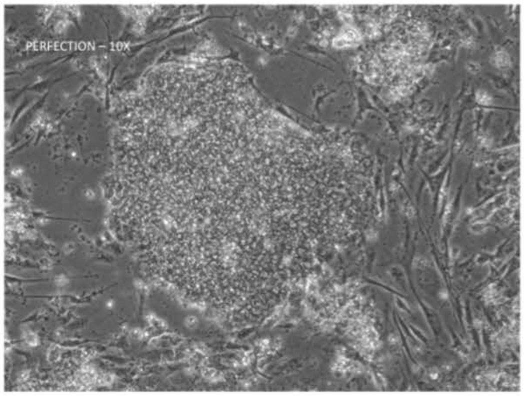

The above image represents the morphology of an established hiPSC line. Note the clear, focused edges of the colonies, which is typical for human pluripotent stem cells.

**References**

Chen, E.Y., Tan, C.M., Kou, Y., Duan, Q., Wang, Z., Meirelles, G.V., Clark, N.R., and Ma'ayan, A. (2013). Enrichr: interactive and collaborative HTML5 gene list enrichment analysis tool. BMC Bioinformatics *14*, 128.

D'Antonio-Chronowska, A., Donovan, M.K.R., Young Greenwald, W.W., Nguyen, J.P., Fujita, K., Hashem, S., Matsui, H., Soncin, F., Parast, M., Ward, M.C.*, et al.* (2019). Association of Human iPSC Gene Signatures and X Chromosome Dosage with Two Distinct Cardiac Differentiation Trajectories. Stem Cell Reports *13*, 924-938.

Geoffroy, V., Pizot, C., Redin, C., Piton, A., Vasli, N., Stoetzel, C., Blavier, A., Laporte, J., and Muller, J. (2015). VaRank: a simple and powerful tool for ranking genetic variants. PeerJ *3*, e796.

Hagai, T., Chen, X., Miragaia, R.J., Rostom, R., Gomes, T., Kunowska, N., Henriksson, J., Park, J.E., Proserpio, V., Donati, G.*, et al.* (2018). Gene expression variability across cells and species shapes innate immunity. Nature *563*, 197-202.

Kilpinen, H., Goncalves, A., Leha, A., Afzal, V., Alasoo, K., Ashford, S., Bala, S., Bensaddek, D., Casale, F.P., Culley, O.J.*, et al.* (2017). Common genetic variation drives molecular heterogeneity in human iPSCs. Nature *546*, 370-375.

Kuleshov, M.V., Jones, M.R., Rouillard, A.D., Fernandez, N.F., Duan, Q., Wang, Z., Koplev, S., Jenkins, S.L., Jagodnik, K.M., Lachmann, A.*, et al.* (2016). Enrichr: a comprehensive gene set enrichment analysis web server 2016 update. Nucleic Acids Res *44*, W90-97.

Landrum, M.J., Lee, J.M., Riley, G.R., Jang, W., Rubinstein, W.S., Church, D.M., and Maglott, D.R. (2014). ClinVar: public archive of relationships among sequence variation and human phenotype. Nucleic Acids Res *42*, D980-985.

Lazaridis, I., Nadel, D., Rollefson, G., Merrett, D.C., Rohland, N., Mallick, S., Fernandes, D., Novak, M., Gamarra, B., Sirak, K.*, et al.* (2016). Genomic insights into the origin of farming in the ancient Near East. Nature *536*, 419-424.

Li, H., and Durbin, R. (2009). Fast and accurate short read alignment with Burrows-Wheeler transform. Bioinformatics *25*, 1754-1760.

Li, H., Handsaker, B., Wysoker, A., Fennell, T., Ruan, J., Homer, N., Marth, G., Abecasis, G., Durbin, R., and Genome Project Data Processing, S. (2009). The Sequence Alignment/Map format and SAMtools. Bioinformatics *25*, 2078-2079.

Tohyama, S., Hattori, F., Sano, M., Hishiki, T., Nagahata, Y., Matsuura, T., Hashimoto, H., Suzuki, T., Yamashita, H., Satoh, Y.*, et al.* (2013). Distinct metabolic flow enables large-scale purification of mouse and human pluripotent stem cell-derived cardiomyocytes. Cell Stem Cell *12*, 127-137.
